# Comprehensive benchmarking of tools for nanopore-based detection of DNA methylation

**DOI:** 10.1101/2024.11.09.622763

**Authors:** Onkar Kulkarni, Reuben Jacob Mathew, Rhea Jana, Lamuk Zaveri, Sreenivas Ara, Tulasi Nagabandi, Nitesh Kumar Singh, Karthik Bharadwaj Tallapaka, Divya Tej Sowpati

## Abstract

Long read sequencing technologies such as Oxford Nanopore (ONT) offer direct, simultaneous detection of DNA base modifications. The recent migration of ONT to the upgraded R10 chemistry has spurred the development of diverse methylation detection models. However, their performance and accuracy remain unclear. Here, leveraging diverse bacterial, plant, and mammalian datasets, we systematically evaluate the current landscape of tools and models for studying DNA methylation using nanopore sequencing. Our results demonstrate that the older models remain the best choice for studying CpG methylation. We note substantial improvement of newer tools in identifying 5-methylcytosine in non-CG contexts, 6-methyladenine, and 4-methylcytosine. We highlight the sensitivity of various tools to confounding methylation nearby. We also assess the computational performance of various tools, and effects of sequencing depth, methylation abundance, read quality, and basecalling mode. Our reusable pipelines and fully open access datasets provide a framework of resources to empower future benchmarking efforts. Our work thus details the strengths and limitations of the state-of-the-art methylation models and outlines practical guidelines for researchers using nanopore sequencing to study DNA modifications.

## Introduction

DNA base modifications are the addition of chemical groups to the canonical bases of DNA. Base modifications such as methylation contribute to epigenetic control of gene expression, genome stability, and genome organization^1,2^. The dynamic nature of DNA methylation has been shown to play crucial roles in development, stress, and aging^3–5^. Depending on the base and the position in the carbon ring to which the methyl group is added, there exist several distinct modified DNA bases, the most common being 5-methylcytosine (5mC - addition of methyl group to the 5th carbon of cytosine), 5-hydroxymethyl cytosine (5hmC - a hydroxyl group at the 5th carbon of cytosine, derived by oxidation of 5mC), 6-methyladenine (6mA - methyl group at the 6th carbon of adenine), and 4-methylcytosine (4mC - addition of methyl group to the 4th carbon of cytosine)^6^. 5mC, specifically in a CG dinucleotide context (CpG) is predominantly found in mammals and its presence in promoters is associated with gene silencing^7^. Though not as well studied, 5mC is also found in sequence contexts other than CpG. In mammals, tissues such as brain harbor as much as 4-5% non-CpG methylation, with several thousand sites shown to have close to 100% methylation^8^. In plants, non-CpG methylation is important for development, stress response, and transposon silencing^9,10^. 6mA and 4mC are more commonly found in prokaryotes, where they mainly contribute to genome protection via restriction modification systems and methylation-mediated mismatch repair^11,12^. However, recent studies report the presence of 6mA and 4mC with crucial regulatory roles in eukaryotes as well^13–15^.

Traditionally, 5mC has been studied using conversion-based methods such as treatment with sodium bisulfite followed by high-throughput short-read sequencing^16^. In addition to being damaging to DNA, bisulfite sequencing is prone to amplification biases as well as difficulty in mapping 5mC in challenging genomic regions^17,18^. Further, other marks such as 6mA and 4mC are not amenable to bisulfite sequencing; their genome-wide analyses have relied on antibody-based methods which lack nucleotide level resolution^19^. Third-generation, single molecule long read sequencing methods such as Pacific Biosciences (PacBio) and Oxford Nanopore Technologies (ONT) offer simultaneous detection of DNA base modifications from sequencing data. In PacBio sequencing, variation in polymerase kinetics such as pulse width and interpulse duration (IPD) enables identification of modified bases^20^.

Nanopore sequencing works by monitoring electrical fluctuations as a DNA strand is translocated through a biological pore^21^. The electrical signal is converted to the corresponding DNA sequence using a basecaller. Differences in the electrical signals around modified bases can be leveraged to capture the information of DNA modifications^22^. ONT have constantly upgraded their technology for better accuracy and stability. Their older flowcells (FCs) with R9 chemistry used the *Escherichia coli* Curli sigma S-dependent growth (CsgG) protein^23^ as the channel, and achieved a modal accuracy of ∼95%. The current generation FCs with R10 nanopores use a bioengineered CsgG-CsgF heterodimer^24^. In addition to being more stable, this “double-barrel” nanopore achieves a modal accuracy of >99%, owing to its longer sensing region, and improves accuracy in homopolymer stretches of up to 9nt^24^. ONT have also recently changed their sampling rate from 4kHz to 5kHz, and demonstrated improved sequencing accuracy with the higher sampling rate^25^. In parallel, there have been steady improvements to the basecallers. Their latest libtorch based tool Dorado offers significant improvements in accuracy and is optimized for utilizing GPU for speed^26^. In May 2024, ONT released their v5 models for basecalling which use transformer models as opposed to the traditional LSTM based neural network models^27^. The new models claim a raw-read modal q-score of 26, and improved accuracy for identification of DNA base modifications when tested on synthetic controls^28^. However, the performance of various nanopore models on real data has not been documented so far. In addition to the models offered by ONT, few other tools support identification of 5mC in CpG context from R10.4 data: DeepBAM, DeepMod2, f5C, and RockFish^29–32^. A recent tool called DeepPlant offers models to detect 5mC in CHG and CHH contexts as well, specifically trained and geared towards plant genomes^33^.

Previous efforts have benchmarked the accuracy of base modification detection from nanopore data, but with limited scope. Earlier work by Yuen et al and Liu et al was restricted to CpG methylation using the older R9.4.1 chemistry^34,35^. Subsequent studies assessed the accuracy of CpG methylation detection from human long read datasets generated on the new R10 chemistry, and compared the accuracy of 5mC detection in CpG contexts between the R9 and R10 chemistries, particularly in the context of primary human tissues^36,37^. Another study evaluated the accuracy of 5-hmC calling and the detection of single-stranded methylation using nanopore duplex sequencing^38^. Despite these important contributions, a consistent limitation across these studies is their almost exclusive focus on CpG methylation. Therefore, the efficacy of nanopore sequencing in studying 5mC in non-CpG context, 6mA, and 4mC remains unknown. Another factor contributing to this lacuna is perhaps the lack of appropriate benchmarking datasets^39^.

Here, using diverse datasets from bacteria, plants, mouse, as well as the widely studied Genome-In-A-Bottle HG002 sample, we benchmark the performance of the methylation models available for current generation nanopore data. We highlight the sensitivity of these models to neighboring methylation, and also evaluate their computational performance, effect of overall methylation abundance, sequencing depth, read quality, sampling rate, and the basecalling mode.

## Results

### Benchmarking strategy

Most existing studies have focussed on benchmarking 5-methylcytosine (5mC), more specifically in CpG dinucleotide context. We set out to perform a comprehensive evaluation of models beyond CpG methylation. The overall strategy, including the datasets generated and used for evaluation, as well as the pipelines followed, is outlined in Figure 1. To profile 5mC in both CpG and non-CpG contexts, we generated whole genome data of two plant species (*Arabidopsis thaliana, Oryza sativa japonica* (rice)), and two mouse samples (whole brain, embryonic stem cells) using nanopore sequencing (Fig 1a). We also performed enzymatic methyl-seq (EMSeq) on these samples as “ground-truth”. Use of exogenous spike-ins revealed the conversion efficiency for all EMSeq data is >99.5% (Table S1). In addition, we used in-house (4kHz) as well as public ONT data (5kHz) of the human HG002 (GM24385) sample for which whole genome bisulfite sequencing (WGBS) data is also available in the public domain.

**Figure 1:**
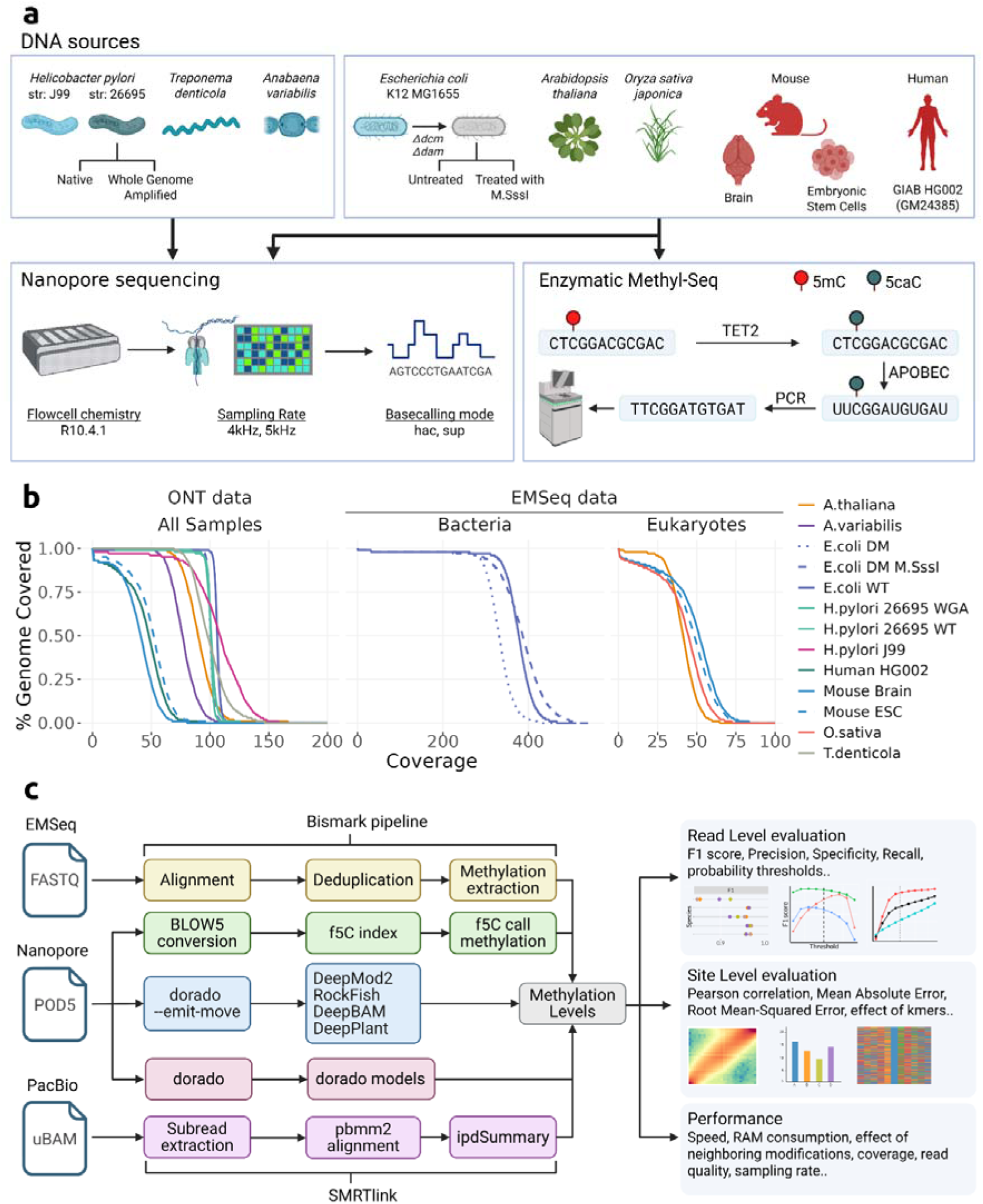
Benchmarking strategy. a) Sources of DNA and whole genome sequencing workflow used in this study. Genomic DNA from bacterial/plant/mammalian samples was sequenced on the R10.4.1 flowcells. A subset of samples (right box on the top) were also subject to Enzymatic Methyl-Seq (EMSeq) as ground truth for 5-methylcytosine (5mC). b) Line plot depicting the coverage distribution of all ONT data (left), EMSeq data for bacterial samples (middle) and EMSeq data for other samples (right). c) Data analysis pipelines. EMSeq data was processed using the Bismark pipeline. Raw nanopore data in POD5 format were either converted to BLOW5 (for f5C) or processed using dorado, including appropriate flags, to enable downstream processing by various tools. PacBio data from the public domain was processed using SMRTLink tools to derive sites harboring 6-methyladenine (6mA) and 4-methylcytosine (4mC). The per-site methylation levels derived via each pipeline were compared to evaluate various tools as per the metrics mentioned on the right. This figure was created with the help of BioRender.com.

For assessing 6mA and 4mC, we relied on bacterial datasets. Bacterial methylation systems are highly-motif driven - methylation is found in specific sequence contexts, and often at close to 100% methylation levels^40–42^. We generated nanopore data from five bacterial species: *Escherichia coli* K12 MG1655, *Helicobacter pylori* str 26695, *H. pylori* str J99, *Anabaena variabilis* ATCC 27893, and *Treponema denticola* ATCC 35405. In addition to the wild type (WT) DNA, we generated data from an *E.coli* mutant (DM), where both the major methylases, *dam* and *dcm*, were knocked out and shown in our previous study to be devoid of 5mC and 6mA in CmCWGG and GmATC contexts respectively^43^. Further, we enzymatically treated the DM gDNA with M.SssI to introduce 5mC in CpG context. We confirmed the methylation status of these *E.coli* samples using EMSeq (Fig S1), and have used this data as ground truth when assessing 5mC. For *H.pylori*, we sequenced both the native (WT) as well as whole-genome amplified (WGA) versions of the DNA. For the other species, only native (WT) DNA was sequenced. For three bacterial species - *E.coli, H.pylori* J99*, T.denticola,* we used the public PacBio data (see Data Availability) as ground truth for 6mA and 4mC. For the other two species, we used motif information from REBASE^44^, which is also derived from PacBio data. We achieved an average coverage of >100x for all bacterial samples, *A.thaliana,* and rice, and ∼50x for mouse and human samples (Fig 1b, Table S2). The median read quality for all datasets except *H.pylori 26695* was >20 (Fig S2a,b). The datasets used in this study and the expected methylation contexts in various bacterial species are summarized in Tables S2 and S3. Taken together, our datasets cover 100% of 9-mers and 97.29% of 11-mers centred for 5mC (Table S4), and 99.9% of 9-mers centred for A’s (bacterial datasets only), used for profiling 6mA models. The overall abundance of various marks ranges from ∼0.5% to ∼98% (Table S5).

Most of the existing tools for methylation detection from ONT data work on the R9 chemistry, and have been benchmarked before. As these flowcells were phased out, we focused our analysis on tools which support methylation calling for the current (R10) FC chemistry. For 5mC, these are DeepBAM, DeepMod2, DeepPlant, f5C, RockFish, and the models provided by ONT via Dorado. For Dorado models, we benchmarked both the BiLSTM based v4 models, as well as the new transformer architecture based v5 models, in both high- (hac) and super-accuracy (sup) modes. Except for Dorado and DeepPlant, other tools support 5mC calling only for CpG contexts. Dorado offers CG specific models (suffix 5mCG), as well as “all-context” models (suffix 5mC) that cover both CpG and non-CG sites. When assessing performance at CpG sites, we used the Dorado 5mCG models for comparison with other CpG tools. With the exception of f5C, other tools require the “move table” output of Dorado for identifying methylated bases, for which we used the v5 sup model (Fig 1c). For comparison of accuracy, we used the sup version for all Dorado models, as they consistently outperformed their hac counterparts (Fig S3a). Similarly, we used the Transformer model of DeepMod2 as it was marginally but consistently better than their BiLSTM model (Fig S3b). For 6mA and 4mC, currently no tools other than those offered by ONT exist. Hence, our analysis is limited to comparison of different versions of the Dorado models. The models used for each analysis across the tools and datasets, and their default probability thresholds are listed in Table S6.

### Read level evaluation

We first evaluated the read-level accuracy metrics of various tools using reads from locations which were reported to be fully unmethylated or methylated based on ground truth. For CpG methylation, our analysis spanned six diverse datasets, with ground truth established from corresponding EMSeq data (WGBS for HG002). The number of sites evaluated for each dataset, and the corresponding accuracy metrics (F1 score, Precision, Recall, and Specificity) are outlined in Table S7. RockFish and Dorado v4r1 were among the tools with highest F1 scores across species, closely followed by Dorado v5r1 and v5r3, f5C, and DeepMod2 (Fig 2a, Fig S4). Most tools had high precision and sensitivity, and the differences in F1 scores could be attributed to the varied recall values. In contrast, the poor performance of DeepPlant was due to low precision, and its plant-specific training bias was evident from its high recall only on plant datasets (Fig S4). DeepBAM showed consistently low F1 scores across the datasets tested (Fig S4). All tools had highest F1 scores on the human HG002 data, possibly reflecting the widespread use of this dataset in tool validations. The scores of all tools were lower on *A.thaliana* and mESC data, but the order of best performing tools was consistent. We then assessed the performance of these tools on non-CpG methylation using the plant and mouse data. We limited our analysis to tools that support non-CpG calling - DeepPlant, and various “all-context” models of Dorado. All Dorado models showed high accuracy across the datasets, with the v4r1 model being slightly better than the rest (Fig 2b). The species-specific trend of DeepPlant was particularly prominent here: it performed on par with the Dorado models on both the plant data, but showed an extremely low F1 score on the mouse datasets, attributable to poor recall value (Fig 2b, Table S7).

**Figure 2:**
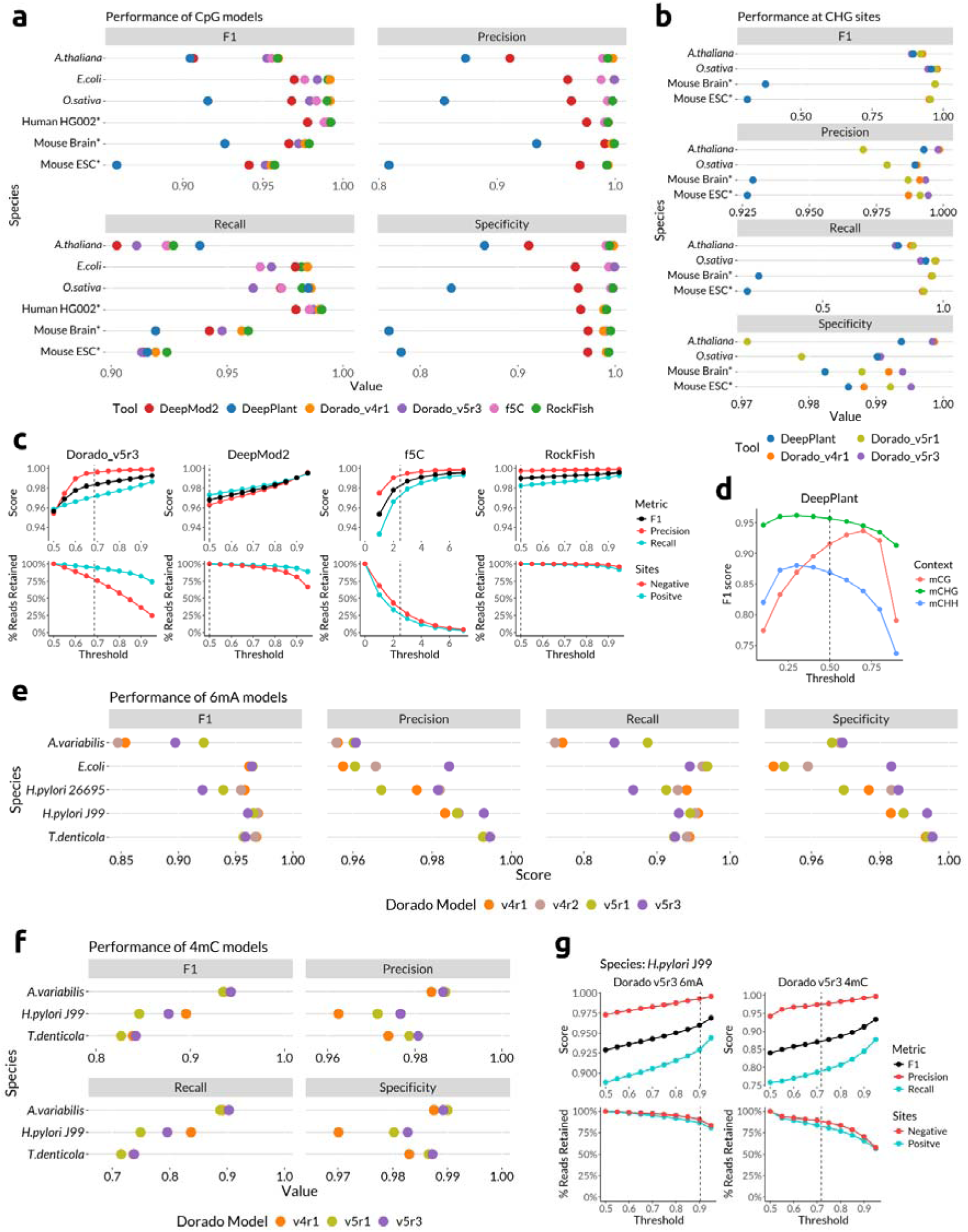
Read level evaluation of various tools. a) Performance metrics of tools on CpG sites. F1, Precision, Recall, and Specificity are represented as dots on a number line (x-axis) colored by tools. Each line represents a dataset. The asterisk next to Human HG002, mouse brain and ESC indicates that the calculations are done using data of only chromosome 1. b) Performance of DeepPlant and various Dorado models in identifying non-CpG methylation. Only 4 datasets were used, where there was abundant non-CpG methylation. c) Effect of probability thresholds on improving performance of tools, shown on O.sativa data. For each tool, top panels show the improvement of metrics as threshold is changed, whereas the bottom panels show the corresponding data loss (reads falling outside the acceptable threshold). Negative and Positive lines in the bottom panels indicate the data loss from sites with 0% or 100% methylation respectively, and contribute to improvement in Precision and Recall in the top panel respectively. Vertical dashed lines indicate the default thresholds used by the tool. d) Effect of probability thresholds (x-axis) on F1 scores (y-axis) of DeepPlant for CG, CHG, and CHH contexts. The vertical dashed line indicates the default threshold (0.5) used by DeepPlant. e) F1, precision, recall, and specificity of various Dorado 6mA models tested on 5 bacterial species, represented as colored dots on a number line. f) Performance of Dorado 4mC models tested on 3 bacterial species, represented on a number line as colored dots. g) Top panel: Effect of probability thresholds on F1, precision, and recall of 6mA (left) and 4mC (right) models (v5r3) of Dorado, tested on data of H.pylori str J99. Bottom panels indicate corresponding data loss. Vertical dashed line is the default threshold used by Modkit.

Most tools use a probability threshold to classify a base as modified or canonical. The tradeoff of choosing a stringent threshold is data loss: reads that fall outside the acceptable threshold are discarded. We sought to understand whether the default thresholds used by the tools are optimal (see Methods for details). Figure 2c summarizes the effect of increasing the thresholds on performance metrics and data loss on the *O.sativa* dataset (Table S8 for all datasets). As expected, the F1, precision, and recall values improve with increasing thresholds in all cases. The default threshold of RockFish was too lenient. Of note, even at the most stringent threshold we tested (0.95; max possible: 1), there was <10% data loss, and the F1 score improved from 0.989 to 0.996 (Table S8). Contrarily, the default threshold of f5C (abs(log-likelihood ratio) >2.5), which was possibly carried over from the original Nanopolish settings, resulted in >70% data loss. For Dorado v5r3, the improvement in Precision plateaued earlier than that of Recall, and the data loss at negative sites was faster than what is observed at positive sites. Hence, Dorado will benefit from a moderate threshold for calling canonical CpG and a stringent threshold for mCpG. Similarly, using a threshold of ∼0.9 improved the F1 score of DeepMod2 from 0.967 to 0.99 while losing <20% data. DeepPlant, unlike other tools, uses a single threshold that splits all the data into unmethylated or methylated bins, and therefore suffers no data loss. Instead, it’s a tradeoff between precision and recall. Our results show that on the data of *O.sativa*, their default threshold of 0.5 was not optimal for any of their three models; their CG model showed the highest F1 score at a threshold of 0.7, while their CHG and CHH models performed best at a threshold of 0.2-0.3 (Fig 2d). These optimal thresholds were consistent for *Arabidopsis* data (Fig S5). However, such a clear optimal threshold was not seen for CHG and CHH models on mouse datasets, further corroborating the plant-specific nature of DeepPlant (Fig S5).

Finally, we assessed the 6mA and 4mC models of Dorado using data of five bacterial species. All 6mA models were nearly identical in performance (F1 >0.95), with the v4 models being slightly better than the v5 models, except in *A.variabilis*, where v5r1 clearly outperformed the others (Fig 2e, Table S7). The deteriorated performance of v4 models in *A.variabilis* was due to presence of neighboring 5mC, as discussed more in the section detailing the effect of neighboring modifications. This indicates that while the v5 models are slightly inferior, they are more robust to confounding modified bases nearby. Contrarily, the F1 scores for the 4mC models was lower (0.8-0.9), indicating a scope for improved models in the future (Fig 2f, Table S7). We used the data from *H.pylori* str.J99 to study the effect of Modkit thresholds on accuracy of 6mA and 4mC calling (Table S8). The default threshold of Modkit for 6mA was already high (0.9) and appeared to be almost optimal, with further increase in stringency leading to slight improvement in the accuracy metrics (F1: 0.96 to 0.97), and an extra ∼7% data loss (Fig 2g left). On the other hand, increasing the threshold for 4mC from the default (0.72) to 0.8 resulted in significant data loss (∼20% more) with marginal improvement in F1 scores (0.88 to 0.9; Fig 2g right). Our results therefore suggest that the default thresholds of Modkit are largely optimal for 6mA and 4mC.

### Site level evaluation

We next evaluated the concordance of per-site methylation levels, which are continuous in nature, particularly for 5mC in eukaryotic species. For this, we calculated the overall correlation with ground truth data using Pearson Correlation Coefficient (higher is better), and the deviation from the expected levels using Mean Absolute Error (MAE; equal weight to deviation across the spectrum; lower is better) and Root Mean-Squared Error (RMSE; higher penalty for more extreme deviations; lower is better).

On *O.sativa* data at CpG sites, the Dorado_v4r1_5mCG model demonstrated the best performance, achieving the highest correlation and lowest MAE and RMSE. It was closely followed by Dorado v5r1 and RockFish. Interestingly, this ranking was inverse to the number of sites each tool profiled: RockFish profiled the most CpG locations (26.9M), followed by the v5r1 (26.3M) and v4r1 (25.8M) Dorado models (Fig 3a). The performance of other tools was variable. f5C achieved high concordance but at the cost of profiling a significantly smaller number (17.6M) of CpG sites (Fig 3a, Table S9). Consistent with our read-level evaluations, DeepBAM showed the lowest concordance due to a high volume of false negatives (FNs), while DeepPlant also performed poorly due to an abundance of false positives (FPs; Fig 3a, Fig S6-7). The performance trend was consistent across the datasets tested, with the exception of f5C, which marginally outperformed other tools on mouse datasets but could profile only ∼10% of the locations captured by Dorado models and RockFish (Fig 3b, Table S9). DeepMod2 performed well on human and mouse datasets, but showed many FPs on the plant data. We also plotted the average CpG methylation around transcription start sites of chromosome 1 (TSS, n = 3,386) in mouse brain and ES cells. All the tools except DeepBAM and DeepPlant showed a characteristic dip at the TSS, closely following the trend of EMSeq data (Fig S8a). DeepPlant showed higher FPs close to the start site, while DeepBAM showed FNs for all the sites profiled around the TSS. Surprisingly, the performance of the newer Dorado v5r3 models, including both the dedicated 5mCG and the “all-context” 5mC versions, were significantly inferior to their v4r1 predecessors, primarily due to a high rate of false negatives (Fig S8b). This suggests that for CpG methylation analysis, the older Dorado_v4r1_5mCG model and RockFish are the most reliable choices. To investigate the source of these errors, we analyzed k-mer-specific biases of these tools. We found that Dorado models tended to underpredict methylation (FNs) at CpG sites flanked by thymines and overpredict (FPs) at CpGs neighboring guanines. While this FP bias appears to be partially corrected in the v5r3 model, the FN bias persists (Fig S9). DeepMod2 also exhibited strong biases, with FPs at TCCG sites and FNs at TCGA/ACGG sites. In contrast, f5C and DeepBAM showed no discernible sequence bias, whereas RockFish and DeepPlant tended to undercall methylation at CpGs adjacent to thymines and adenines, respectively (Fig S9).

**Figure 3:**
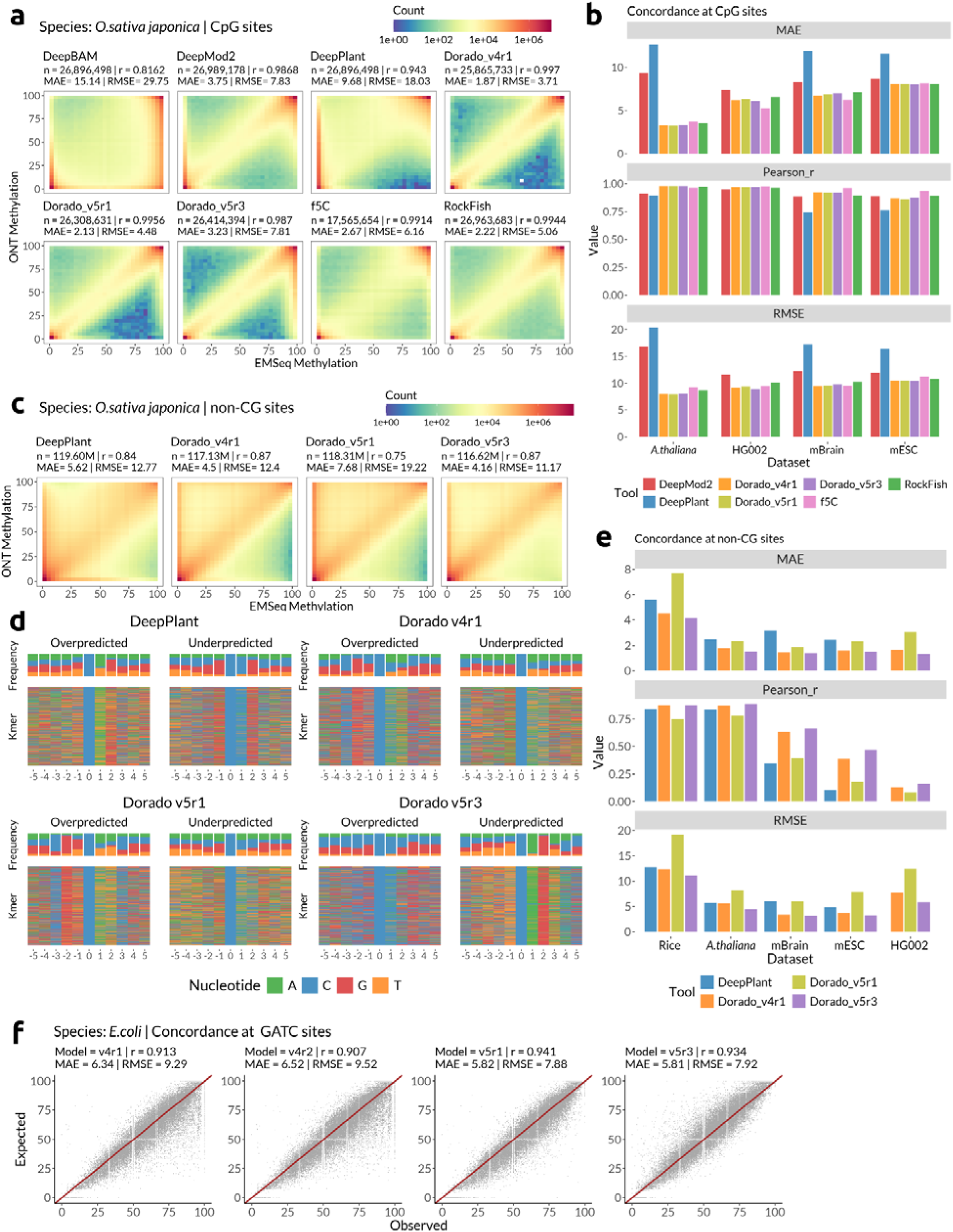
Site-level evaluation of methylation calling tools. r = Pearson correlation coefficient; MAE = Mean Absolute Error; RMSE = Root Mean-Squared Error. a) Correlation heatmap of CpG sites from rice data for each tool compared to EMSeq data (ground truth). n indicates the number of CpG sites which were profiled by both the tool and EMSeq, and covered by >=20 reads. b) MAE, Pearson r, and RMSE values of various tools at CpG locations from 4 different datasets: Arabidopsis thaliana, human HG002 GIAB cell line, mouse brain, and mouse ESC (embryonic stem cells). c) Correlation heatmaps of non-CG sites from rice data for DeepPlant and 3 different versions of Dorado all-context 5mC models, compared to EMSeq data as ground truth. n indicates locations which had at least 20x coverage reported by both EMSeq and the respective tool. d) K-mers that contribute to most disagreement with ground-truth data for each tool in O.sativa. Only non-CpG sites were chosen. 5000 most disagreeing locations were plotted in each case; Overpredicted - the nanopore tool calls more methylation than ground-truth, Underpredicted - nanopore tool calls less methylation compared to ground-truth. e) MAE, Pearson r, and RMSE values of DeepPlant and Dorado all-context 5mC models on 5 different datasets, profiled for non-CpG sites; Rice (Oryza sativa japonica), Arabidopsis thaliana, mBrain - Mouse whole brain, and mESC - mouse embryonic stem cells, HG002 - human HG002 GIAB cell line. The datasets are arranged in decreasing order of non CpG methylation abundance (Table S5). f) Concordance of various Dorado 6mA models calculated on GATC sites (n = >36k) from a synthetic E.coli dataset derived from mixing known proportions of reads from WT and DM data. The red line indicates the median diagonal of perfect correlation.

Next, we assessed the performance of DeepPlant and the Dorado 5mC models at non-CpG sites. On rice data, all tools showed lower concordance than they did at CpG sites (Fig 3c), primarily due to an increase in false positives (FPs). Overall, Dorado_v5r3 performed the best when profiling all non-CG sites together. To dissect this further, we stratified non-CG sites into CHG and CHH contexts across all datasets. On plant data, Dorado_v5r3 remained the top performer for both contexts, followed by Dorado_v4r1 and DeepPlant (Fig 3e, Fig S10). While correlation at CHG sites was high for almost all tools (r > 0.9), it was significantly lower on mouse and human datasets, which have lower non-CG methylation. Despite this low correlation, the MAE and RMSE values in mammalian data were comparable to those in plants (Table S9). This was due to the FP calls by all the models. The prominence of FP calls, and therefore the drop in correlation, was inversely proportional to the abundance of non-CG methylation (Table S5, Fig S11, S12a), a trend that was also seen on CHH sites of rice, which are less methylated compared to CHG sites (Fig S10a). However, we found these FPs could be substantially reduced by applying a more stringent modbase probability threshold; using a stringent modbase threshold of 0.975 on mESC data, while leaving the canonical C threshold at the default of 0.79, we observed a significant reduction in FPs while not losing true positives (Fig S12b). We next sought to understand which kmers contributed the most to disparity in methylation calls. Dorado v4r1 and v5r1 models overpredicted methylation at Cs with preceding Gs and succeeding As/Ts (Fig 3d). This bias appears to be rectified in the v5r3 model, but it instead showed a strong FN bias in CAG contexts. Contrarily, DeepPlant overpredicted methylation at CAG/CTG sites, and called false negatives at C’s surrounded by CGs. In summary, our site-level analysis corroborates our read-level findings for CpG methylation. For non-CpG methylation, Dorado_v5r3 is the top performer, but we recommend using stringent calling thresholds, particularly for genomes or sequence contexts with low modification levels.

Unlike eukaryotic genomes which show a continuum of 5mC levels in various contexts, bacterial methylation systems are usually binary in nature^41^. Hence, we could not directly use our data to calculate concordance metrics such as correlation. To achieve this, we created a “mixed” dataset of *E.coli*, where reads from WT and DM were combined in known proportions to achieve a continuum of GATC methylation levels (see Methods). On this data, the v5r1 model showed the highest Pearson r, and the lowest RMSE (Fig 3f). This is in line with our earlier observation that the v5r1 model was the most consistent across datasets. We then analyzed the performance of these models at known methylated contexts across our bacterial data (Table S3). Overall, the performance of the models was consistent across motifs, with occasional exceptions such as the CTAmAT motif in *T.denticola* for 6mA (Fig S13), and mCCNNGG motif in *H.pylori* J99 for 4mC (Fig S14). In general, the v5r1_6mA and v5r3_4mC models showed the tightest distributions.

### Effect of neighboring modifications

Approximately 8-10 nt of DNA contribute to the fluctuation of electrical signals at a time due to the longer sensing region of the current generation nanopores^24^. Hence, a modified base present in the vicinity of the target base can influence the accuracy of methylation calls^22^. To check the extent of this influence, we profiled the performance of tools in identifying methylated C’s and methylated A’s (mC and mA hereafter) neighboring known methylated contexts. First, we profiled the impact of nearby CpG methylation on identifying methylated and unmethylated CpGs (mCG and umCG). Unsurprisingly, when both CGs in a CGCG context are either methylated or unmethylated, the performance of the tools was similar to their overall performance (Fig S15a). However, when profiling a umCG next to mCG, several tools picked false positives, with DeepPlant and DeepMod2 showing the strongest bias followed by f5C and RockFish (Fig 4a). All versions of both 5mCG and 5mC Dorado models were robust in this scenario (Fig S15b). While assessing the effect of mCH on umC in the immediate vicinity, we observed minimal overall bias, with Dorado_v5r1 performing the best (Fig S16a). We next tested for this bias when profiling a umCG next to mA. The bias was much lower, with tools performing only slightly worse than their performance in identifying umCG next to umA (Fig 4b, left and middle). Contrarily, f5C failed to call mCGs next to mA (Fig 4b, right), perhaps because it groups nearby CGs and attributes a combined value to them, in line with the original nanopolish algorithm^22,31^.

**Figure 4:**
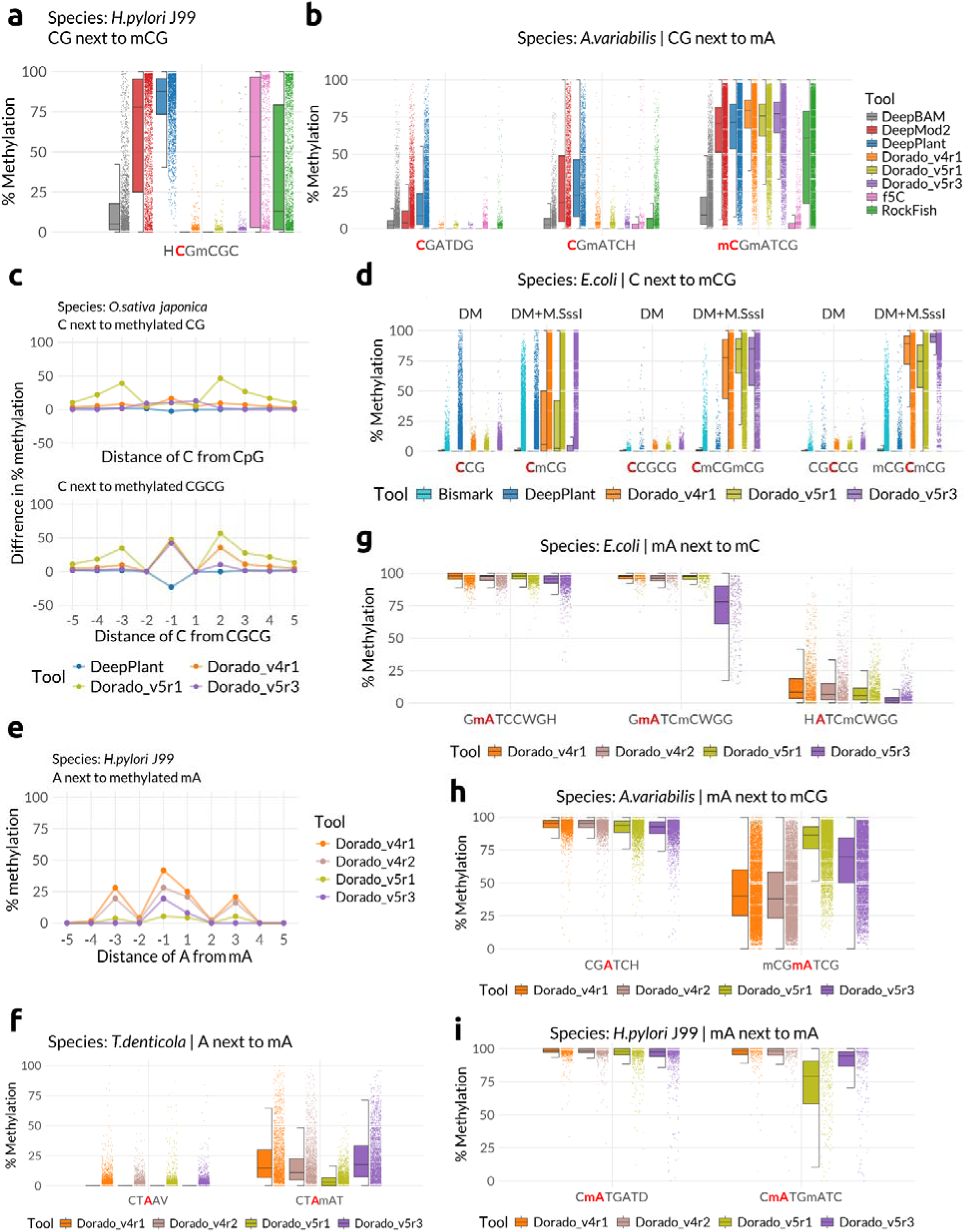
Effect of neighboring methylation on accuracy of various methylation calling tools. In all plots, the profiled base is highlighted in red font. The methylation status of the bases is indicated with a preceding “m”. In jitter-box plots, the jitter plot (right half) displays the %methylation of each evaluated site as a dot, whereas the boxplot component (left half) summarises the distribution of data with quartile ranges and median value. a) Methylation levels of an unmethylated CG adjacent to a methylated CpG (mCG) in H. pylori J99, showing false positive 5mC calls by several tools. b) Methylation levels of CG next to methylated or unmethylated A in A.variabilis, demonstrating mA-induced false positive and false negative CG calls. c) The difference in methylation calls for a cytosine as a function of its distance from a single (top) or double (bottom) methylated CpG in O.sativa. Negative values indicate underestimation of methylation near an mCG. EMSeq data is used as ground-truth. Values plotted are median disagreement levels. d) Performance of various tools on non-CG methylation in E.coli with (DM + M.SssI) and without (DM) neighboring CpG methylation, highlighting false positive calls induced due to mCGs. e) False positive methylation calls of adenines as a function of its distance from a methylated adenine (mA) in H.pylori J99. Values plotted are median methylation levels reported by various tools. f) Performance of various models in identification of unmethylated adenine next to an unmethylated (CTAAV) or methylated (CTAmAT) adenine, in T.denticola. g) Reported levels of methylated/unmethylated adenine in E.coli when located next to methylated or unmethylated cytosine. h) Methylation levels of an adenine reported by various tools when located next to an unmethylated (CGATCH) or methylated (mCGmATCG) cytosine. i) False negative calls of methylated adenine when present in the vicinity of another methylated adenine, observed in H.pylori J99 data.

We extended this analysis to profiling mC and unmethylated Cs (umC) neighboring various modifications. Dorado models called many FPs when umC was in the immediate vicinity of mCG (Fig 4c, top). This bias was exacerbated by the presence of two mCGs, and when the umC was present 3nt away from mCG (Fig 4c, bottom, Fig 4d). Out of the tested models, the v5r3 model was the most robust. DeepPlant did not suffer from this issue. Conversely, DeepPlant underpredicted methylation levels of C next to an mCG (Fig S17). This FN bias worsens when the C is adjacent to two mCGs (Fig S19). This explains our earlier observation that DeepPlant has more FNs compared to Dorado models, which had an FP bias (Fig 3c). When calling umC next to mA, Dorado models were the most accurate, with least FPs (Fig S16b,S20c). Similarly, mA did not affect nearby mC calling but DeepPlant showed FNs irrespective of the methylation status of the nearby adenine (Fig S16d).

Next, we analysed the effect of neighbouring modifications on accurately calling adenine methylation. We assessed umA at different base positions from mCG sites. Dorado models except v5r3 called many FPs, particularly for umA that is 3nt or further away from mCG (Fig S20a). Dorado v5r1 model performance was worse than other Dorado models. Conversely, the effect of mCG on mA was the least on Dorado_v5r1_6mA. Dorado v4 models showed more FNs than those of v5 models (Fig 4h). While the Dorado v5r3 model produced the fewest false positives (FPs) when assessing the effect of mCH on umA, it also resulted in the highest number of false negatives (FNs) when evaluating the effect of mCH on mA. (Fig 4g, Fig S16c). Further, we assessed performance of Dorado models in calling adenine neighboring mA. All Dorado models called false positives in calling umA next to mA, with Dorado_v5r1 being the least affected (Fig 4e, Fig 4f, Fig S20b). Instead, the v5r1 model showed abundant FNs compared to all other Dorado models when calling mA near mA (Fig 4i). In summary, our results highlight a critical pitfall when studying 5mC in non-CG contexts and 6mA in the vicinity of other modifications. Depending on the context, modification, and distance, the models suffer from both false positives and false negatives. At present, the v5r3 models appear to be the most reliable for non-CG methylation and v5r1 for identifying 6mA.

### Performance of v5.2 models

At the end of May 2025, ONT released a stable version of Dorado (v1.0) along with their new v5.2r1 base modification models, touting significant improvements in their false positive and false negative rates. While the v5.2r1 model performed similarly to v5r3 for CpG methylation, it failed to identify methylation in CCWGG on WT data of *E.coli* (Fig S21a). This issue was fixed by ONT in their subsequent release of v5.2r2 models for 5-methylcytosine^45^. Hence, we assessed the performance of v5.2r2 models for 5mC, both in CpG and non-CG contexts, and the v5.2r1 models for 6mA and 4mC.

At the read level, the v5.2r2 models, especially the all-context C model in non-CG contexts, showed superior precision and specificity compared to the older Dorado models (S21b). However, the overall F1 scores were comparable to that of older models, owing to poor recall values (Fig 5a, S21b). This result was particularly prominent for the 5mCG model. A similar observation was also observed for v5.2r1 6mA and 4mC models (Fig 5b, S22a). However, they profiled fewer reads compared to the older versions (Table S7). We further extended our analysis to site-level concordance. Surprisingly, the v5.2r2 5mCG model underperformed compared to the older models, mainly due to false negatives (Fig 5c). Profiling the kmers that contributed to this discordance revealed that the model had a propensity to call false negatives when the target CpG was surrounded by other CG residues (Fig 5d). However, the all-context C model showed marked improvement, with a substantial reduction in false positive calls (Fig 5e). Kmer analysis identified that the source of remaining false positives was adjacent G’s/T’s while the model underpredicted calls in CAG context (Fig 5f), consistent with what was observed in v5r3 models (Fig 3d).

**Figure 5:**
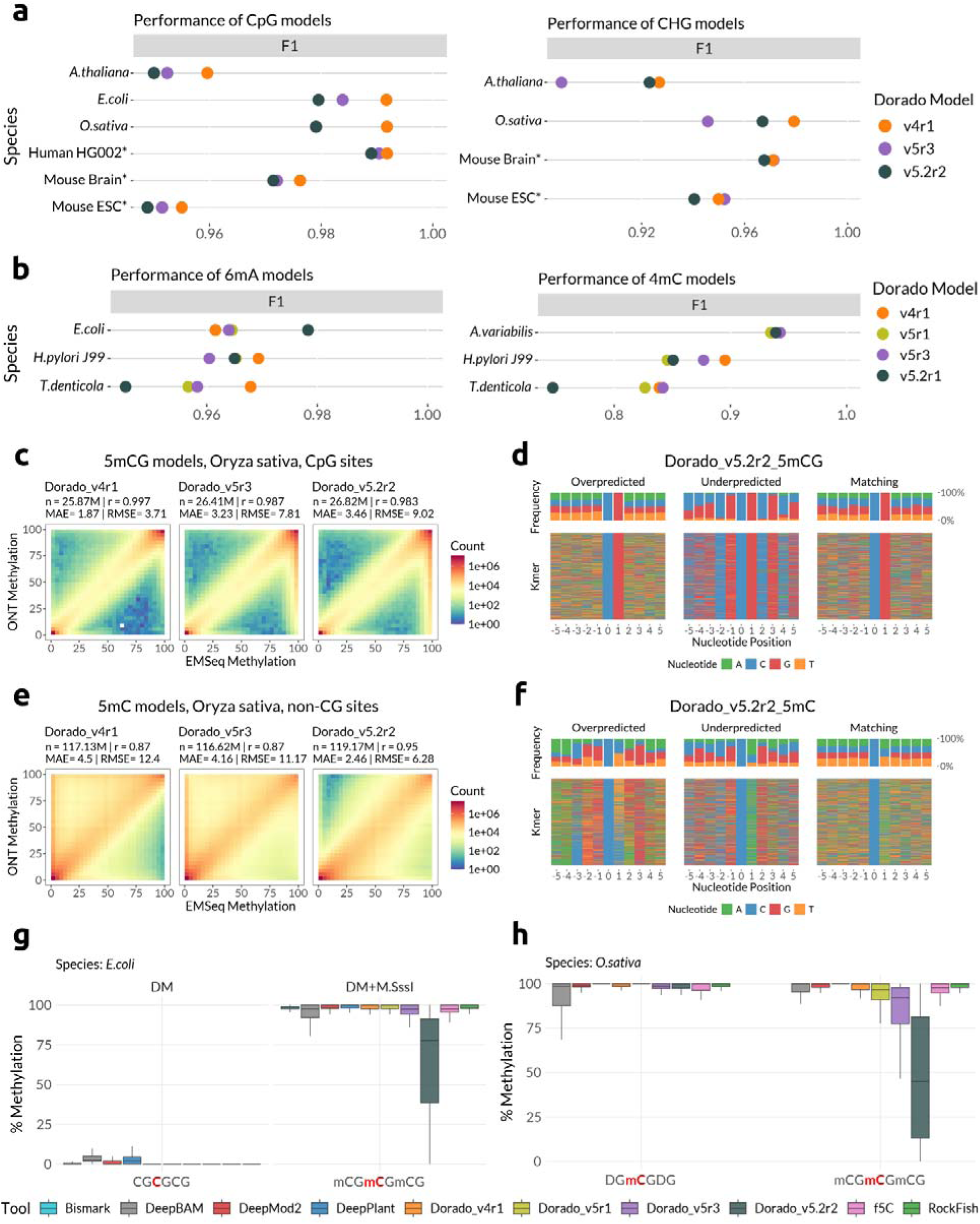
Analysis of the new v5.2 Dorado models. a) Read level evaluation of v5.2r2 CpG (left) and all-context C (right) models in comparison to v4r1 and v5r3 models. F1 scores are depicted as colored dot on a number line (x-axis). Each line represents a dataset. b) F1 scores of v5.2r1 6mA (left) and 4mC (right) models, compared to older dorado models. c, e) Site level evaluation of v5.2r2 5mCG (c) and 5mC (e) models on rice data, visualized as correlation heatmaps for each tool compared to EMSeq data (ground truth). n indicates the number of CpG/CHG sites which were profiled by both the tool and EMSeq, and covered by >=20 reads. d, f) K-mers that contribute to most disagreement with ground-truth data for each tool in O.sativa. d: CpG sites; f: non CpG sites. 5000 most disagreeing locations were plotted in each case; Overpredicted - the nanopore tool calls more methylation than ground-truth, Underpredicted - nanopore tool calls less methylation compared to ground-truth. g) Effect of neighboring CpG methylation on false negative rate of v5.2r2 5mCG models, evaluated on E.coli data. h) False negative rate of v5.2r2 5mCG model due to neighboring CpG methylation, as evaluated on rice data. Only those CpG sites which are 100% methylated as per EMSeq data were considered.

We next assessed if the v5.2 models displayed the same sensitivity to nearby confounding methylation that was observed in earlier models. The v5.2r1 6mA model displayed a vast improvement in false positives neighboring other methylated bases (Fig S22b). A similar improvement was also seen for false positives in many contexts when we used the v5.2r2 all context 5mC model (Fig S23), indicating that the new v5.2 models offer better resolution into single nucleotide methylation trends compared to earlier models. In contrast, the v5.2r2 5mCG model showed poor performance in identifying methylated CGs neighboring other mCGs (Fig 5g, h). This result explains the elevated false negative rate that we observed in our site-level evaluations, as well as the kmer trend seen at underpredicted loci (Fig 5c, d). Hence, the older v4r1 5mCG model remains the best choice when the analysis is restricted to CpG sites alone.

### Other considerations

#### Computational performance

The speed and computational resources required to run various tools have a significant impact on their practical utility. We measured the throughput of the tools benchmarked here (expressed as megabases processed per second) as well as their RAM consumption, evaluated on various bacterial datasets. The computational performance of the tested tools varied significantly across both processing speed and memory usage (Fig 6a, 6b). The Dorado models offered the highest throughput while consistently requiring a moderate 12-14 GB of RAM. Specifically, older versions like v4r1 were fastest for 5mCG detection, whereas the newer v5r3 and v5.2 models showed the highest speeds for 6mA and 4mC analysis. The hac versions ran 2-3 times faster than their sup counterparts. In contrast, RockFish, and DeepMod2 were more memory-efficient, using less than 10 GB of RAM, but operated at much lower processing speeds. These speeds also exclude the time taken to generate the Dorado move-table BAM. f5C was one of the fastest tools, beating the sup Dorado_5mCG model in both speed and memory consumption. Due to a CUDA version mismatch, we could not run RockFish using FlashAttention^46^, which is likely to improve its processing speed significantly. Some tools had high memory requirements; DeepPlant was the most resource-intensive, consuming an average of over 84 GB of RAM on bacterial datasets, followed by DeepBAM at over 31 GB (Fig 6b). Further, their performance did not scale linearly as the dataset size increased. Our initial attempts to run DeepPlant and DeepBAM on the full datasets of rice and mouse yielded no output after running for several weeks. Hence, we created an optimization pipeline (see Methods) that split the dataset into small chunks of POD5 and corresponding move-table BAM files. On these smaller pairs, DeepPlant ran at a similar speed to that of bacterial datasets. These results show a direct trade-off between processing speed, memory usage, and optimization across the evaluated tools.

**Figure 6:**
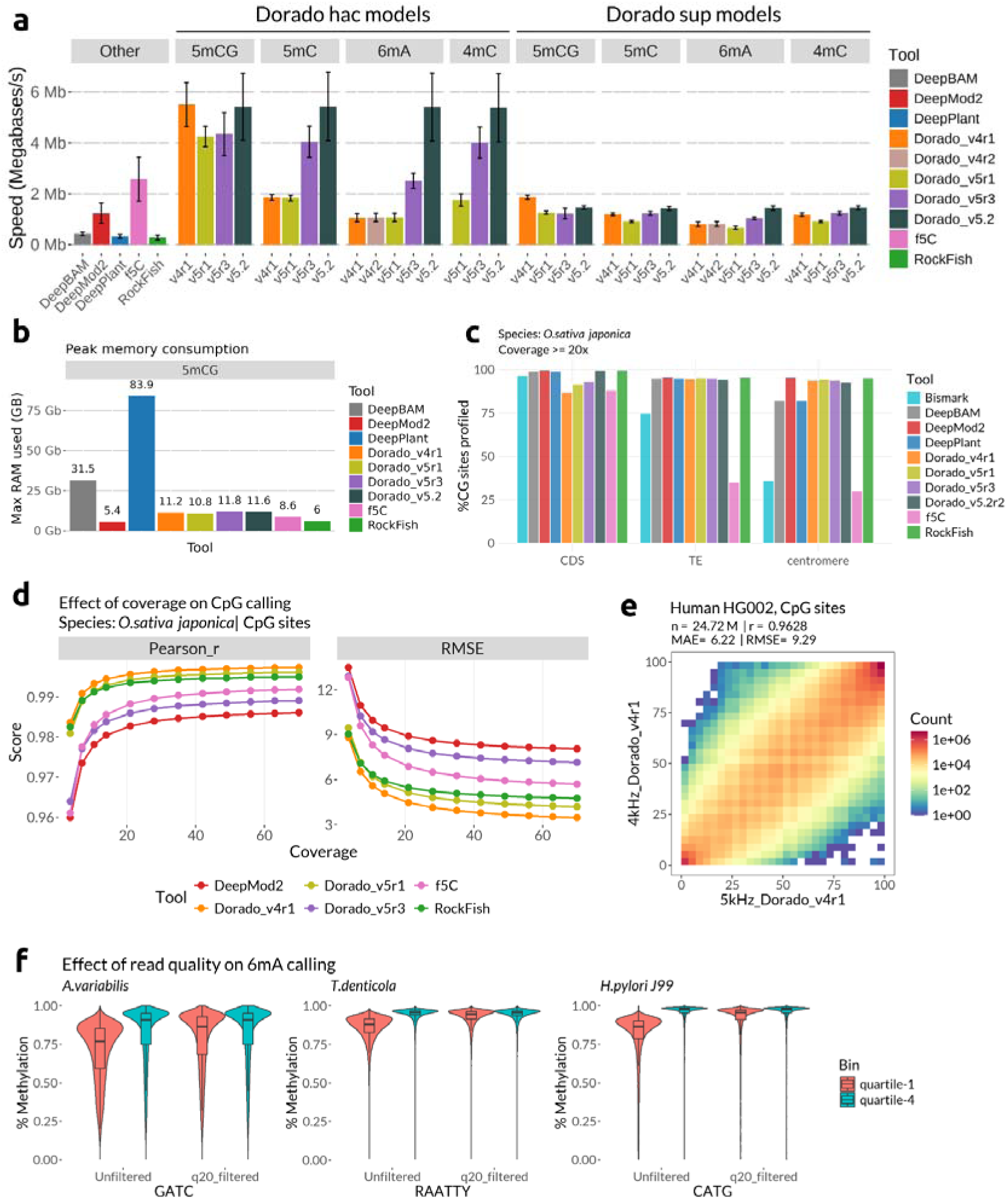
Computational performance and factors influencing the accuracy of DNA methylation callers. a) Comparison of processing speed (Megabases/s) of various models for detecting different modifications, tested on various bacterial datasets. The performance of Dorado hac (high-accuracy) and sup (super-accuracy) models are shown separately. Values reported are arithmetic mean of at least 8 measurements. Error bars represent standard deviation. b) Peak RAM consumption (GB) for variou 5mCG models and tools. Values reported are the average of >8 measurements on different bacterial datasets. c) Percentage of CpG sites profiled by different tools in distinct genomic regions (CDS: coding exons, centromeres, and TE: transposable elements) of the O.sativa japonica genome. Only sites covered by at least 20 reads were reported. d) Effect of sequencing coverage on the accuracy of CpG calling in O.sativa, measured by Pearson correlation (left) and RMSE (right) for top-performing tools. e) Concordance heatmap of CpG methylation calls for the human HG002 sample between data generated at 4kHz and 5kHz sampling rates, analyzed with the Dorado v4r1 model. f) Effect of applying a Phred quality score filter (q20) on 6mA methylation calling in three bacterial species. The plots show methylation distributions for reads from the lowest (quartile-1) and highest (quartile-4) quality bins, both before and after filtering.

#### Choosing between basecalling modes

As discussed in the computational performance section, the hac models of Dorado run significantly faster than the sup models. We assessed the accuracy differences between these two basecalling modes at read-level using several different datasets. For 5mCG (CpG) methylation, the performance of hac models is nearly identical to the sup models across all metrics, indicating the faster hac mode is sufficient for this context (Fig S24a). However, for detecting 6mA and 4mC modifications, the sup models consistently provide a clear advantage, showing higher F1 and other metrics (Fig S24b,c). This suggests that the additional computational cost of the sup mode is beneficial for achieving maximum accuracy for these non-CpG modifications. Of note, wherever we were successful in running the v5.2r1 models, their hac models were highly accurate, indicating that future models are likely to offer high speed without compromising accuracy.

#### Sequencing depth and number of profiled locations

Conversion based methods such as EMSeq and WGBS are shown to be not uniform in informing about bases across the genome - a caveat that long read sequencing is expected to solve. Indeed, we observe that long read data profiles significantly more CpGs compared to the corresponding EMSeq data, particularly when paired with tools such as DeepPlant, DeepMod2 and Rockfish, which suffer from minimal data loss (Fig 6c). This difference is further prominent when looking at challenging genomic regions such as centromeres and repeat elements (Fig 6c). This advantage also extends to Cs in non-CG contexts (Fig S25a).

To check the effect of sequencing depth on accuracy of CpG calls, we subsampled reads from the rice data from 5% to 100% of the total data, repeating this 5 times to account for subsampling biases (see Methods). We then profiled the concordance using Pearson_r and RMSE as a function of the median coverage of the data (Fig 6d). The improvement in correlation plateaued after 20x coverage for all the tools, with the best performing tools (Dorado v4r1, v5r1, and RockFIsh) reaching the plateau earlier. However, there continued to be a decrease in RMSE even at the highest coverage tested. In light of this, we suggest a median coverage of at least 20x to be suitable for routine methylation studies. Taken together with our observations on false positives and the impact of raw read quality (see below), we recommend generating additional data when studying less abundant modifications, as a significant data loss is expected due to stringent filtering criteria.

#### Effect of sampling rate

Currently, data generated on ONT machines use the new 5kHz sampling rate (the number of electrical signal data points), which was increased from the older rate of 4kHz to improve the read quality. Support for methylation calling on the 4kHz data is limited to the v4 models of Dorado and Rerio. We asked if there was a difference between the models geared for these two sampling rates. The CpG methylation calls of both the models were highly concordant on the human HG002 data (Fig 6e). The same was applicable for CHG calls too, as observed on the *E.coli data* (Fig S25b left). But the 4kHz model was significantly impacted by the neighboring methylation, both when profiling nonCG methylation (Fig S25b) and 6mA (Fig S25c). The corresponding 5kHz models were more robust, highlighting the advantages of the increased signal resolution the new sampling rate offers.

#### Impact of read quality

A major improvement with the increase in the sampling rate as well as the release of the new transformer-model architecture by ONT is the increase in the modal Phred score (q-score). Hence, we assessed if the accuracy of base modification detection is dependent on the underlying read-quality. To do this, we separately analyzed reads belonging to the lowest quality quartile (Q1) and the highest quartile (Q4) from three different bacterial datasets using the v5r3_6mA_sup model. In the unfiltered data for all three species, reads from Q1 consistently show lower and more variable methylation levels compared to reads from Q4 at known methylated contexts (Fig 6f). This discrepancy indicates that low-quality reads introduce a systematic bias, and can lead to an underestimation of the true methylation level. We then repeated this analysis after filtering for reads with a minimum Phred score of 20. After applying a q20 filter, the methylation distributions of the Q1 and Q4 bins become nearly identical (Fig 6f). This result shows that filtering low quality reads is a crucial step to remove quality-dependent biases. Hence, we recommend a minimum q-score of 20 to achieve more robust methylation calls.

## Discussion

Direct sequencing of the native, non-amplified DNA by long-read technologies such as ONT enables simultaneous detection of base modifications present on the DNA. This advantage has been extensively leveraged to study DNA methylation using nanopore sequencing, especially CpG methylation in eukaryotic genomes. However, long-read sequencing has the ability to capture all DNA modifications concurrently, as opposed to traditional conversion based methods that measure only one modification at a time. This power is contingent on the availability of good models that can accurately identify various DNA modifications. In this study, in addition to benchmarking all the CpG callers currently available for the R10 flow cell chemistry, we assess the performance of non-CpG, 6mA, and 4mC models.

Our results highlight the subtle differences in performance of different CpG methylation callers, particularly in challenging scenarios. RockFish was consistently among the top performing tools and also retained the most amount of data, even at highly stringent thresholds, highlighting its utility when dealing with low-coverage datasets. While its processing speed was low, running it with FlashAttention is likely to speed it up several fold, making it a practical choice. However, its accuracy was significantly affected by neighboring CpG methylation. While this may not be of concern when analyzing datasets such as those of mammalian genomes, where the methylation status of consecutive and nearby CpGs is often similar, caution is warranted when applying RockFish to densely methylated regions or when varied methylation of nearby CpGs is expected. The v4r1_5mCG model, which is highly accurate, works out of the box with Dorado and is optimized for GPU utilization is another top choice. This version consistently outperformed newer Dorado models, including the latest v5.2r2 version. Our finding that the hac model of v4r1 performs on par with the sup model suggests that for most studies focused on CpG methylation, particularly at population scale, the hac model is now sufficient, providing an optimal balance of speed and accuracy. DeepMod2 also performed well on human and mouse data, with minimal data loss. In contrast, both DeepBAM and DeepPlant exhibited lower overall performance for CpG sites, and the use of a stringent default threshold by f5C resulted in significant data loss. Additionally, DeepPlant overrepresented the percentage methylation at CG sites in promoters, while DeepBAM underreported it in both mouse brain and ESCs. This is consistent with our previous evaluation, where we attribute the poor performance of DeepBAM to a large number of FNs and DeepPlant due to high FPs. There is, thus, a direct consequence of the tool accuracy on further analyses profiling methylation-level changes.

For non-CpG methylation, DeepPlant was accurate on the plant data, but this did not generalize to other datasets, underscoring the bias imparted by its training data. This result emphasizes the need for diverse datasets in model development. The v5r3_5mC model of Dorado performed the best in most scenarios, but was prone to false positives in datasets with low abundance of non-CG methylation. While its FP rate is an improvement over older models, this could still impact downstream results; for example, changes in promoter methylation levels might be falsely attributed to changes at non-CG residues. Using a stringent threshold for positive calls mitigates this issue, and we recommend this approach when working with low abundance datasets. The 5mC models were substantially affected by neighboring CpG methylation, particularly when the target base is ∼3nt away. The effect was further exaggerated when multiple mCGs were present around the target base. DeepPlant was less susceptible to this but tended to underestimate methylation near mCGs and its plant-specific performance limits its utility to other data.

The impact of neighboring methylation was also seen in 6mA models to varying degrees. The v5r1 model was the most robust, but still showed several FPs when profiling As 3nt away from mAs. It also tended to undercall adenine methylation neighboring other methylated adenines. These results highlight a critical age-old challenge^22^: the electrical signal of a modified base is not generated in isolation. Enhancing the “attention” mechanism within transformer architecture models could allow them to better weigh the influence of adjacent bases, and lead to more robust and context-aware modification calling. The improvements offered by the new v5.2 models in this regard are a glimpse that the upcoming versions are headed in that direction. However, as bias still remains in some contexts, single-nucleotide trends in densely methylated regions should be interpreted with circumspection.

ONT sequencing is a rapidly evolving technology at the level of flow cell chemistry, sequencing parameters, and software. We show that the CpG calls from data generated on both the 4kHz and 5kHz sampling rates were concordant, further extending the work by Genner et al^37^ who have performed a similar comparison between the R9 and R10 chemistries. These results indicate that methylation calls of cohorts generated on different sampling rates and chemistries are comparable, and can be analyzed together. Our study has several limitations. We used EMSeq data as the ground truth, which can introduce certain biases as discussed earlier in the manuscript. In addition, we did not benchmark 5-hydroxymethyl cytosine (5hmC), another commonly studied mark that has been evaluated in some previous efforts^38^. Our analysis of 4mC was also limited to few sequence contexts due to its relatively rare nature. Furthermore, for two bacterial datasets, we relied on REBASE for methylation context information due to the lack of corresponding PacBio data.

Despite these limitations, our work makes several key contributions that advance the field beyond previous efforts. First, we demonstrate that the latest models are not universally superior and identify clear winners for specific tasks: Dorado_v4r1 and RockFish (especially for low coverage data) for CpG methylation, Dorado_v5r3 models for non-CG methylation and 4mC, and Dorado_v5r1 for 6mA. We provide practical guidelines on read quality thresholds (>q20) and optimal coverage: >20x after necessary filters. We also highlight the gaps and pitfalls of the current models, and offer mitigation strategies. Furthermore, to address the challenges encountered during our work due to speed of various tools, inconsistent support, and lack of clear instructions, we optimized several analysis scripts. These scripts, as well as our complete Snakemake and Nextflow pipelines are provided in an accompanying GitHub repository to aid others. Finally, the diverse, validated dataset created in this study represents a key resource that can serve future benchmarking efforts and provide a foundation for building more sophisticated and accurate models that can unlock the full potential of native DNA sequencing.

## Methods

### Datasets

For *Escherichia coli* str. K12 substr. MG1655, we used the wildtype (WT) strain, as well as a *dam- dcm-* double mutant (DM) which lacks canonical 5mC and 6mA (in CmCWGG and GmATC contexts respectively). This DM strain was extensively characterized in our previous work^43^. For the human data, we used the genomic DNA of HG002 GIAB cell line. Genomic DNA of *Helicobacter pylori* str J99, *Anabaena variabilis* (ATCC 27893), and *Treponema denticola* (ATCC 35405) was procured from the American Type Culture Collection. Genomic DNA from *Helicobacter pylori* str. 26695, mouse and plant samples were kind gifts from other labs in our institute (see Acknowledgements). We generated the unmethylated counterpart of *H.pylori* str. 26695 by performing whole genome amplification (WGA). For *E.coli,* mouse, and plant samples, we performed Enzymatic Methyl-seq (EMSeq) as ground truth for 5mC. The ground truth bisulfite data for HG002 was downloaded from the ONT public repository (see Data availability). PacBio data of three bacterial species was downloaded from open data resource of PacBio (see Data availability).

### Genomic DNA extraction

Overnight cultures of *E.coli* were centrifuged, washed in PBS and treated with a mixture of Lysozyme and SDS followed by Proteinase K treatment. Genomic DNA was purified using Cetyltrimethylammonium bromide (CTAB) followed by Phenol:Chloroform:Isoamyl alcohol. The gDNA was then washed with Chloroform:Isoamyl amyl alcohol and precipitated using Isopropanol. The gDNA was spooled using a glass rod, washed with ethanol and air dried. The spooled gDNA was dissolved in EB (Qiagen, Germany) and quantified using Qubit (ThermoFisher Scientific, US).

### *in vitro* DNA Methylation

For *in vitro* DNA methylation, the double mutant *E.coli* was used. The gDNA was treated with M.SssI (NEB, USA) for CpG methylation The gDNA was treated with the enzyme in a mix containing S-Adenosyl methionine (SAM) and its corresponding buffer, for 4 hours at 37°C. The samples were purified using Ampure XP beads (Beckman Coulter, USA) and quantified using Qubit (ThermoFisher Scientific, US).

### Whole Genome Amplification

Genomic DNA from *E.coli* K12 MG1655 or *H.pylori* str. 26695 was used as a template for whole genome amplification. To amplify the genetic material, the PCR Barcoding Expansion Kit (Nanopore, UK) was used according to the manufacturer’s protocol. 100 ng of genomic DNA was A-tailed using NEBNext Ultra II End Repair / dA-tailing Module (NEB, US) and purified using Ampure XP beads (Beckman Coulter, US). Barcode adapters were ligated to the sample and purified. The samples were PCR amplified for 15 cycles using compatible PCR Barcodes from the kit and LongAmp Taq 2X master mix (NEB, US). The amplified product was purified, and subsequently quantified using Qubit (ThermoFisher Scientific, US).

### Nanopore library preparation and sequencing

Nanopore libraries were prepared using Native Barcoding Kit V14 (Oxford Nanopore, UK) according to the manufacturer’s protocol. In brief, the gDNA or WGA DNA was treated with NEBNext Ultra II End Repair/dA-Tailing Module and NEBNext FFPE DNA Repair Mix (NEB, US). The samples were purified using Ampure XP beads (Beckman Coulter, US) and barcodes were ligated to the DNA fragments. Samples were purified once more and quantified using Qubit (ThermoFisher Scientific, US). Equimolar amounts of the barcoded samples were pooled and ligated to the native adapter. Post purification, the library was quantified and 50 fmol was loaded onto an R10.4.1 MinION flow cell or an R10.4.1 PromethION flow cell and sequenced.

### Enzymatic Methyl-seq library preparation

200ng of purified genomic DNA was used for each library preparation. For mouse and plant samples, three different exogenous spike-ins were used as controls. Two spike-ins were common for all - Lambda DNA as unmethylated control, and pUC19 DNA treated with M.SssI as methylated control. Both these controls were part of the NEBNext Enzymatic Methyl-seq (EMSeq) Kit (NEB, US). In addition, two amplicons were used as unmethylated controls - a 1781bp amplicon of the YejM gene from *E.coli* for the mouse samples, and a 4599bp amplicon of the human CYP21A2 gene for the plant samples. The primers used for generating these amplicons are given below.

YejM-FP: 5’ GTCCTCTAGAATGGTAACTCATCGTCAG 3’

YejM-RP: 5’ GAGCAAGCTTTCAGTTAGCGATAAAAC 3’

CYP21A2-FP: 5’ GGACACTATTGCCTGCACAGTTG 3’

CYP21A2-RP: 5’ TGTCCACCTCTTTCACCACG 3’

DNA was processed using the NEBNext Enzymatic Methyl-seq (EMSeq) Kit (NEB, US) as per the manufacturer’s protocol. DNA was fragmented using Covaris Focussed Sonicator to an average fragment size of 300 bp and DNA was treated with an end prep enzyme mix followed by adapter ligation. DNA was oxidized using a TET2 containing mix and denatured with Formamide. The denatured DNA was treated with an APOBEC mix to deaminate the Cytosines. Indexes were incorporated into the DNA fragments using PCR and purified afterwards. Post quantification, the samples were pooled at an equimolar ratio and run on an Illumina Novaseq 6000 at PE150.

### EMSeq Data Processing

EMSeq data was processed following the Bismark pipeline in the nf-core MethylSeq workflow v24.10.5. Briefly, the quality of Illumina raw reads was checked using FastQC v0.12.1. Adapters were trimmed and low quality reads were filtered using Cutadapt v4.8 with default parameters^47^. The trimmed data was mapped to the respective reference genomes using Bismark v0.24.2^48^. The reference genomes used for various species are listed in Table S11. The mapped data was deduplicated and cytosine methylation was extracted for all the cytosines using ‘deduplicate_bismark’ and ‘bismark_methylation_extractor’ respectively. Processed bisulfite data for the human HG002 cell line was downloaded from the ONT open data repository hosted on AWS (see Data Availability).

### PacBio Data Processing

PacBio CCS reads from three bacterial species (*E.coli* K12 MG1655*, H.pylori J99, T.denticola*) were downloaded from PacBio website (See Data Availability). All processing was done using SMRTlink tools v13.1^49^. BAM files of unmapped CCS reads were merged using samtools, and index using ‘pbindex’. CCS reads were converted to “pseudo”-subreads using ‘ccs-kinetics-bystrandify’. Reads were aligned to the respective reference genomes, sorted, and indexed using pbmm2. Methylated bases (6mA and 4mC) were identified using ipdSummary with the command “$SMRTlink_home/smrtcmds/bin/ipdSummary -j 50 $sample.aligned.bam --reference $sample.fa --identify m6A,m4C --minCoverage 20 --methylFraction --pvalue 0.01 --gff $sample.modbase.gff”. The GFF file contains details of the modified base, its methylation percentage (frac), and 95% confidence intervals (fracLow and fracUp). These details were used to retain only those sites which are fully methylated: fracLow=frac=fracUp=1. In parallel, ipdSummary was also run with a p-value cutoff of 1, to report all sites irrespective of their methylation status. This was used to identify sites which are fully unmethylated, by filtering for sites with IPD Ratio of <1 (no deviation of IPD from what is expected for a canonical base). These subsets of locations were used as ground truth locations for 6mA and 4mC.

### Nanopore basecalling and quality control

We primarily used the Dorado basecaller v0.9.1 for basecalling all nanopore reads in POD5 file format. Most other tools require the “move table” output of Dorado to call base modifications. For this, we basecalled the POD5 datasets using Dorado v5.0.0 super-accuracy (sup) model with the ‘--emit-moves’ flag to generate BAM files that were compatible with downstream processing using DeepMod2, DeepBAM, DeepPlant, and RockFish.

Unaligned BAM files produced by Dorado were then converted to FASTQ files using ‘samtools fastq’^50^ with the ‘-T “*”’ argument to preserve all the read tags included in the BAM file; these reads were then aligned to the appropriate reference genomes (Table S11) using Minimap2 v2.28-r1209^51^. Quality checks were performed on the aligned-bam files using NanoStat^52^ v1.6.0 and the coverage of the dataset was calculated using mosdepth^53^ v0.3.10. All bacterial datasets were downsampled to an average depth of 100x using Rasusa^54^ v2.1.0. Reads with average Phred score <10 and reads shorter than 500nt were filtered out using a custom Python script.

### Methylation Calling

#### Dorado

For modification calling, dorado was run multiple times with appropriate models. Dorado v0.9.1 was used for methylation calling using the v4 and v5 models on the 5kHz datasets. Dorado v0.5.3 was used for basecalling 4kHz data, and Dorado v1.0 was used to test the most recent v5.2 models. Both high-accuracy (hac) and super-accuracy (sup) models were tested wherever available. The output of Dorado is a “modbam” file, with read-wise methylation information encoded in ML and MM tags. Modkit v0.4.4 (v0.5.1 for v5.2 models) (https://github.com/nanoporetech/modkit) was used to derive per-site methylation information in BED format using the ‘pileup’ function. Example commands used are:

~~~
“dorado basecaller doradomodels/dna_r10.4.1_e8.2_400bps_sup@v5.0.0 \
<POD5_DIR> --reference reference.fa \
--min-qscore 10 --modified-bases-models \
doradomodels/dna_r10.4.1_e8.2_400bps_sup@v5.0.0_5mCG_5hmCG@v3 | \ samtools sort -O BAM -o $sample.bam”
“modkit pileup $sample.bam $sample.DoradoOutput.bed -t 40 --ref \ reference.fa --ignore h”
~~~

In addition to the methylation percentage, Modkit provides information about the read coverage as well as the number of supporting reads. Modkit also filters out info from reads which have a base that is different from the expected target base, either due to a genuine variation in the sample, or due to sequencing errors.

#### DeepBAM

The BAM file with move table information from Dorado v5r3 sup model was first sorted by fn tag to arrange reads by the POD5 file they belonged to, and output was then used as input to DeepBAM. DeepBAM ‘extract_and_call_mods’ was run using their 2024 pytorch models, with default parameters of kmer size (51), batch size (1024), and 40 (8 main and 4 sub threads each) worker threads, which gives read-wise methylation probabilities. These were aggregated into per-site methylation values using their aggregation script. The example commands are:

~~~
“samtools sort -t fn $sample.movetable.bam -O BAM -o
$sample.movetable.fn_sorted.bam
DeepBAM extract_and_call_mods <POD5_DIR>
$sample.movetable.fn_sorted.bam reference.fa DNA $sample.readwise.out LSTM_20240524_newfeature_script_b9_s15_epoch25_accuracy0.9742.pt 51 8 4 1024 CG 0”
“python deepbam_aggregate.py --file_path $sample.readwise.out --aggregation_output $sample.DeepBamOutput.tsv --threshold 0.5”
~~~

#### DeepMod2

DeepMod2 offers both BiLSTM and Transformer models. Their new v5.0 models were run on the move table BAM file output of Dorado v5r3 sup model with default parameters, using both BiLSTM and Transformer models. DeepMod2 outputs several files, including read-wise and site-level information. The “output.per_site” file, where the strand-wise information is available, was used for further comparison. The example command used was:

~~~
“python deepmod2 detect --bam $sample.movetable.bam --input <POD5_DIR>
--model bilstm_r10.4.1_5khz_v5.0 --file_type pod5 --threads 40 --ref reference.fa --seq_type dna --output $sample.DeepMod2_BiLSTM.out”
~~~

#### DeepPlant

DeepPlant was run in a manner similar to DeepBAM. Dorado v5r3 sup BAM file with move table info was first sorted by the the fn tag using samtools sort to arrange the reads by the POD5 file they belonged to. The re-sorted BAM file was then used as input. The CG, CHG, and CHH models of DeepPlant were run using default kmer sizes (51, 51, 13), batch size of 2048, and 40 (8 main and 4 sub threads each) worker threads. The read-wise methylation probabilities were aggregated into site-level values using the same aggregation script that was used for DeepBAM (see above). The DeepPlant command used was:

~~~
“samtools sort -t fn $sample.movetable.bam -O BAM -o
$sample.movetable.fn_sorted.bam
DeepPlant extract_and_call_mods <POD5_DIR>
$sample.movetable.fn_sorted.bam reference.fa DNA <OUTPUT_DIR> DeepPlant/model/bilstm/ 51 51 13 8 4 2048”
~~~

#### f5C

f5C v1.5 was used. Unlike other tools, f5C works directly with ONT signal data, without the need of a basecalled BAM or move table information. Raw signal in POD5 format was converted to BLOW5 format using blue-crab. The reads were indexed using ‘f5C index’, and read-wise methylation was called using ‘f5C call-methylation’ using the ‘--pore r10’ flag. The default output of f5C is non-stranded. To enable comparison with other tools, all of which report the calls by each strand, we added the strand information based on the reference nucleotide using an in-house Python script (f5C_restrand.py). This stranded read data was aggregated using ‘f5C meth-freq’ using the flag ‘-s’ to report the methylation frequency of each CpG rather than the default “grouped” output. The commands used were as follows:

~~~
“blue-crab p2s <POD5_DIR> -t 20 -o $sample.blow5” “f5c_x86_64_linux_cuda index --slow5 $sample.blow5 $sample.v5r3.fastq”
“f5c_x86_64_linux_cuda call-methylation --cuda-dev-id 1 -x hpc-high -K 2048 -B 40.0M -t 20 --slow5 $sample.blow5 -b $sample_v5r3.bam -g reference.fa -r $sample.v5r3.fastq --pore r10”
“python in_house/scripts/f5c_restrand.py -i $sample.f5c.readwise.tsv - r reference.fa -o $sample.restranded.tsv”
“f5c_x86_64_linux_cuda meth-freq -i $sample.restranded.tsv -s”
~~~

#### RockFish

RockFish built from the R10.4.1 branch was run on the move table BAM file of Dorado v5r3 sup model. RockFish inference was run using a batch size of 16384, 40 worker threads, and 1 GPU. The R10.4.1 branch of RockFish generates a three column readwise output that is different from their earlier branch and is therefore incompatible with their original data aggregation script. Hence, each read id and its positions were converted into their respective reference genome based coordinates using “extract_ref_pos” Python script provided by RockFish. This information was aggregated to derive per-site methylation levels using an optimized version of “rockfish_aggregate”. The commands used for each step were:

~~~
“rockfish inference -i <POD5_DIR> --bam_path $sample.movetable.bam --model_path rockfishmodels/rf_5kHz.ckpt -r -t 40 -b 16384 -d 1”
“python rockfish/scripts/extract_ref_pos.py --workers 40
$sample.movetable.bam reference.fa > $reference_lookup” # generates a remapped tsv
”python rockfish/scripts/rockfish_intersect.py -i oredictions.tsv -r
$reference_lookup.tsv -o $predictions_remapped.tsv”# generates a remapped tsv
“python in_house/scripts/rockfish_aggregate.py -i
$predictions_remapped.tsv-o $sample.consolidated.tsv”
~~~

All pipelines used above are provided as snakemake files in our accompanying GitHub repository (see Data availability). To account for coverage biases, we excluded all sites that were covered <20x from evaluation. Analysis was limited to the main chromosomes of various species, excluding sex chromosomes, alternate contigs, chloroplast, mitochondria, and plasmid contigs. When comparing metrics with DeepPlant for HG002, mouse Brain and mouse ESC data, we limited our evaluation to chromosome 1 of respective genomes. However, metrics for the entire genome for other tools is provided in corresponding supplementary tables. Comparison of per-site values with EMSeq data was done using custom Python and R scripts. The data was summarized and visualized using dplyr and ggplot2 in the R statistical environment (see Data Availability).

### Assessing methylation calls across genomic features

We downloaded the knownGene table for the mouse assembly GRCm39/mm39 from UCSC Table Browser in BED format and a GTF file containing gene information of *O.sativa* from NCBI (see Data availability). As we could run DeepPlant only on chromosome 1 data of mouse, we limited our Transcription Start Site (TSS) analysis to only that chromosome. For mouse datasets, the coordinates of the TSS on chromosome 1 were retrieved based on gene orientation and extended by 5 kb on either side. The methylation profiles of the entire 10 kb region for all the TSS (n = 3,386) were averaged and visualized as a line plot using R. This analysis was performed on both mouse ESC and brain datasets.

For *O. sativa*, coding sequence (CDS) locations were extracted from the GTF file using awk. Centromeric and transposable element (TE) regions were taken from the DeepPlant study^33^. To quantify the number of cytosines within these features in the genome, we used BEDtools v2.30.0^55^ nuc function. Subsequently, cytosines overlapping centromeric, CDS, and TE regions were identified separately for each tool using BEDtools intersect. We ensured that each genomic location was uniquely assigned to a single feature category by removing overlapping regions between features using the bedtools subtract function. The percentage of cytosines covered by each tool within each genomic feature was calculated based on the number of tool-covered cytosines relative to the total number of cytosines present in that feature across the genome.

### K-mers of disagreement

We looked at K-mers that contribute to the most disagreement with ground-truth EMSeq data for each tool in *O.sativa*. Sites showing more methylation by nanopore tool than EMSeq are defined as overpredicted and sites showing less methylation by nanopore tool than EMSeq are the underpredicted locations. We selected 5000 most overpredicted and underpredicted cytosine locations (CG and nonCG context). As control, we also selected 5000 random locations where the reported and expected methylation levels matched perfectly (“matching sites”). For these groups of sites, we extracted the 10 neighboring base positions (5 upstream and 5 downstream) and visualized the sequence context as tile plots using a custom R script. Each base was represented as a tile, with rows ordered in descending magnitude of disagreement, placing the most discordant sites at the top.

### Calculating F1 scores

F1 score calculation was performed slightly differently for different modifications of interest. For 5mC, EMSeq data (and public bisulfite data for HG002) was used to derive locations that are covered by at least 20 reads and are 100% methylated. We used BEDtools v2.30.0 intersect function for the intersections. These sites were defined as positive sites. A corresponding number of sites which are covered >=20x and are 0% methylated were used as negative sites. For 6mA and 4mC, PacBio data was used as ground truth for three species: *E.coli*, *H.pylori* J99, *T.denticola*. All sites covered >=20x, and where the lower bound of 95% confidence interval of methylated fraction (fracLow) is reported as 1 (indicating 100% methylation), with a p value <0.01 are considered positive sites. An equal number of negative sites were chosen where the IPD Ratio was reported as <1, indicating no deviation from canonical inter-pulse duration. For *H.pylori* 26695 and *A.variabilis*, motifs that are known to be methylated were chosen from REBASE. The reads at positive sites classified as methylated and unmethylated by the tools were counted as true positives (TP) and false negatives (FN) respectively. Similarly, reads classified as methylated and unmethylated from negative sites were treated as false positives (FP) and true negatives (TN) respectively. These values were used to calculate the F1 score, precision, recall, and specificity, employing the following formulae.

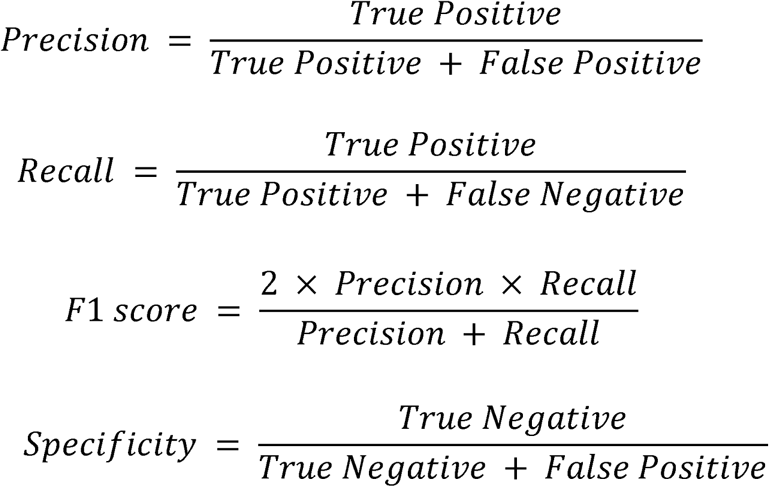

### Studying the effect of probability thresholds

#### Dorado

For Dorado models, we used the Modkit --filter-threshold option to specify the threshold value globally for all modification calls of a given base. Modkit does not define any specific default threshold, and sets it dynamically for each input dataset. The global thresholds, beginning from 0.5 to 0.95 in steps of 0.05, were run on the input BAM files as follows:

~~~
modkit pileup --filter-threshold $threshold --ref $ref --only-tabs --ignore h $in.bam $out_modkit.bed
~~~

#### DeepMod2

DeepMod2 offers the --mod_t and --unmod_t options to set the probability threshold for a per-read prediction to be considered modified or unmodified, respectively. Only predictions with a probability greater than or equal to mod_t, or less than unmod_t are considered for calculation of per-site modification levels. We set the mod_t probability thresholds from 0.5 to 0.95 in steps of 0.05, meanwhile the corresponding unmod_t thresholds were set from 0.5 to 0.05 in decreasing steps of 0.05. The DeepMod2 per-read files were then merged to obtain the per-site output at each threshold, as shown in the command below.

~~~
python deepmod2 merge --prefix $prefix_out --output $outdir --mod_t
$MOD_T --unmod_t $UNMOD_T --input $in.per_read --cpg_out
~~~

#### f5C

The meth-freq -c option of f5C was used to set the call threshold for the log likelihood ratio. The default is 2.5. We analysed the methylation frequencies at different log likelihood ratios, starting from 0 to 10, in increments of 1 log likelihood ratio. Stranded readwise methylation calls were used as input. A custom script was used to account for strandedness while aggregating the methylation calls. Performance metrics were calculated on the methylation frequencies for each log likelihood ratio.

~~~
f5c_x86_64_linux_cuda meth-freq -c $log_likelihood_ratio -i
$call_meth_stranded.tsv -s -o $meth_freq_out.tsv
~~~

#### RockFish

The default RockFish’s script to calculate methylation frequencies (calculate_site_freq.py) has a hard-coded probability threshold and does not allow users to change it directly. Hence, we created a modified in-house script where this threshold could be provided as a command-line argument. With this we were able to profile the methylation frequencies at probability thresholds of 0.5 to 0.95 to identify modified calls, and corresponding thresholds of 0.5 to 0.05 to identify unmodified calls, in increments and decrements of 0.05.

~~~
python rockfish_aggregate2.py -i $predictions.tsv -u $MOD_T -l
$UNMOD_T -o $meth_freq_out.tsv
DeepPlant/DeepBAM
~~~

DeepBAM includes an aggregation script (scripts/deepbam_aggregate.py) that allows users to select the threshold used for aggregating the readwise data info sitewise data. Since the column order of both DeepPlant and DeepBAM are the same, this script works for results generated by either tool. The script uses the set threshold value as a simple filter to split the reads as methylated or unmethylated. The output of this script is a single tsv file that contains the number of methylated reads for a given threshold at a given coverage, where the coverage remains a constant for a given locus.

~~~
python scripts/deepbam_aggregate.py --file_path
$deepbam/plant_readwise.tsv --threshold 0.5 --aggregation_output
$output.tsv
~~~

Additionally, the number of reads at sites that are expected to be 0% (negative sites) and 100% (positive sites) methylated were counted to account for data loss at each threshold.

### Site level evaluation

For 5mC, only those sites which were covered by at least 20 reads in both EMSeq and ONT data were considered for evaluation. For 6mA and 4mC, where PacBio data was available, locations covered >=20x in both PacBio and ONT data were evaluated. In the other cases, sites which were covered >=20x in ONT data were considered. Pearson correlation coefficient was used to measure the correlation between the ground truth and the values reported by various tools, using the cor() function in R. Mean Absolute Error (MAE) and Rooted Mean Squared Error (RMSE) were calculated using the mae() and rmse() functions from the Metrics package in R. The concordance between EMSeq and ONT data is visualized as a heatmap using the geom_bin2d() function of the ggplot2 package in R.

For 6mA, we analyzed site-level concordance by generating a mixed dataset of *E.coli*. Reads from Ecoli_WT and Ecoli_DM were used as methylated and unmethylated respectively. A BED file containing GATC sites was first generated using ‘modkit motif bed’. After this, the BED file was filtered for locations that were 20KB apart; this ensures that reads that span one motif would not span other GATC locations, which is crucial for the subsequent steps. All GATC sites for the first and last 500kb of the genome were assigned a target fraction of 100% and 0% methylation respectively. Each of the remaining filtered sites is then assigned a random fraction between 0 and 1. Reads from WT and DM are combined to achieve this random fraction of methylation level, keeping track of read ids from each data. Once this step is completed for all filtered sites, the expected methylation levels of all the GATC sites in the genome are calculated based on the read ids that map to each site.

### Studying the effect of coverage

We evaluated the effect of subsampling reads at site-level, as at read-level, the classification is always the same for each read. We used data from *O.sativa japonica* to evaluate the effect of coverage on CpG methylation calling, as this was the dataset that had the best coverage overall, and was also the dataset on which we could successfully run all the tools on the full data, including DeepBAM and DeepPlant. We fractionated the reads as 5% to 100% of the total data. The subsampling was done using rhyper() function in R, and was repeated 5 times for each tool using 5 different seeds to offset any sampling biases.We restricted the performance evaluation to those sites covered by at least 20 reads in the full dataset. Average Pearson R and RMSE values of the 5 subsamples were calculated, and visualized as line plots.

### Evaluating the effect of read-quality

NanoPlot^52^ was used to generate a readwise file containing the average quality of each read. This was filtered into two bins, one where the average quality was greater than Q20, and another where the full dataset was considered. These were then further split into their respective quartile 1 (bin with bottom 25% reads by quality) and quartile 4 (bin with top 25% reads). Once the bins were prepared the respective bam files were subset and the modbed files were regenerated to evaluate the 6mA calling performance. Subsequently the BED files were annotated for the fasta regions they spanned, the motifs were extracted and the methylation rate was compared species and motif wise.

### Studying the effect of neighboring modifications

To assess the effect of neighboring methylated cytosines, we analyzed datasets from *E. coli*, *H. pylori* strain J99, *O. sativa*, and HG002. For 6mA modifications, only bacterial datasets were used. Locations covered by at least 20 reads were retained for the analysis. We evaluated the influence of methylation on ten adjacent base positions surrounding the methylated motif (5 upstream and 5 downstream). To minimize the confounding effects of nearby modifications, particularly at longer distances, all intervening methylated bases between the motif and the base of interest were filtered out. The impact of methylation on neighboring positions was visualized as the difference between methylation percentages from EMseq and corresponding tool. Methylation percentage was visualised wherever EMseq was not available. Visualization was done using line and box plots generated with the geom_boxplot, geom_line, and geom_point functions from the ggplot2 package in R. Additionally, context-specific jitter-box plots were created using the geom_half_boxplot and geom_half_point functions from the gghalves package, with sequence context information obtained via a custom script.

### Evaluation of computational performance

The runtime, CPU usage, and maximum RAM consumed by various tools and models was evaluated on various bacterial datasets using the gnu “time -v” command. From the output, wall_time, percent_cpu, and max_res_size were recorded. The runtime was normalized for the size of data, and reported as megabases processed per second. Values reported are arithmetic mean for all the bacterial data (8 datasets in total). All tools were run with 40 CPU cores (see commands in Methylation calling section above). All tests were done on a server with a dual Intel Xeon Gold 6548Y+ processor, 256GB RAM, 2x L40s GPUs running Ubuntu 22.04.5 LTS.

## Author contributions

OK, KBT, and DTS designed the study. OK, RJM, RJ performed data analysis with help from NKS. LZ and SA performed gDNA isolations and nanopore sequencing. TN performed EMSeq. KBT and DTS acquired funding. OK, RJM, RJ, KBT and DTS wrote the manuscript. All authors read and approved the manuscript.

## Acknowledgements

The authors acknowledge Dr Santosh Kumar and Apoorva Etikala for the genomic DNA of *Helicobacter pylori* str. 26695, Dr Khader Valli Rupanagudi and Dr Reddy P Kommaddi from Centre for Brain Research, Bengaluru, for providing the GM24385 (HG002) GIAB cell line, Dr Hitendra Patel and Anjana Sharma for genomic DNA of *Oryza sativa japonica*, Dr Imran Siddiqi and Sai Kiran Ginkuntla for the genomic DNA of *Arabidopsis thaliana*, and Dr P Chandrasekhar and Niharika Tiwary for mouse brain tissue and embryonic stem cells. Valli Undamatla from the NGS facility of CCMB is acknowledged for her help in EMSeq, and Rakeshpal Bhagat for his help with nanopore sequencing. We acknowledge Dr Shambhavi Garde for the YejM amplicon and Sathya Swetha Na for the CYP21A2 amplicon. We acknowledge the CCMB High-Performance Computing (HPC) facility and CSIR-4PI for providing the computational resources used in this study, and Geetha Thanu and Aparna Kumari for their technical support. We are thankful to Mainak Chakraborty, Beryl Rabindran, and AWS for enabling us to host the datasets on the Registry of Open Data on AWS.

## Data Availability

Raw signal data in POD5 format, as well as processed data is available via the Registry of Open Data hosted on AWS at https://registry.opendata.aws/ont_basemod_data/. Reference aligned Modbam files generated in this study are deposited to the NCBI SRA under the BioProject number PRJNA1172918. Nextflow and Snakemake pipelines to reproduce the analysis or benchmark new data, and code used to process and visualize data from this work is available at https://github.com/SowpatiLab/ONT_Methylation_Benchmarking.

Other data sources:

*O.sativa* GTF file: https://ftp.ncbi.nlm.nih.gov/genomes/all/GCF/034/140/825/GCF_034140825.1_ASM3414082v1/ GCF_034140825.1_ASM3414082v1_genomic.gtf.gz

GRCm39/mm39 assembly gene annotations: https://genome.ucsc.edu/cgi-bin/hgTables Rerio repository: https://github.com/nanoporetech/rerio

HG002 WGBS data: s3://ont-open-data/gm24385_mod_2021.09/bisulphite/bismark/Bacterial PacBio data: https://downloads.pacbcloud.com/public/dataset/2021-11-Microbial-96plex/demultiplexed-reads/

Reviewer access token: https://dataview.ncbi.nlm.nih.gov/object/PRJNA1172918?reviewer=trd0039k0pg7sm6u45j4e4nhui

## Code Availability

All custom code used in this work is deposited to a GitHub repository and is accessible at https://github.com/SowpatiLab/ONT_Methylation_Benchmarking. Nextflow and Snakemake pipelines to reproduce the analysis or benchmark new data is also available at the same GitHub link.

## Supplementary Information

### Supplementary Tables

Table S1: Conversion efficiency of EMSeq datasets as measured on exogenous spike-ins

Table S2: Nanopore and EMSeq datasets generated in this study and their quality metrics

Table S3: Sequence contexts expected to carry methylation in various bacterial species used in this study

Table S4: Representation of 9-mers and 11-mers covered by the datasets used for evaluation of 5mC

Table S5: Overall abundance of various marks in the species and datasets used in the study

Table S6: Versions of models used for calling methylation using various tools

Table S7: Read level performance metrics and associated details across the tools and datasets

Table S8: Effect of changing probability thresholds on various F1 metrics, and corresponding data loss

Table S9: Site level evaluation metrics for 5mC calls across the datasets

Table S10: Computational performance metrics of various tools utilized in this study

Table S11: Reference genome versions used in the study

### Supplementary Figures

Figure S1: Validation of different *E.coli* samples using EMSeq data

Figure S2: Sequencing coverage and quality across various samples used for benchmarking

Figure S3: Variance in performance of different model-subtypes offered by each tool on bacterial datasets

Figure S4: Read-level evaluation of various tools

Figure S5: Read-level evaluation of the effect of probability thresholds on DeepPlant’s performance

Figure S6: Site-level evaluation of methylation calling tools using correlation heatmaps at CpG sites for Arabidopsis and Human data

Figure S7: Site-level evaluation of methylation calling tools using correlation heatmaps in CpG context for mouse brain and ESC data

Figure S8: CpG methylation calling around Transcription Start Sites (mouse) and correlation heatmaps on rice data

Figure S9: K-mers of disagreement at CpG locations on rice data

Figure S10: Site-level evaluation of methylation calling tools using correlation heatmaps in non-CG context, on plant data

Figure S11: Site-level evaluation of non-CG methylation using correlation heatmaps, on human HG002 and mouse brain data

Figure S12: Site-level evaluation of non-CG methylation using correlation heatmaps on mouse ESC data, and effect of adjusting probability thresholds on false positives

Figure S13: Context-wise evaluation of 6mA calling tools

Figure S14: Context-wise evaluation of 4mC calling tools

Figure S15: Evaluating the effect of neighboring modifications for 5mCG calling tools

Figure S16: Evaluating the effect of neighboring modifications in various contexts

Figure S17: Evaluating the effect of neighboring CG methylation on rice data

Figure S18: Evaluating the effect of neighboring CG methylation on E.coli and human data

Figure S19: Effect of two neighboring methylated CpGs

Figure S20: Evaluating the effect of neighboring modifications on adenine methylation calling

Figure S21: Analysis of the new v5.2 5mC models of Dorado

Figure S22: Analysis of the new v5.2 6mA and 4mC models of Dorado

Figure S23: Evaluating the effect of neighboring modifications on performance of v5.2 models

Figure S24: Read-level comparison of Dorado 5mCG, 6mA and 4mC high-accuracy (hac) and super-accuracy (sup) models.

Figure S25: Sites profiled by EMSeq and Nanopore, and susceptibility of 4kHz and 5Hz models to neighboring methylation

## References

1. Jones, P. A. Functions of DNA methylation: islands, start sites, gene bodies and beyond. Nat. Rev. Genet. 13, 484–492 (2012).

2. Schübeler, D. Function and information content of DNA methylation. Nature 517, 321–326 (2015).

3. Smith, Z. D. & Meissner, A. DNA methylation: roles in mammalian development. Nat. Rev. Genet. 14, 204–220 (2013).

4. Schultz, M. D. et al. Human body epigenome maps reveal noncanonical DNA methylation variation. Nature 523, 212–216 (2015).

5. Zhou, W. et al. DNA methylation loss in late-replicating domains is linked to mitotic cell division. Nat. Genet. 50, 591–602 (2018).

6. Hofer, A., Liu, Z. J. & Balasubramanian, S. Detection, Structure and Function of Modified DNA Bases. J. Am. Chem. Soc. 141, 6420–6429 (2019).

7. Luo, C., Hajkova, P. & Ecker, J. R. Dynamic DNA methylation: In the right place at the right time. Science 361, 1336–1340 (2018).

8. Jang, H. S., Shin, W. J., Lee, J. E. & Do, J. T. CpG and Non-CpG Methylation in Epigenetic Gene Regulation and Brain Function. Genes 8, 148 (2017).

9. Wang, Z. & Baulcombe, D. C. Transposon age and non-CG methylation. Nat. Commun. 11, 1221 (2020).

10. Zhang, H., Lang, Z. & Zhu, J.-K. Dynamics and function of DNA methylation in plants. Nat. Rev. Mol. Cell Biol. 19, 489–506 (2018).

11. Au, K. G., Welsh, K. & Modrich, P. Initiation of methyl-directed mismatch repair. J. Biol. Chem. 267, 12142–12148 (1992).

12. Arber, W. & Dussoix, D. Host specificity of DNA produced by Escherichia coli: I. Host controlled modification of bacteriophage λ. J. Mol. Biol. 5, 18–36 (1962).

13. Boulias, K. & Greer, E. L. Means, mechanisms and consequences of adenine methylation in DNA. Nat. Rev. Genet. 23, 411–428 (2022).

14. Luo, G.-Z., Blanco, M. A., Greer, E. L., He, C. & Shi, Y. DNA N6-methyladenine: a new epigenetic mark in eukaryotes? Nat. Rev. Mol. Cell Biol. 16, 705–710 (2015).

15. Rodriguez, F., Yushenova, I. A., DiCorpo, D. & Arkhipova, I. R. Bacterial N4-methylcytosine as an epigenetic mark in eukaryotic DNA. Nat. Commun. 13, 1072 (2022).

16. Krueger, F., Kreck, B., Franke, A. & Andrews, S. R. DNA methylome analysis using short bisulfite sequencing data. Nat. Methods 9, 145–151 (2012).

17. Ji, L. et al. Methylated DNA is over-represented in whole-genome bisulfite sequencing data. Front. Genet. 5, (2014).

18. Grunau, C., Clark, S. J. & Rosenthal, A. Bisulfite genomic sequencing: systematic investigation of critical experimental parameters. Nucleic Acids Res. 29, e65 (2001).

19. Beaulaurier, J., Schadt, E. E. & Fang, G. Deciphering bacterial epigenomes using modern sequencing technologies. Nat. Rev. Genet. 20, 157–172 (2019).

20. Flusberg, B. A. et al. Direct detection of DNA methylation during single-molecule, real-time sequencing. Nat. Methods 7, 461–465 (2010).

21. Ip, C. L. C., et al. MinION Analysis and Reference Consortium: Phase 1 data release and analysis. Preprint at 10.12688/f1000research.7201.1 (2015).

22. Simpson, J. T. et al. Detecting DNA cytosine methylation using nanopore sequencing. Nat. Methods 14, 407–410 (2017).

23. Goyal, P. et al. Structural and mechanistic insights into the bacterial amyloid secretion channel CsgG. Nature 516, 250–253 (2014).

24. Van der Verren, S. E. et al. A dual-constriction biological nanopore resolves homonucleotide sequences with high fidelity. Nat. Biotechnol. 38, 1415–1420 (2020).

25. Oxford Nanopore [@nanopore]. Improvements in latest MinKNOW update: •5kHz sampling rate replaces 4kHz sampling rate - for improved accuracy •POD5 is now the default raw data file type •Addition of 5mC/5hmC Remora Model •Automatically stop your runs with Run Until Learn more: https://bit.ly/3MFtbHf https://t.co/2Pizs3LP1z. *Twitter* https://x.com/nanopore/status/1659469919971360769 (2023).

26. nanoporetech/dorado. Oxford Nanopore Technologies (2024).

27. Transforming Basecalling in Genomic Sequencing. Oxford Nanopore Technologies https://nanoporetech.com/blog/transforming-basecalling-in-genomic-sequencing (2024).

28. London Calling 2024: Update from Clive Brown. (2024).

29. Stanojević, D., Li, Z., Bakić, S., Foo, R. & Šikić, M. Rockfish: A transformer-based model for accurate 5-methylcytosine prediction from nanopore sequencing. Nat. Commun. 15, 5580 (2024).

30. Ahsan, M. U., Gouru, A., Chan, J., Zhou, W. & Wang, K. A signal processing and deep learning framework for methylation detection using Oxford Nanopore sequencing. Nat. Commun. 15, 1448 (2024).

31. Gamaarachchi, H. et al. GPU accelerated adaptive banded event alignment for rapid comparative nanopore signal analysis. BMC Bioinformatics 21, 343 (2020).

32. Bai, X. et al. DeepBAM: a high-accuracy single-molecule CpG methylation detection tool for Oxford nanopore sequencing. 10.1093/bib/bbae413.

33. Chen, H.-X. et al. Accurate cross-species 5mC detection for Oxford Nanopore sequencing in plants with DeepPlant. Nat. Commun. 16, 3227 (2025).

34. Yuen, Z. W.-S. et al. Systematic benchmarking of tools for CpG methylation detection from nanopore sequencing. Nat. Commun. 12, 3438 (2021).

35. Liu, Y. et al. DNA methylation-calling tools for Oxford Nanopore sequencing: a survey and human epigenome-wide evaluation. Genome Biol. 22, 295 (2021).

36. Sigurpalsdottir, B. D. et al. A comparison of methods for detecting DNA methylation from long-read sequencing of human genomes. Genome Biol. 25, 69 (2024).

37. Genner, R. et al. Assessing DNA methylation detection for primary human tissue using Nanopore sequencing. Genome Res. 35, 632–643 (2025).

38. Halliwell, D. O., Honig, F., Bagby, S., Roy, S. & Murrell, A. Double and single stranded detection of 5-methylcytosine and 5-hydroxymethylcytosine with nanopore sequencing. Commun. Biol. 8, 1–13 (2025).

39. Fu, Y., Timp, W. & Sedlazeck, F. J. Computational analysis of DNA methylation from long-read sequencing. Nat. Rev. Genet. 1–15 (2025) doi:10.1038/s41576-025-00822-5.

40. Blow, M. J. et al. The Epigenomic Landscape of Prokaryotes. PLOS Genet. 12, e1005854 (2016).

41. Tourancheau, A., Mead, E. A., Zhang, X.-S. & Fang, G. Discovering multiple types of DNA methylation from bacteria and microbiome using nanopore sequencing. Nat. Methods 18, 491–498 (2021).

42. Krebes, J. et al. The complex methylome of the human gastric pathogen Helicobacter pylori. Nucleic Acids Res. 42, 2415–2432 (2014).

43. Kulkarni, O. et al. NEMO: Improved and accurate models for identification of 6mA using Nanopore sequencing. 2024.03.12.584205 Preprint at 10.1101/2024.03.12.584205 (2024).

44. Roberts, R. J. et al. SURVEY AND SUMMARY: A nomenclature for restriction enzymes, DNA methyltransferases, homing endonucleases and their genes. Nucleic Acids Res. 31, 1805–1812 (2003).

45. nanoporetech. v5.2 DNA 5mC model not reporting non-CpG methylation · Issue #1440 · nanoporetech/dorado. GitHub https://github.com/nanoporetech/dorado/issues/1440.

46. Dao, T., Fu, D. Y., Ermon, S., Rudra, A. & Ré, C. FlashAttention: Fast and Memory-Efficient Exact Attention with IO-Awareness. Preprint at 10.48550/arXiv.2205.14135 (2022).

47. Martin, M. Cutadapt removes adapter sequences from high-throughput sequencing reads. EMBnet.journal 17, 10–12 (2011).

48. Krueger, F. & Andrews, S. R. Bismark: a flexible aligner and methylation caller for Bisulfite-Seq applications. Bioinformatics 27, 1571–1572 (2011).

49. SMRT Link. PacBio https://www.pacb.com/smrt-link/.

50. Li, H. et al. The Sequence Alignment/Map format and SAMtools. Bioinformatics 25, 2078–2079 (2009).

51. Li, H. Minimap2: pairwise alignment for nucleotide sequences. Bioinformatics 34, 3094–3100 (2018).

52. De Coster, W. & Rademakers, R. NanoPack2: population-scale evaluation of long-read sequencing data. Bioinformatics 39, btad311 (2023).

53. Pedersen, B. S. & Quinlan, A. R. Mosdepth: quick coverage calculation for genomes and exomes. Bioinformatics 34, 867–868 (2018).

54. Hall, M. B. Rasusa: Randomly subsample sequencing reads to a specified coverage. J. Open Source Softw. 7, 3941 (2022).

55. Quinlan, A. R. & Hall, I. M. BEDTools: a flexible suite of utilities for comparing genomic features. Bioinformatics 26, 841–842 (2010).

