## Supplementary Figures for "Comprehensive benchmarking of tools for nanopore-based detection of DNA methylation"

### Fig S1

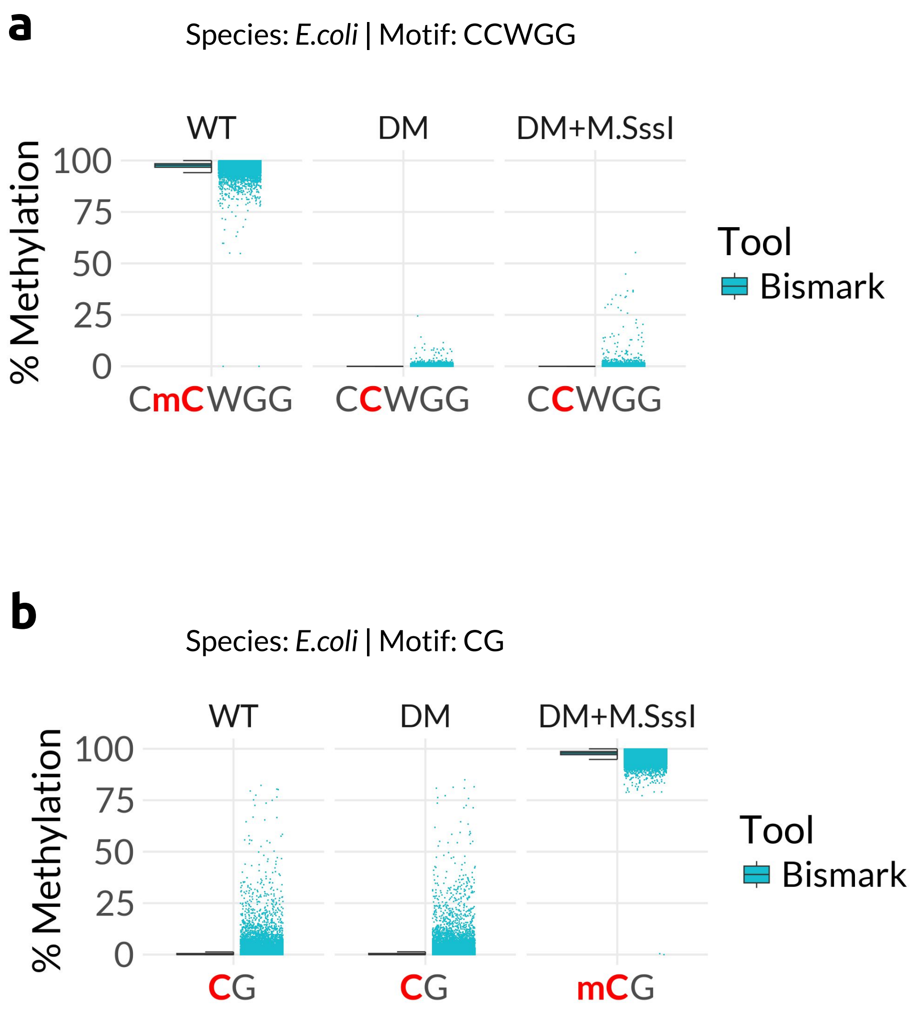


Fig S1: Validation of different *E.coli* samples using EMSeq data (Bismark) visualized as Jitter-box plot. In Jitter-box plots, boxplot shows distribution of the data with quartile range and median value. The jitter part represents percent methylation for each site profiled as a dot. The color of the dot represents the tool used to generate that data. a) Jitter-box plot showing methylation percentage in CCWGG context for *E.coli* (WT, DM, DM+M.SssI) EMSeq data. b) Jitter-box plot showing methylation percentage in CpG context for *E.coli* (WT, DM, DM+M.SssI) EMSeq data.

Fig S2

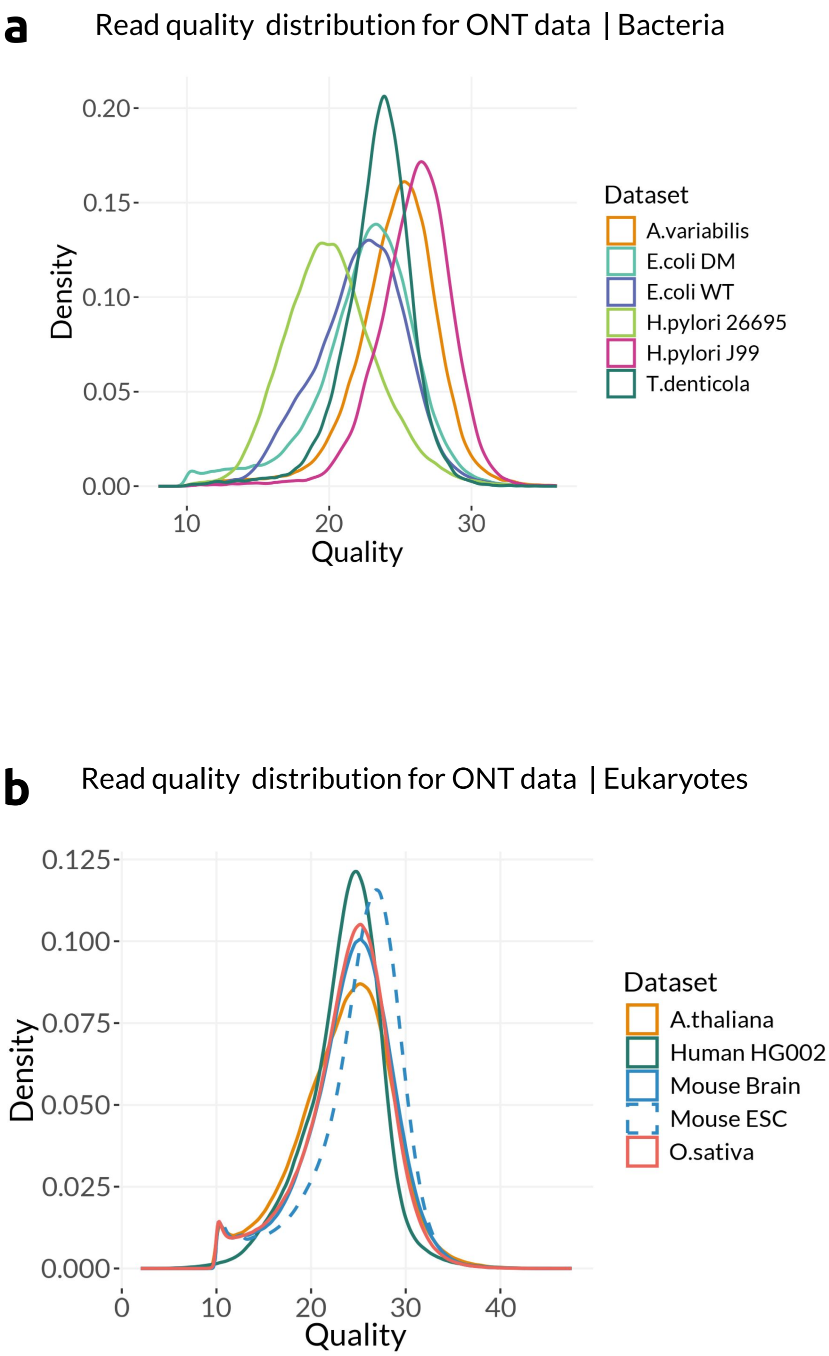


Fig S2: Sequencing quality across various samples used for benchmarking.
Density plots showing the variance of quality score distribution across ONT data for a) bacterial samples, and b) eukaryotic samples used in this study.

### Fig S3

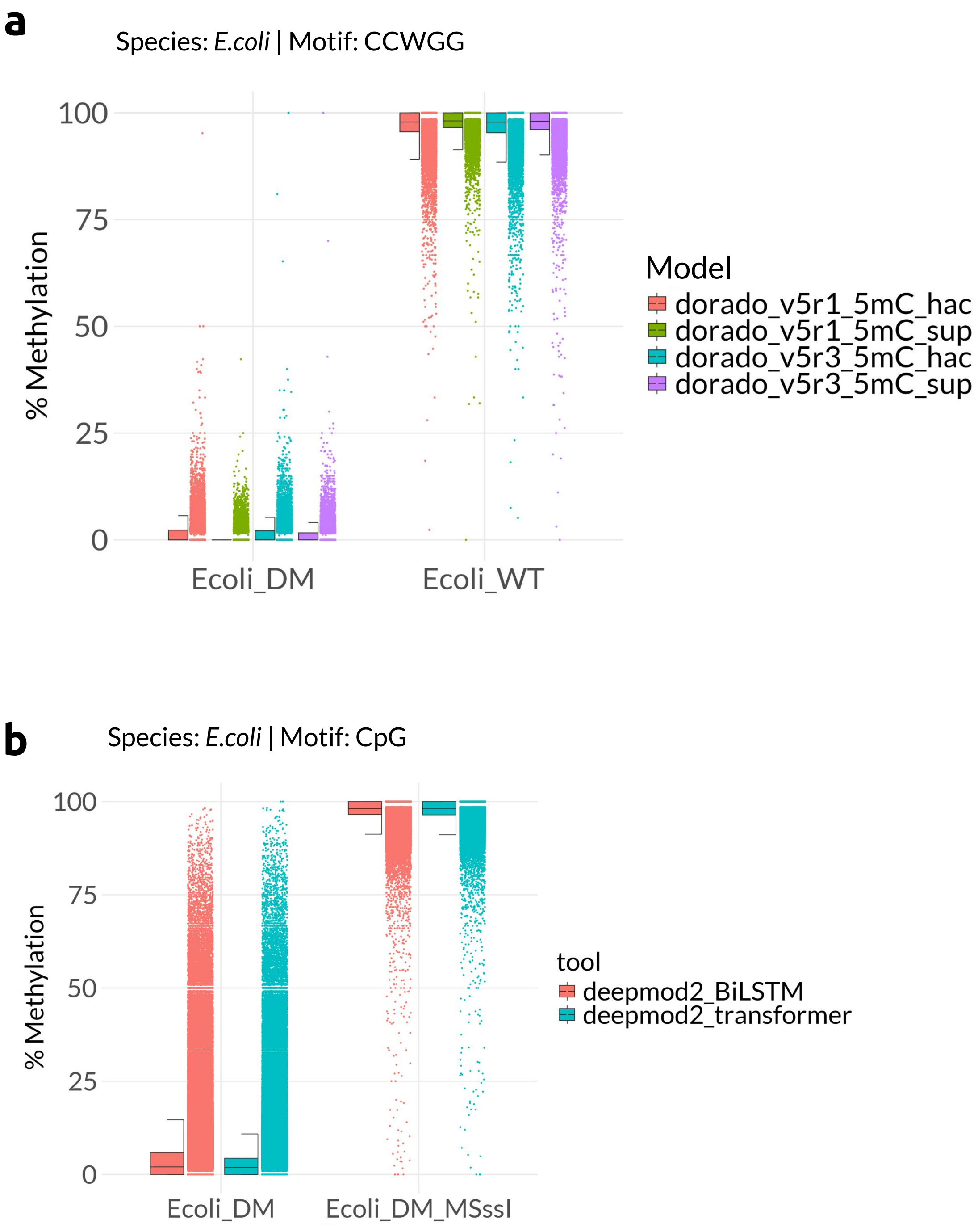


Fig S3: Variance in performance of different model-subtypes offered by each tool on bacterial datasets. a) Jitter-box plot depicting the CcWGG context performance differences between the high-accuracy (hac) and super-accuracy (sup) model variants offered by Dorado for *E.coli* WT and DM samples. b) Jitter-box plot depicting the CpG context performance differences between the BiLSTM and transformer model architectures offered by DeepMod2 in *E.coli* DM and DM+M.SssI samples.

### Fig S4

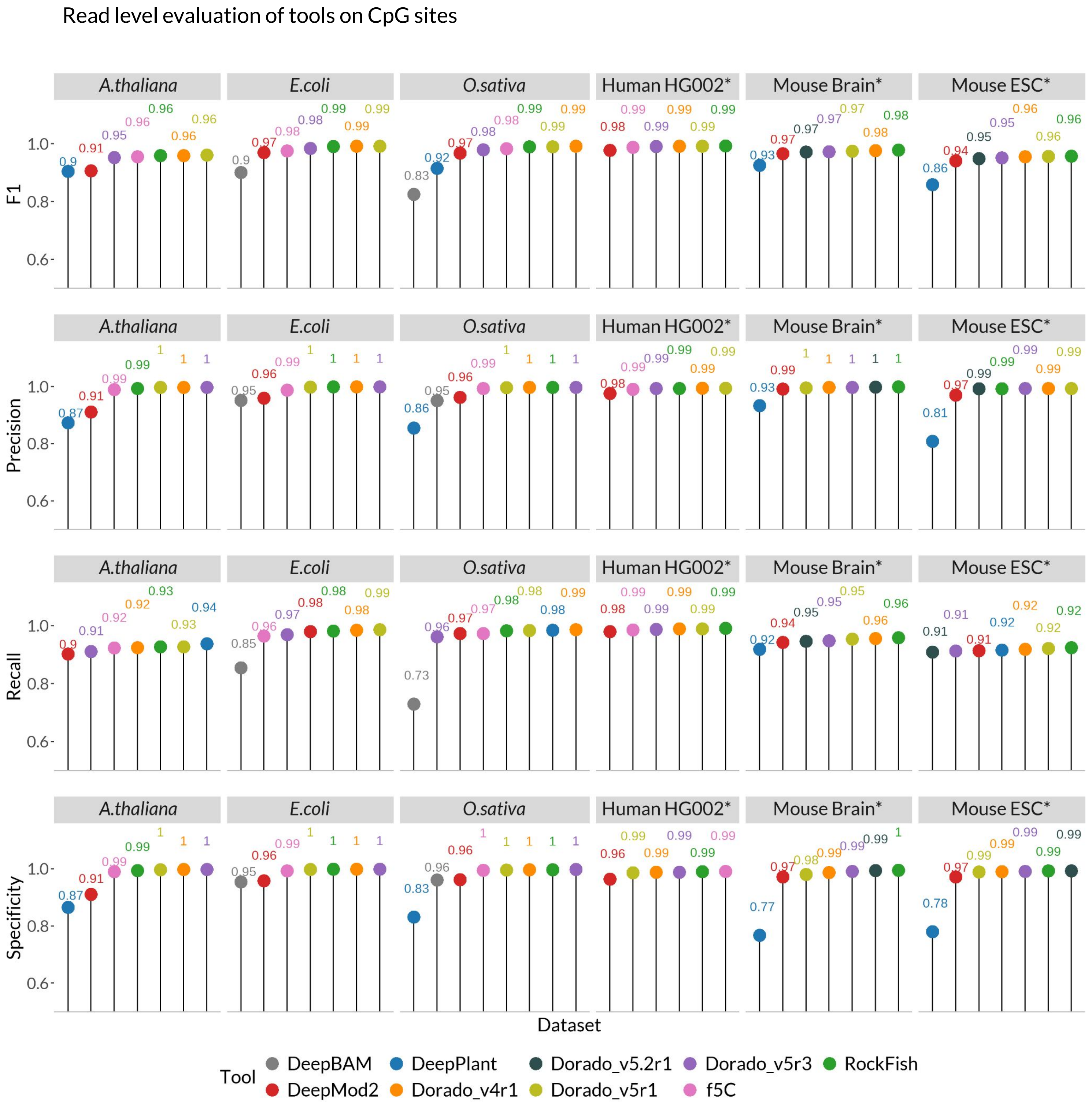


Fig S4: Read level evaluation of various tools. Performance metrics of tools on CpG sites. F1, Precision, Recall, and Specificity are represented as lollipops colored by tools for each dataset. The score is also indicated next to each lollipop for clarity. The asterisk next to Human HG002, mouse brain and ESC indicates that the calculations are done using data of only chromosome 1. The y-axis is scaled from 0.5 instead of 0 to highlight subtle differences in the scores.

### Fig S5

Effect of probability thresholds on DeepPlant's performance

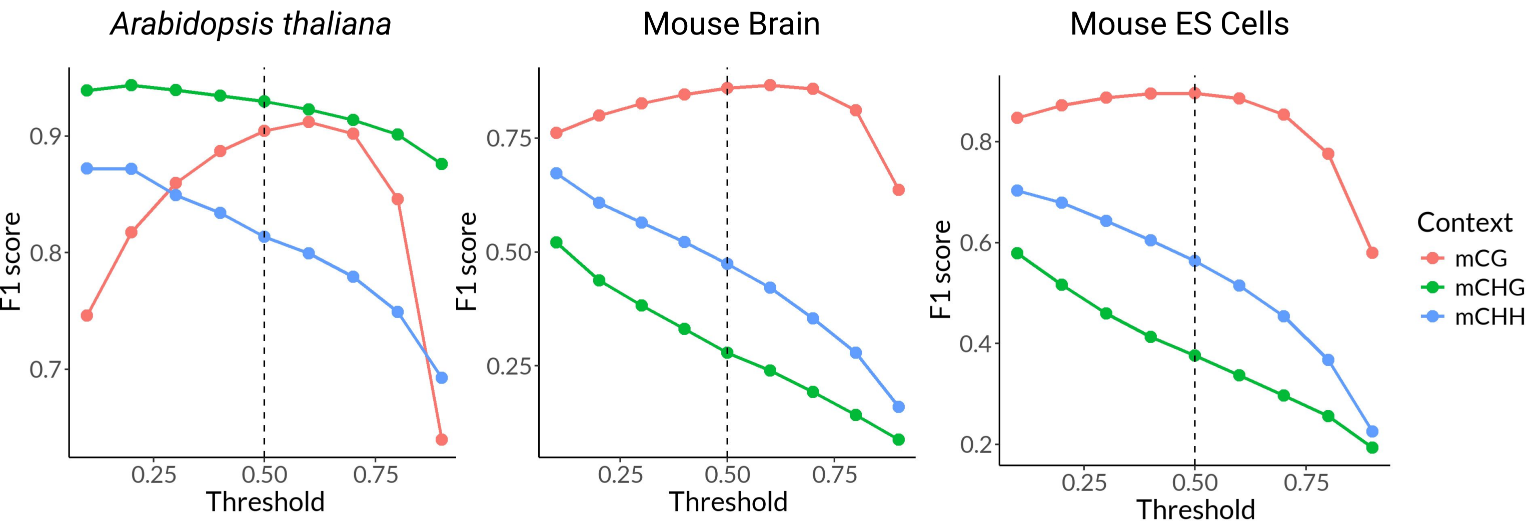


Fig S5: Read-level evaluation of the effect of probability thresholds on DeepPlant's performance. The F1 scores (y-axis) are plotted in a line plot against the thresholds (x-axis) of DeepPlant for CG, CHG, and CHH contexts of *A.thaliana*, mouse brain and ES cells. The vertical dashed line indicates the default threshold (0.5) used by DeepPlant.

### Fig S6

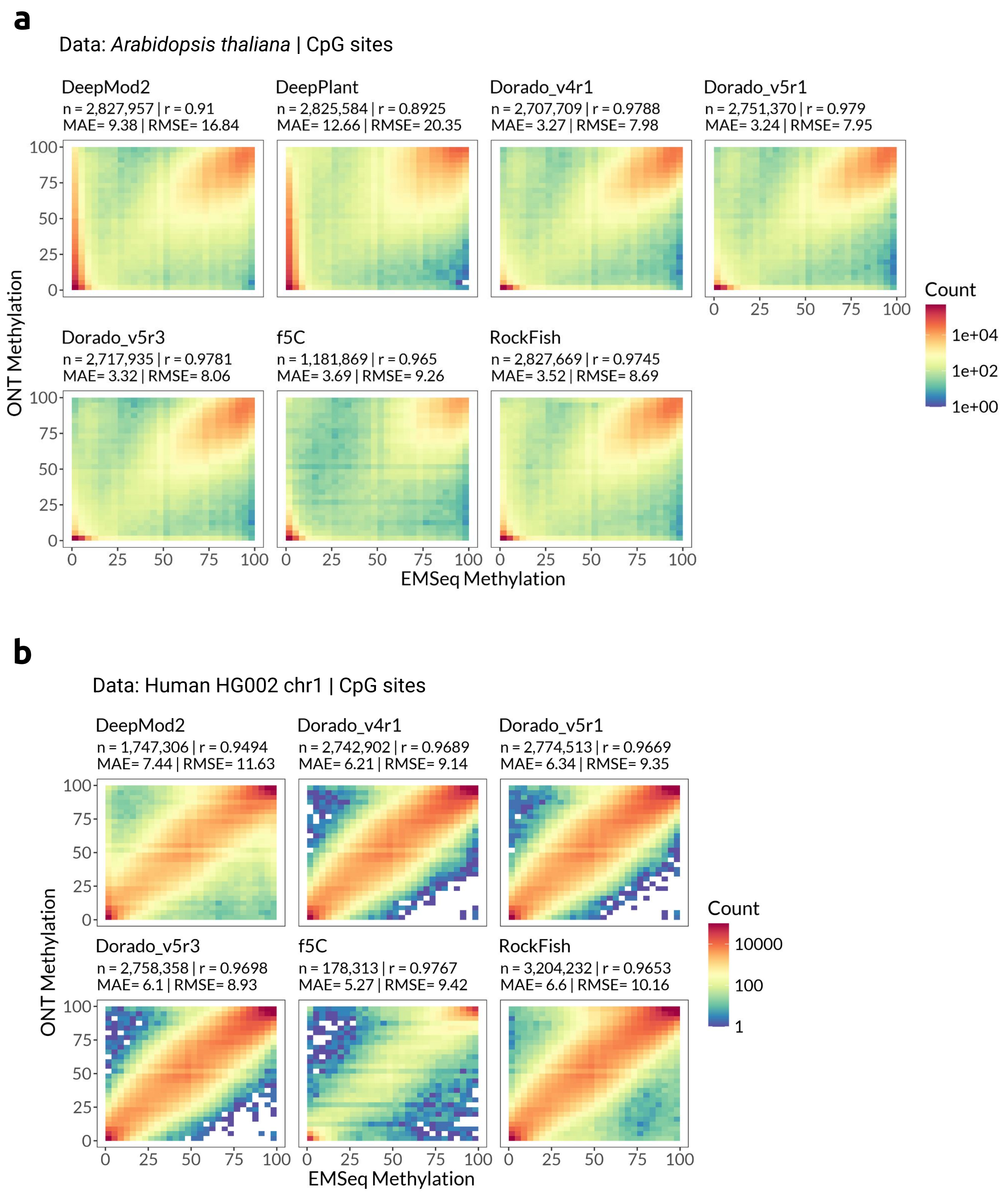


Fig S6: Site-level evaluation of methylation calling tools using correlation heatmaps at CpG sites for Arabidopsis and Human data. n = number of CpG sites profiled and covered by >=20 reads; r = Pearson correlation coefficient; MAE = Mean Absolute Error; RMSE = Root Mean-Squared Error. a-b) Correlation heatmaps of CpG sites from *A.thaliana* data (a) and human chromosome 1 of HG002 data (b), for each tool compared to EMSeq data (ground truth).

### Fig S7

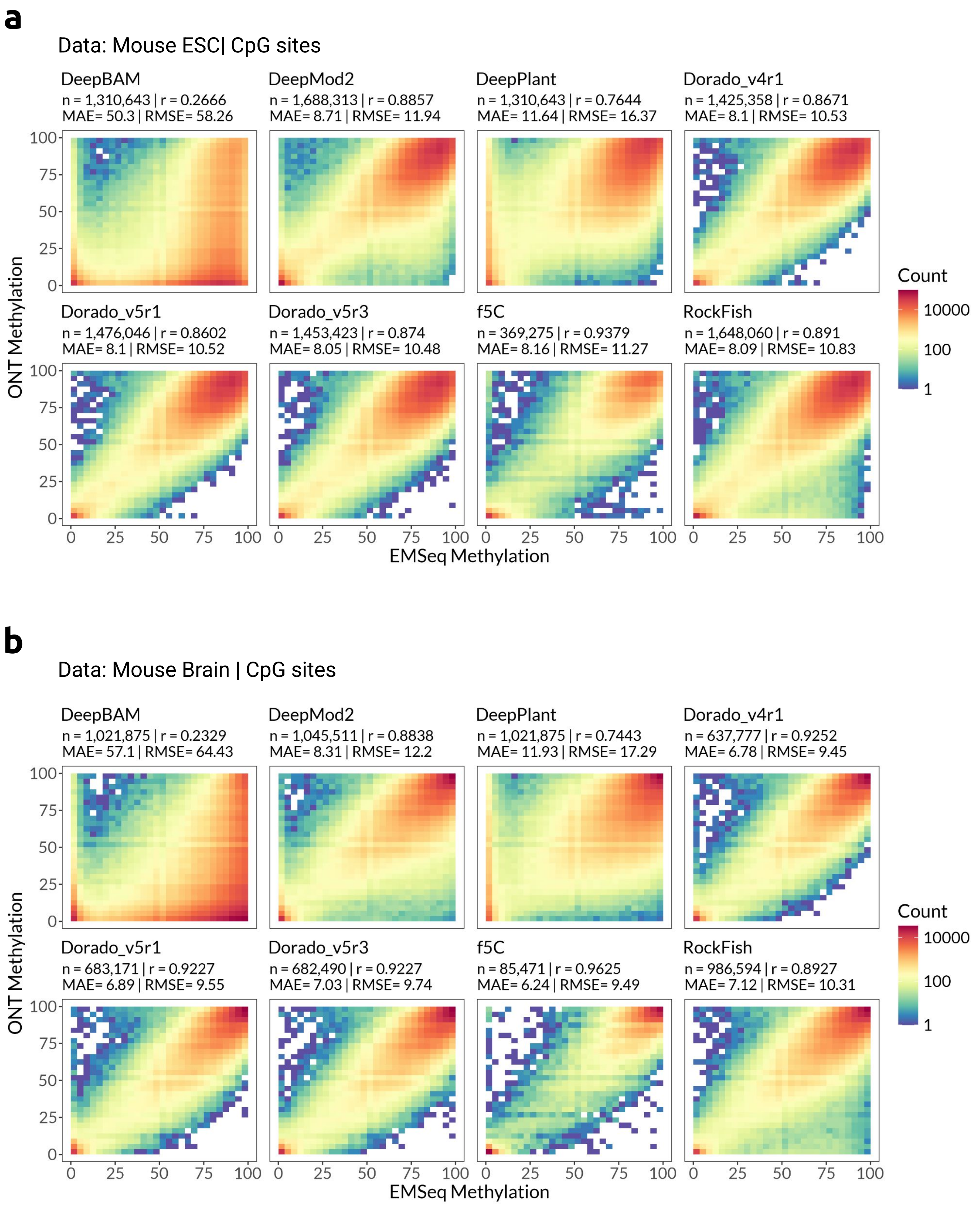


Fig S7: Site-level evaluation of methylation calling tools using correlation heatmaps in CpG context for mouse brain and ESC data. n = number of CpG sites profiled and covered by >=20 reads; r = Pearson correlation coefficient; MAE = Mean Absolute Error; RMSE = Root Mean-Squared Error. a-b) Correlation heatmaps of CpG sites from mouse embryonic stem cells (a) and mouse brain (b) data, for each tool compared to EMSeq data (ground truth)

### Fig S8

**a**

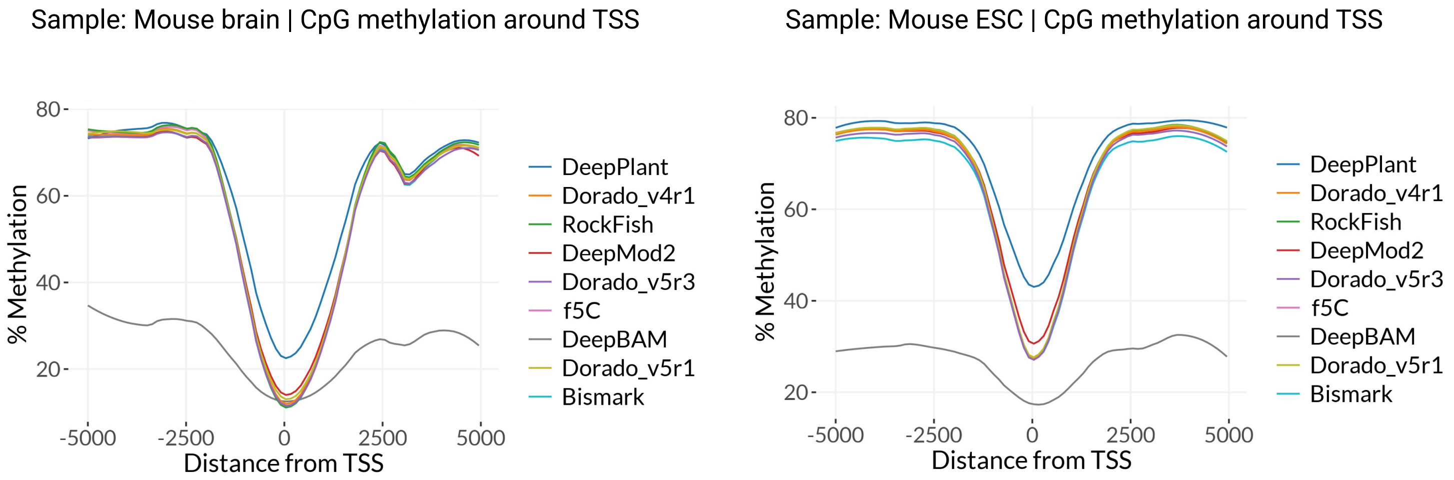


**b**

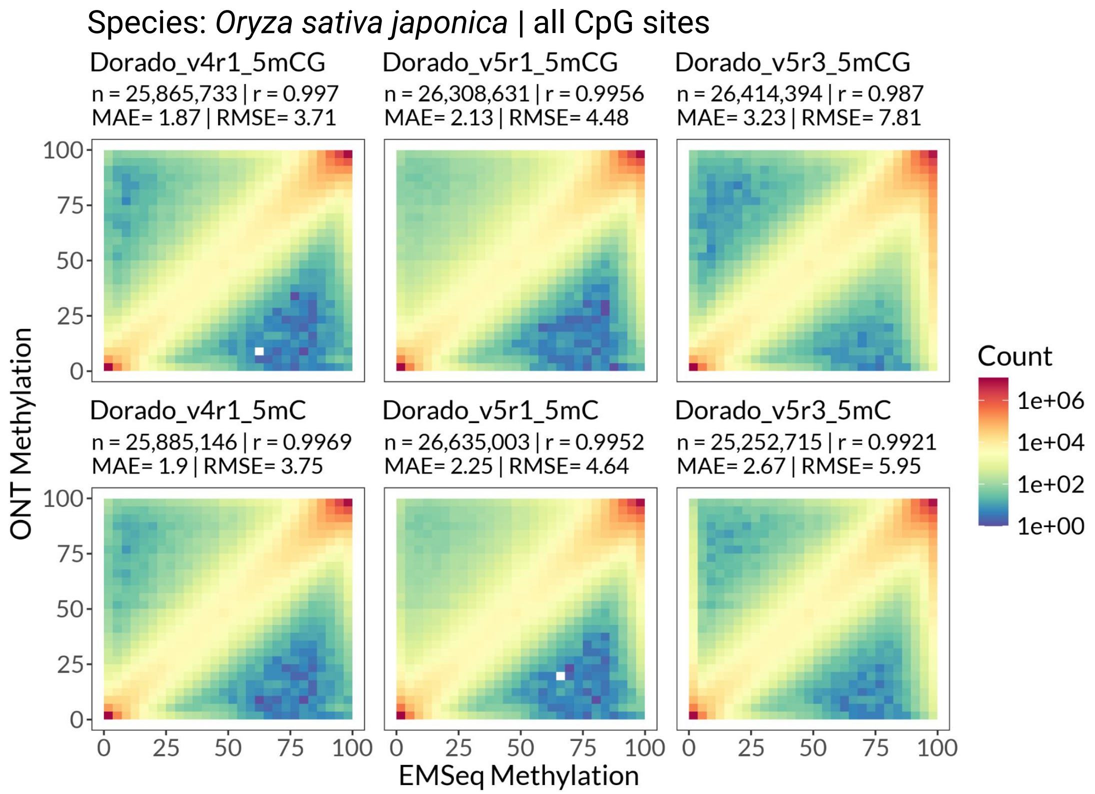


Fig S8: CpG methylation calling around Transcription Start Sites (mouse) and correlation heatmaps on rice data. a) The methylation percentage of CpG sites spanning 5kb upstream and downstream of the transcription start sites (TSS) on chromosome 1 plotted as a line plot for each methylation tool, for mouse brain and ESC data. b) Evaluation of CpG-sites from rice data. n = number of CpG sites profiled and covered by >=20 reads; r = Pearson correlation coefficient; MAE = Mean Absolute Error; RMSE = Root Mean-Squared Error. Correlation heatmap of all CpG sites profiled by each of the Dorado all-context 5mC and 5mCG models compared to the EMSeq ground truth.

### Fig S9

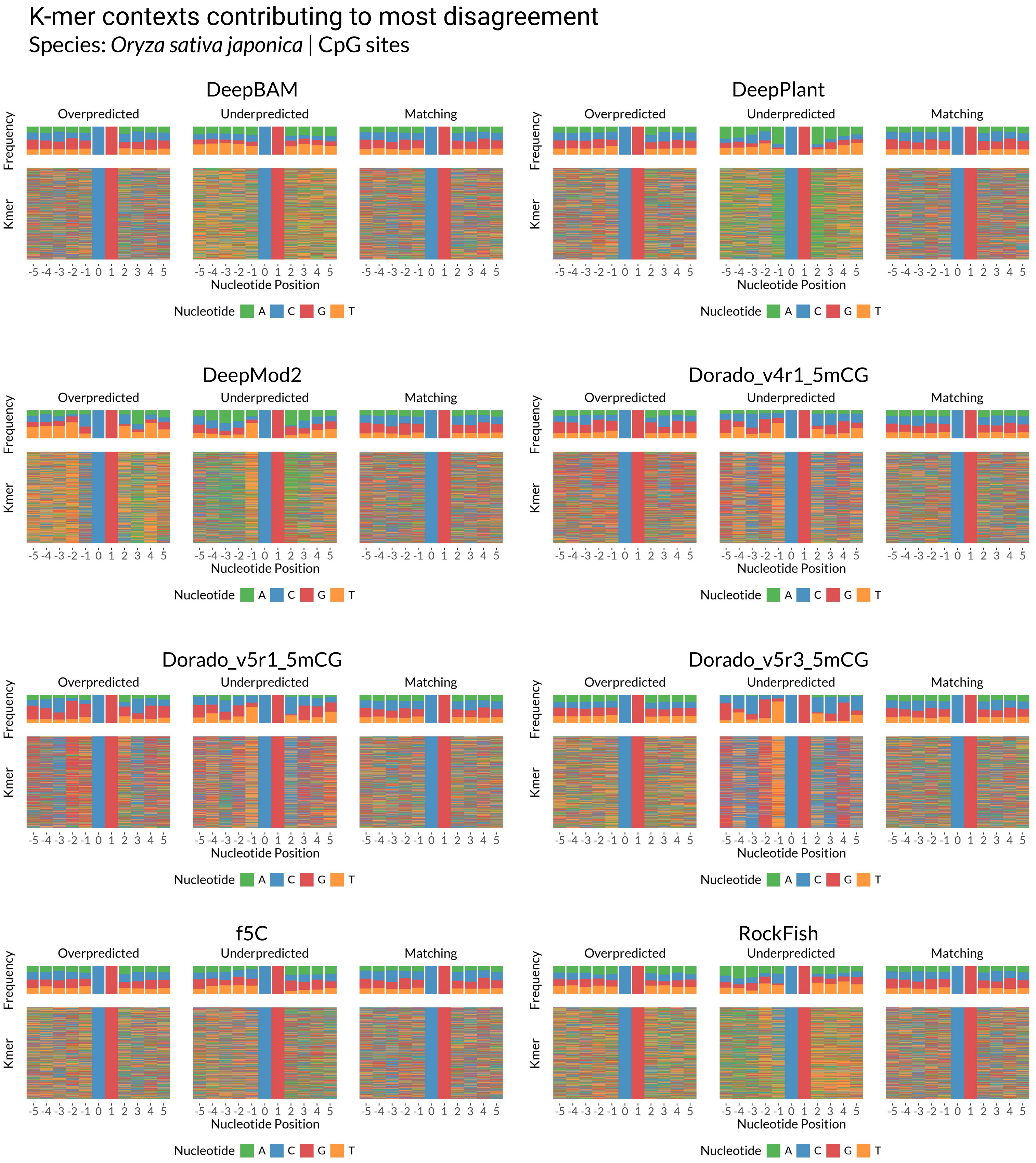


Fig S9: K-mers of disagreement at CpG locations on rice data. 5000 most disagreeing locations were plotted in each case; Overpredicted - the nanopore tool calls more methylation than ground-truth, Underpredicted - nanopore tool calls less methylation compared to ground-truth. Matching - 5000 randomly sampled locations where the difference of methylation level between the ground truth and reported value is 0.

### Fig S10

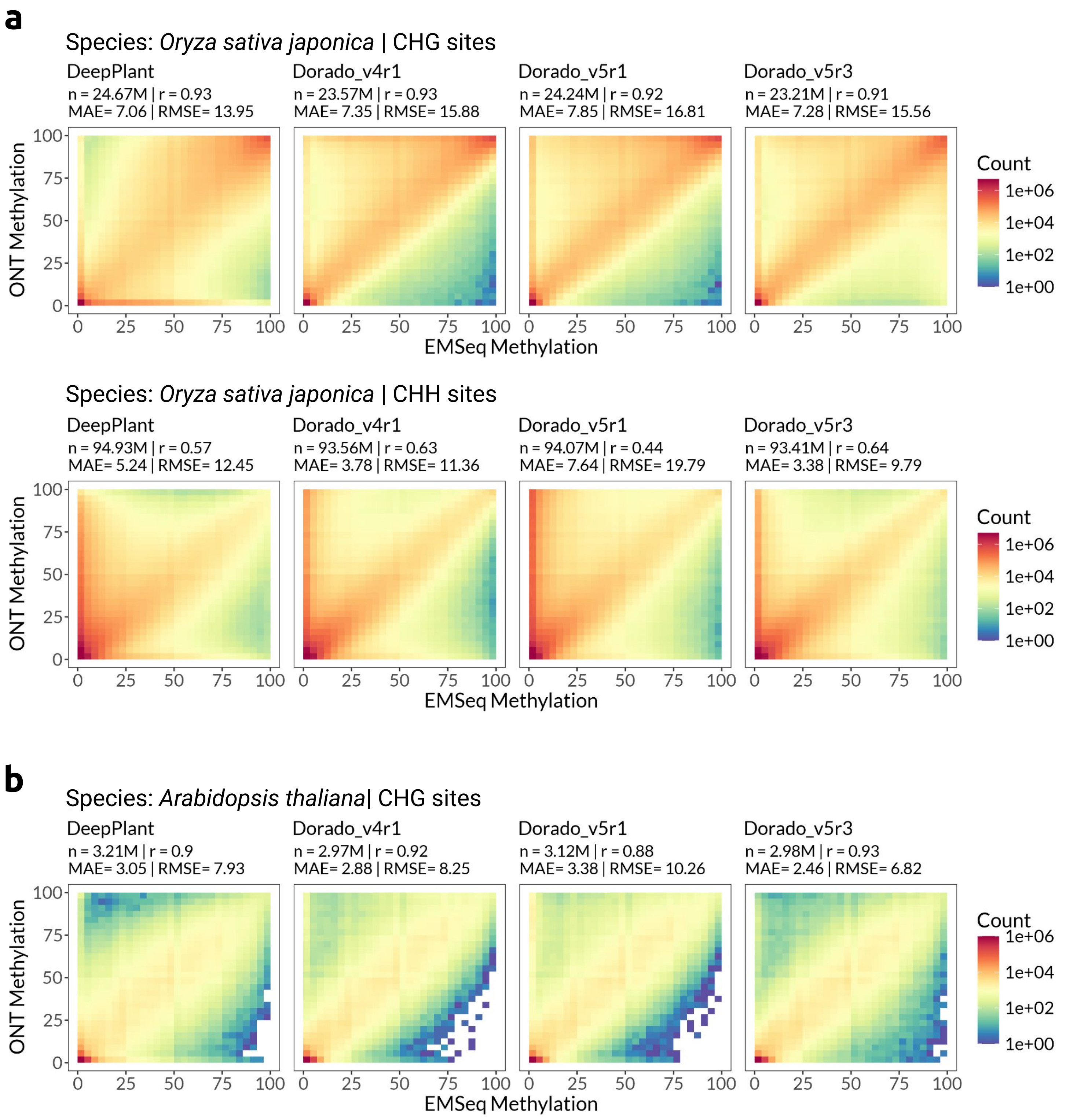


Fig S10: Site-level evaluation of methylation calling tools using correlation heatmaps in non-CG context, on plant data. n = number of sites profiled and covered by >=20 reads; r = Pearson correlation coefficient; MAE = Mean Absolute Error; RMSE = Root Mean-Squared Error. a) Correlation heatmap of CHG (top) and CHH (bottom) sites from *O.sativa* data for each tool compared to the EMSeq ground truth. b) Correlation heatmap of CHG sites from *A.thaliana* data for each tool compared to the EMSeq data ground truth.

### Fig S11

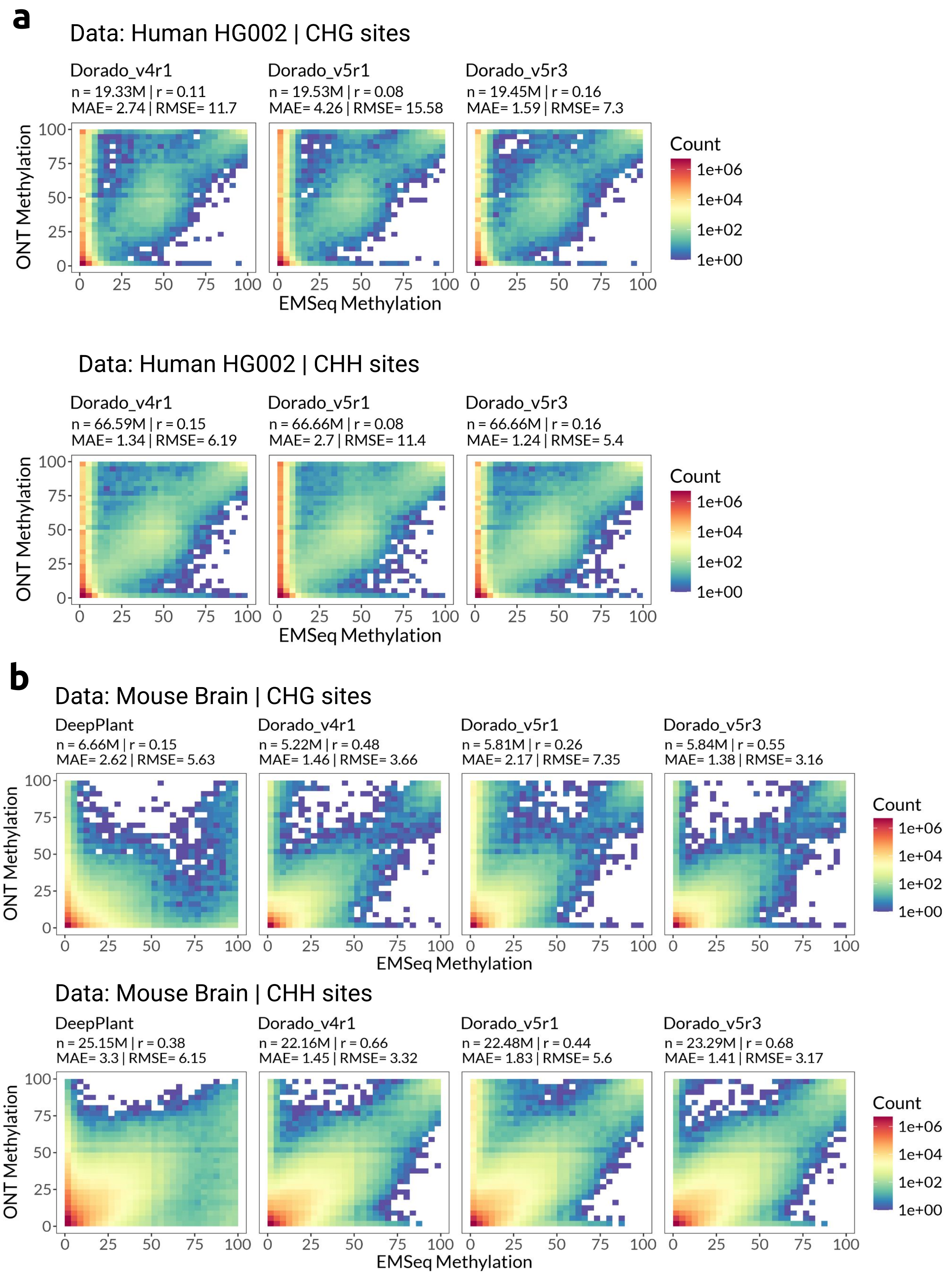


Fig S11: Site-level evaluation of non-CG methylation using correlation heatmaps, on human HG002 and mouse brain data. n = number of sites profiled and covered by >=20 reads; r = Pearson correlation coefficient; MAE = Mean Absolute Error; RMSE = Root Mean-Squared Error. a) Correlation heatmap of CHG (top) and CHH (bottom) sites from chromosome 1 of human data for the Dorado all-context 5mC models (a), chromosome 1 of mouse brain data for DeepPlant and Dorado all-context 5mC models (b) compared to the EMSeq ground truth.

### Fig S12

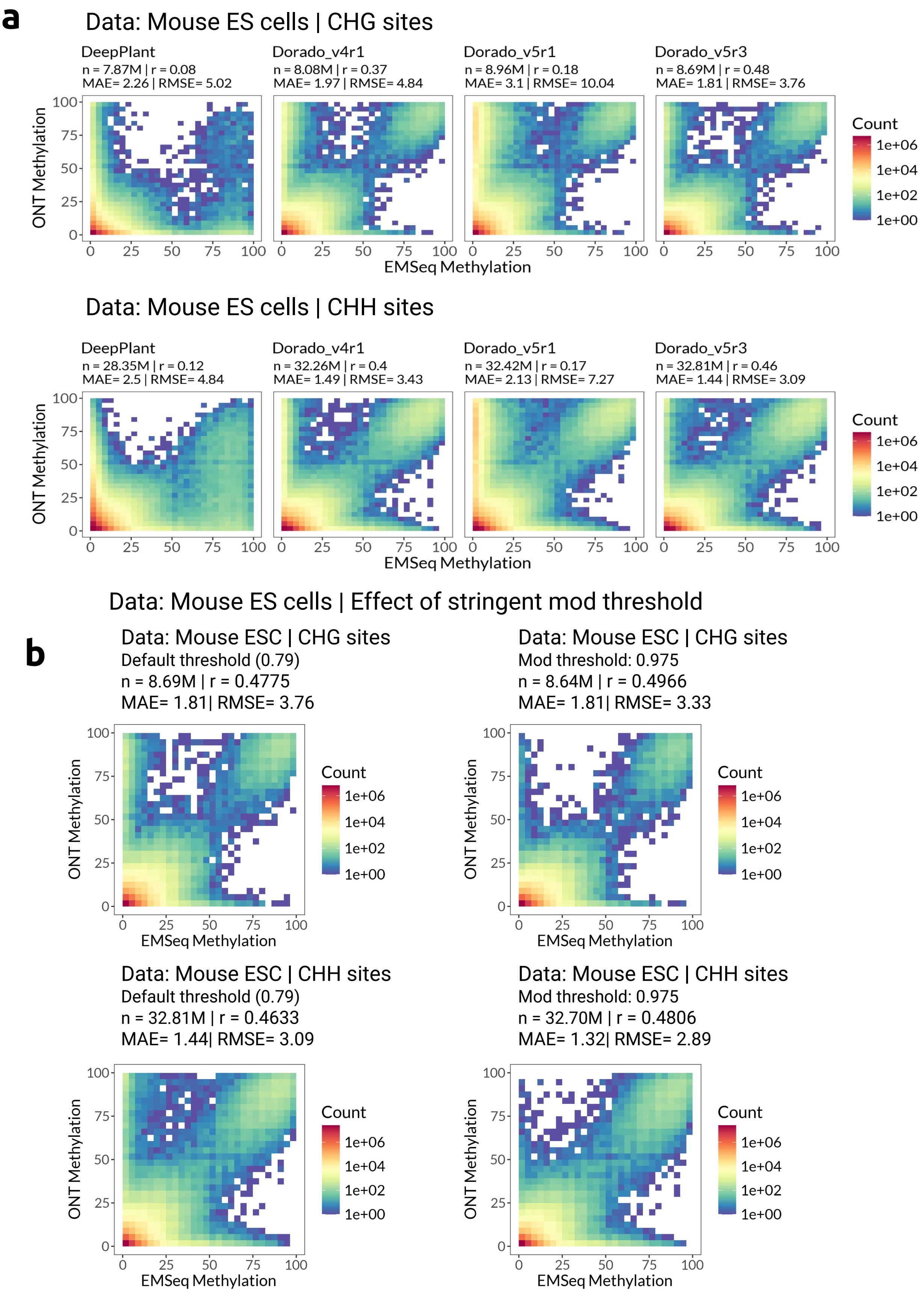


Fig S12: Site-level evaluation of non-CG methylation using correlation heatmaps on mouse ESC data, and effect of adjusting probability thresholds on false positives. n = number of sites profiled and covered by >=20 reads; r = Pearson correlation coefficient; MAE = Mean Absolute Error; RMSE = Root Mean-Squared Error. a-b) Correlation heatmap of CHG (top) and CHH (bottom) sites from mouse ESC chromosome 1 data for (a) each tool compared to the EMSeq ground truth. (b) Dorado v5r3 model with default threshold (0.79, left) and a stringent mod threshold (0.97, right) compared with the EMseq ground truth.

### Fig S13

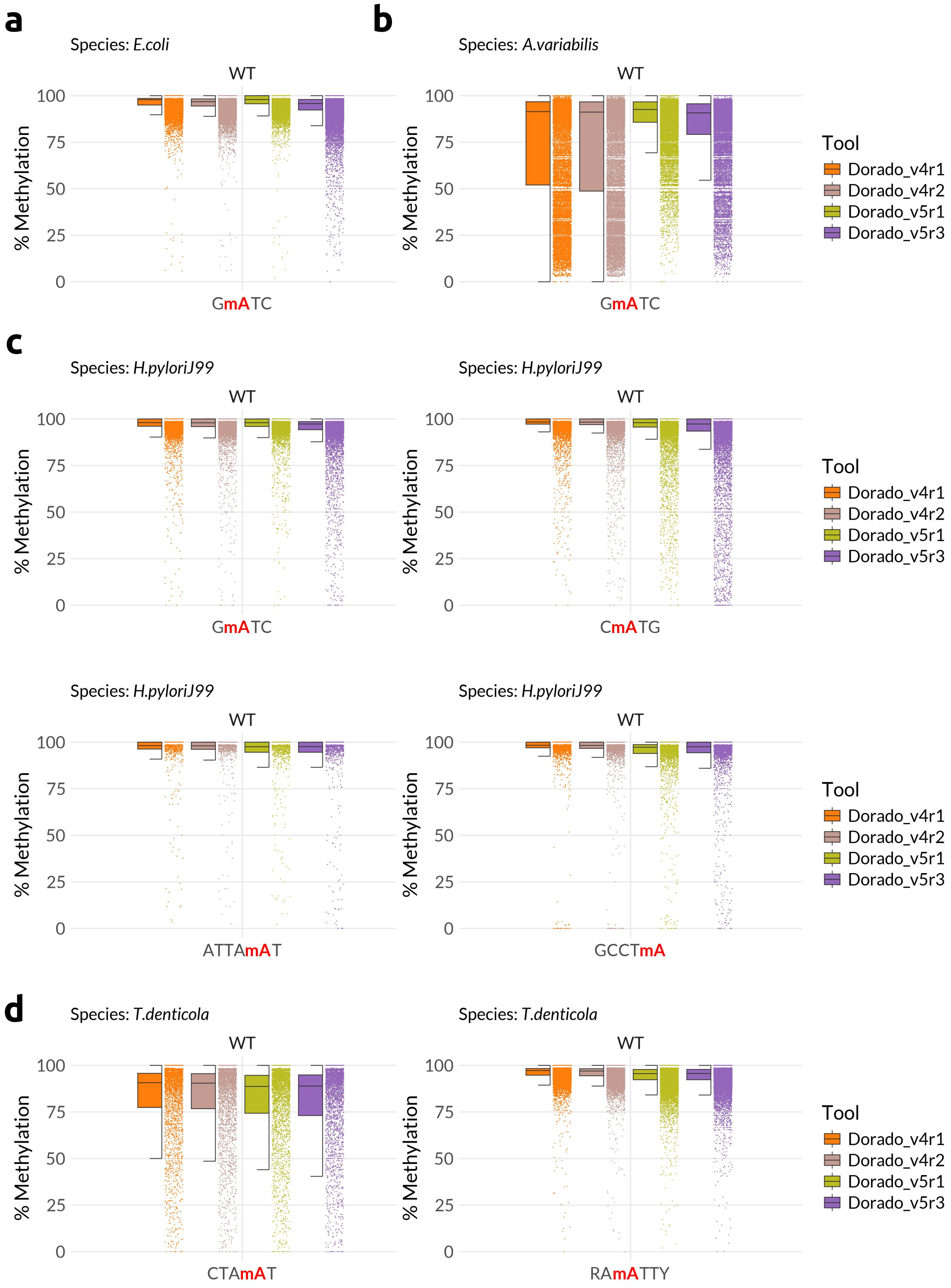


Fig S13: Context-wise evaluation of 6mA calling tools. a-d) Jitter-box plots depicting each tool's performance on GATC context in *E.coli* WT sample (a), GATC context in *A.variabilis* WT sample (b), GATC, CATG, ATTAAT, GCCTA contexts in *H.pylori* J99 WT sample (c) and CTAAT, RAATTY contexts in *T.denticola* WT sample (d).

### Fig S14

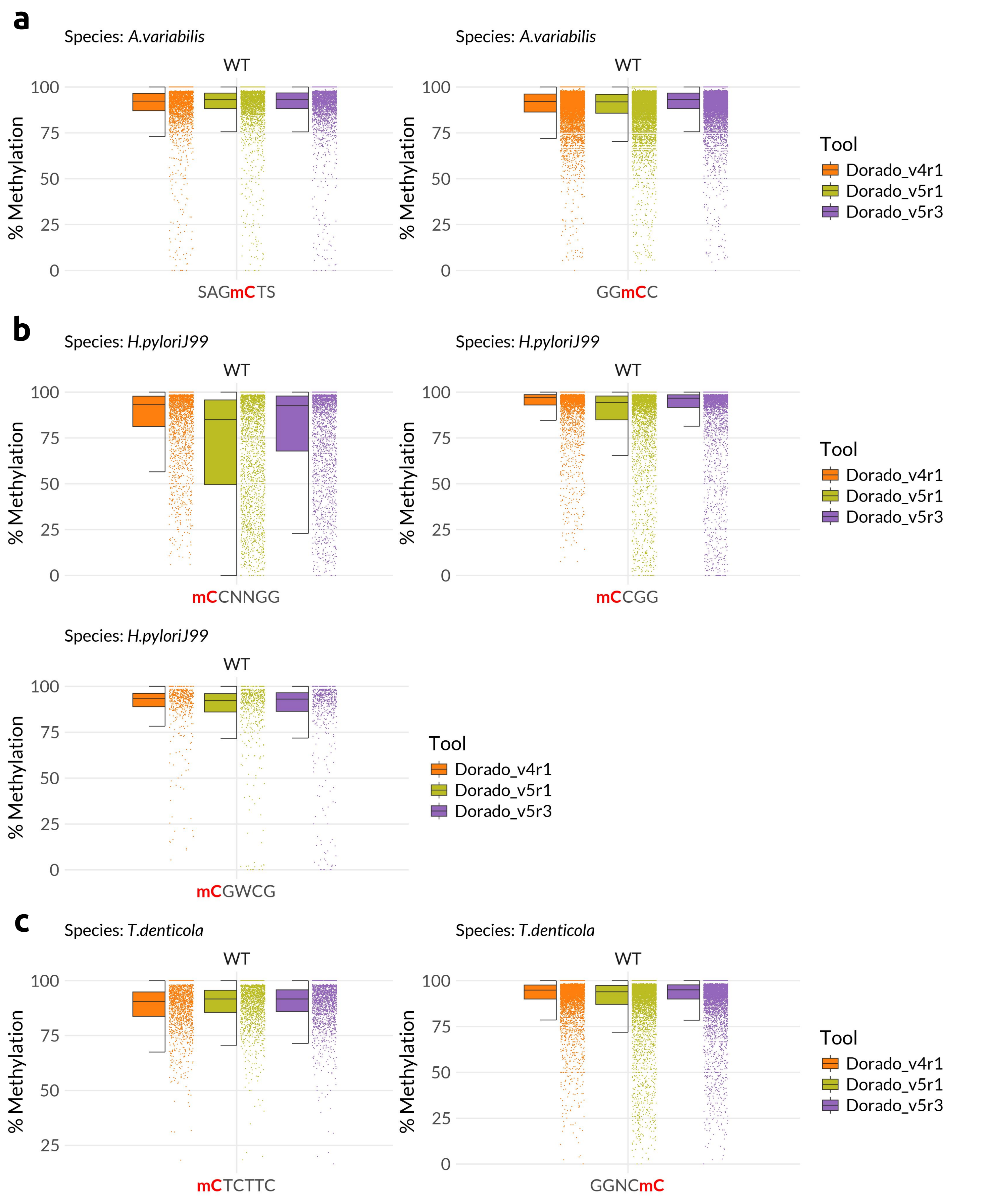


Fig S14: Context-wise evaluation of 4mC calling tools. a-c) Jitter-box plots depicting each tool's performance on SAGCTS, GGCC contexts in *A.variabilis* WT sample (a), CCNNGG, CCGG, CGWCG contexts in *H.pylori* J99 WT sample (b) and CTCTTC, GGNCC contexts in *T.denticola* WT sample (c).

### Fig S15

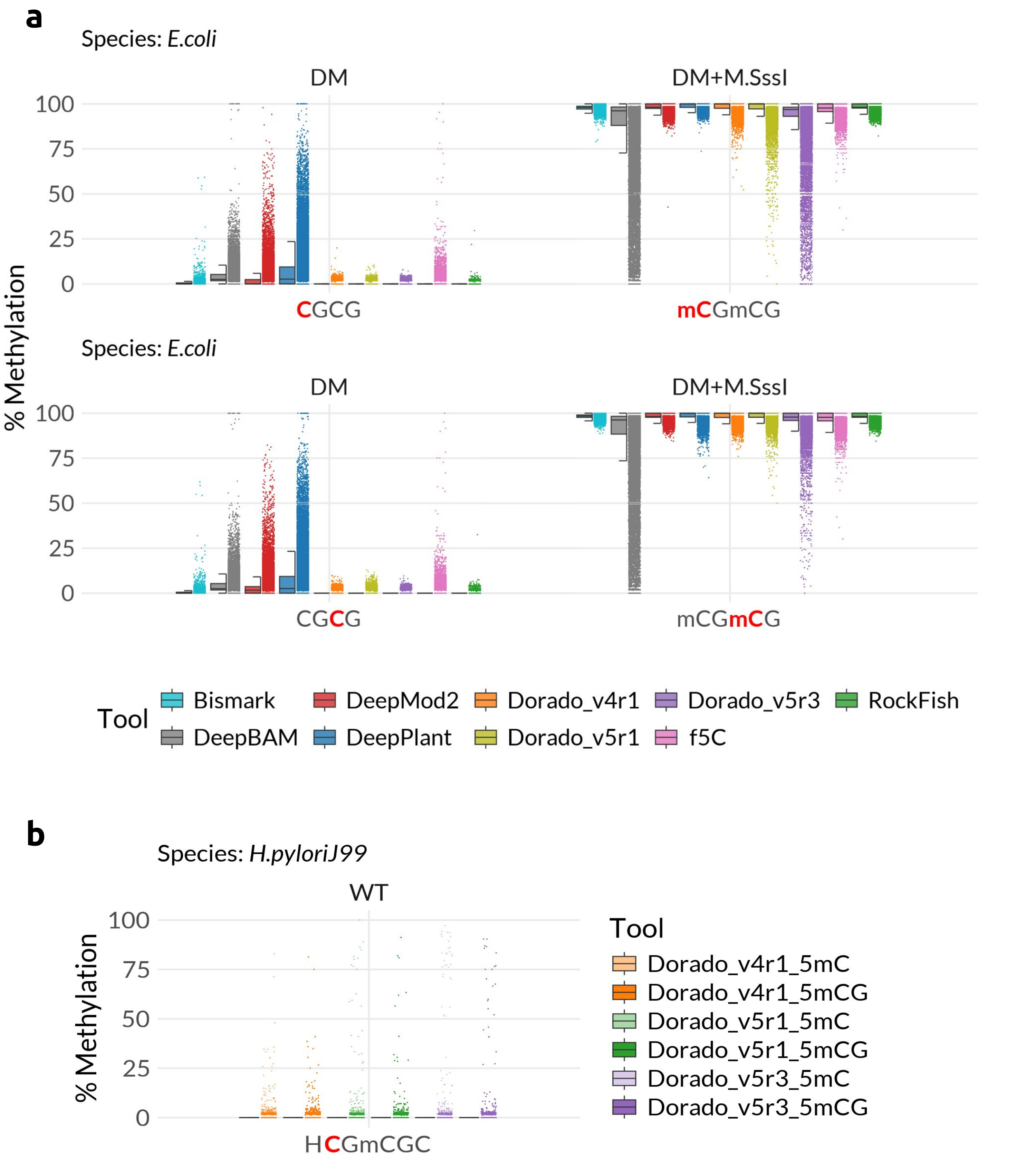


Fig S15: Evaluating the effect of neighboring modifications for 5mCG calling tools. The profiled C is shown in red and the methylation status is indicated by a preceding "m". a) Jitter-box plot depicting each tool's performance on the CGCG context in *E.coli* DM and DM+M.SssI samples. Both the consecutive CGs are expected to be methylated in the latter sample. b) Jitter-box plot depicting the performance of Dorado all-context 5mC and 5mCG models on HCGmCGC context in *H.pylori* J99 WT sample. The profiled C, depicted in red is part of a CG motif preceding an mCG.

### Fig S16

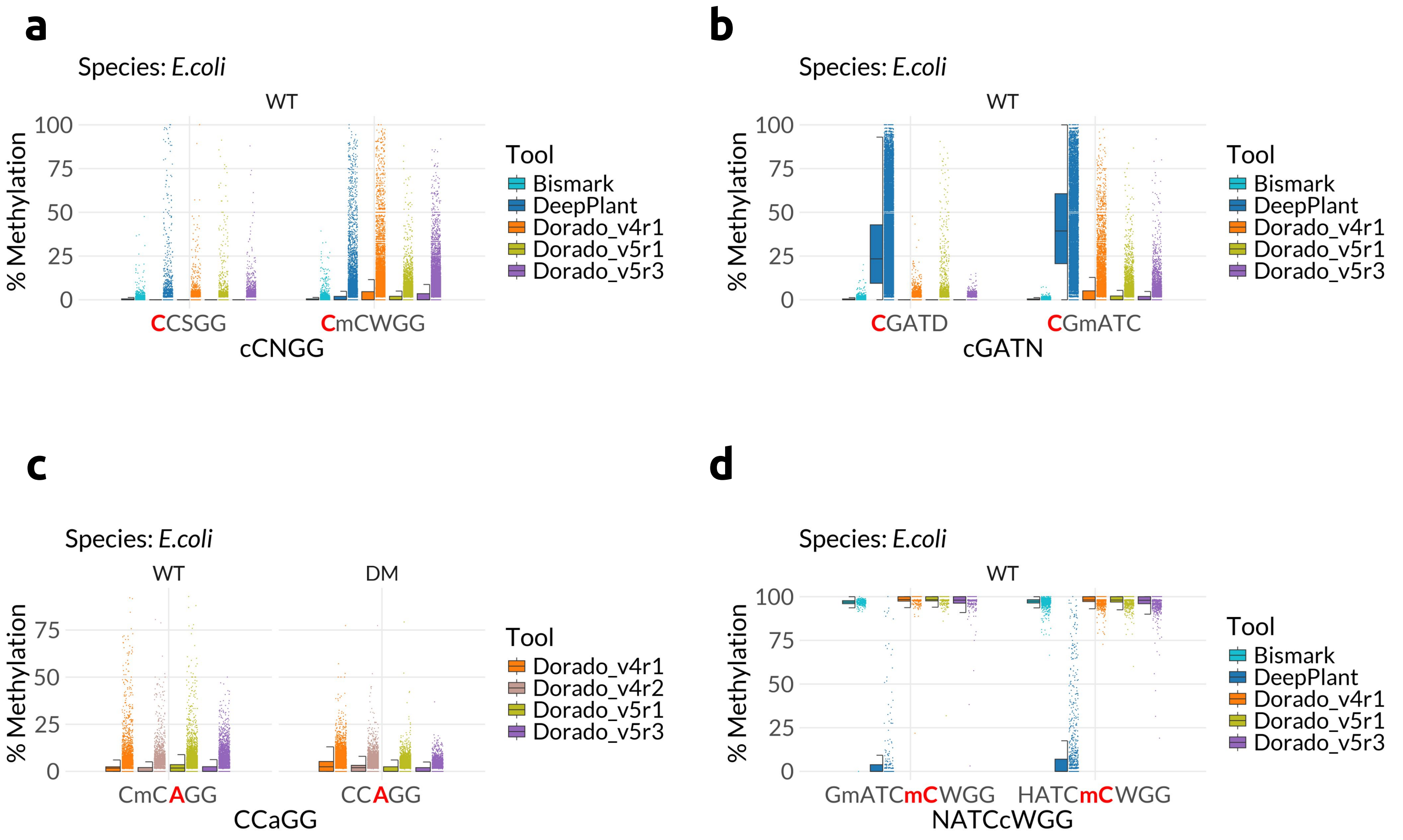


Fig S16: Evaluating the effect of neighboring modifications in various contexts. Profiled base is shown in red. The methylation status of the bases is indicated with a preceding “m”. a) Jitter-box plot depicting effect of mC on umC for each tool in CCNGG context (CCSGG,unmethylated; CCWGG,methylated) in *E.coli* WT sample. b) Jitter-box plot depicting effect of mC on umC for each tool in cGATN context (CGATD,unmethylated; CGATC,methylated) in *E.coli* WT sample. c) Jitter-box plot depicting effect of mC on umA for each tool in CCAGG context in *E.coli* DM (unmethylated) and WT (methylated) samples. d) Jitter-box plot depicting effect of mA on mC for each tool in NATCCWGG context (HATCCWGG,unmethylated; GATCCWGG,methylated) in *E.coli* WT sample.

Fig S17

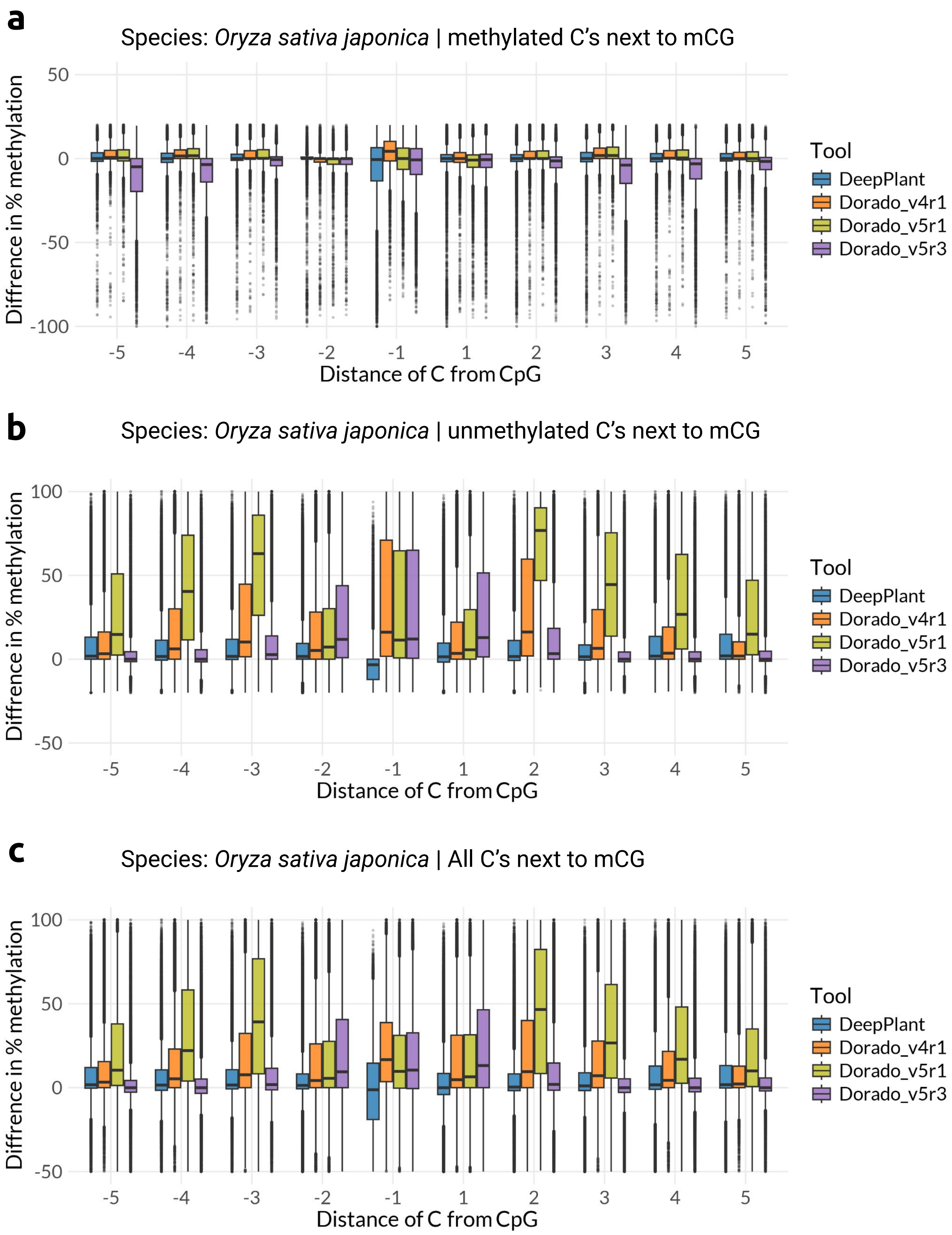


Fig S17: Evaluating the effect of neighboring CG methylation on rice data. The difference in methylation calls for a cytosine as a function of its distance from methylated CpG. Negative values indicate underestimation of methylation near mCG. EMSeq data used as ground truth. a-c) Boxplot showing difference in methylation calls from the tool and EMSeq data for sites where profiled C is methylated (>=80% methylation) (a), profiled C is unmethylated (<=20% methylation) (b) and for all Cs neighboring mCG (c) in *O.sativa*.

Fig S18

**a**

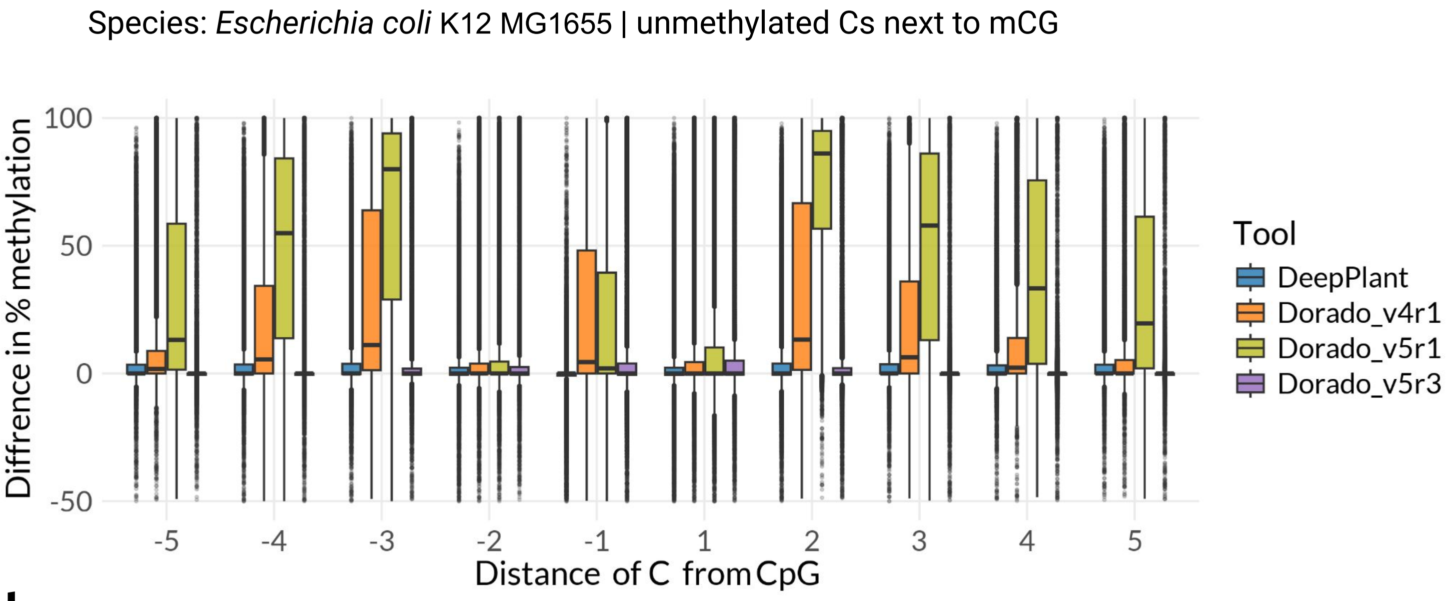


**b**

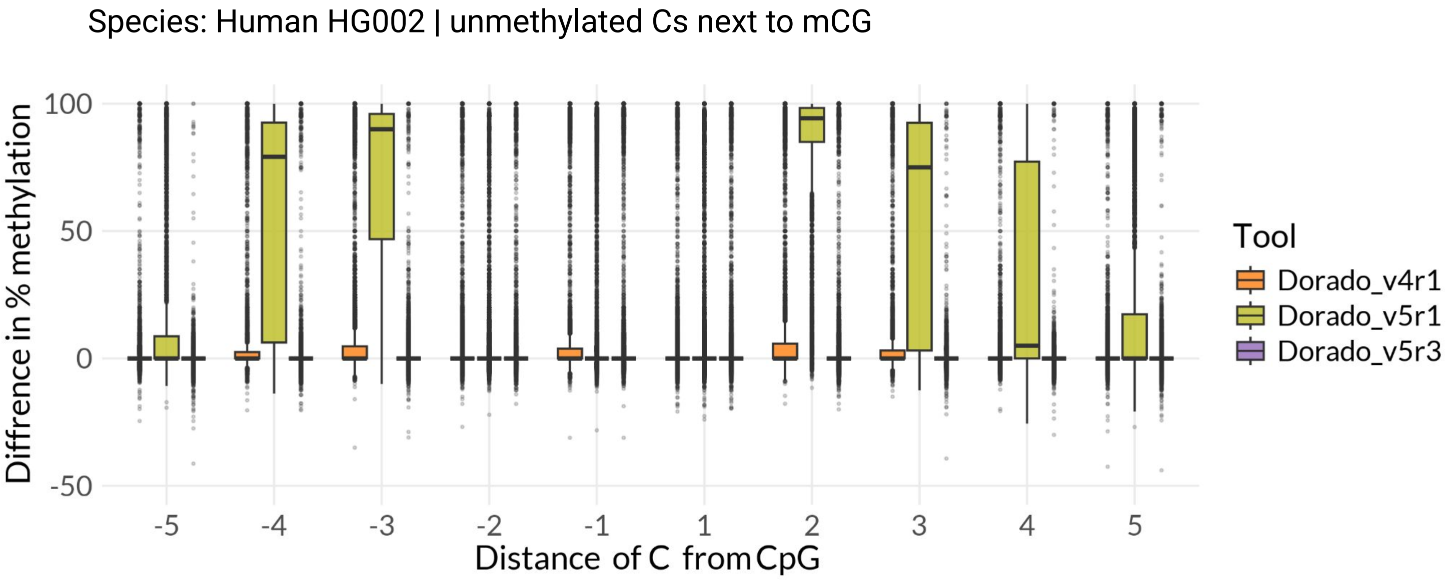


Fig S18: Evaluating the effect of neighboring CG methylation on *E.coli* and human data. The difference in methylation calls for a cytosine as a function of its distance from methylated CpG. Negative values indicate underestimation of methylation near mCG. EMSeq data is used as ground truth. a-b) Boxplots showing difference in methylation calls from the tool and EMSeq data for unmethylated Cs neighboring mCG in *E.coli* (a), unmethylated Cs neighboring mCG in human data (b).

Fig S19

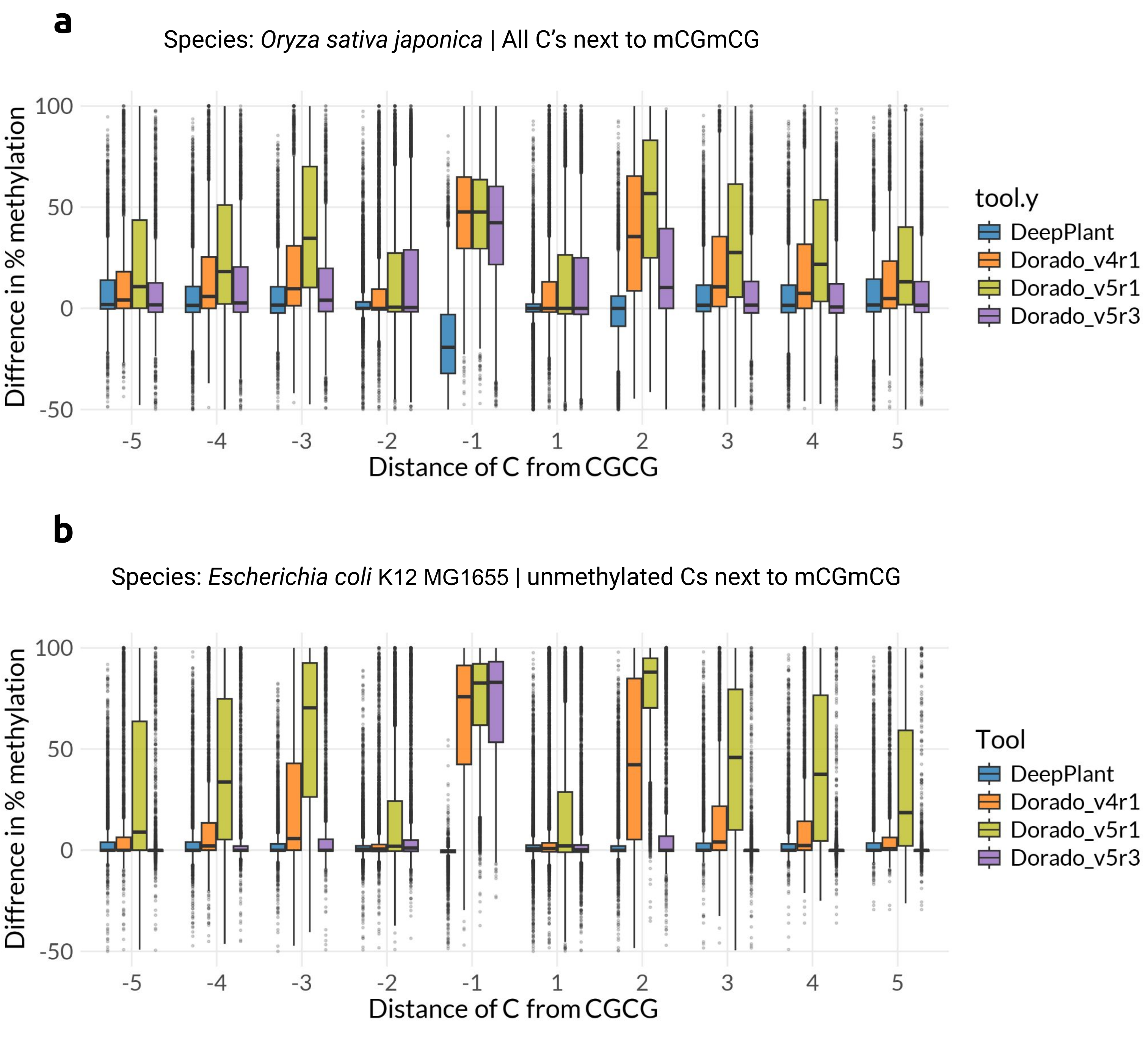


Fig S19: Effect of two neighboring methylated CpGs. The difference in methylation calls for a cytosine as a function of its distance from CGCG sites where both CGs are methylated (mCGmCG). Negative values indicate underestimation of methylation near mCGmCG. EMSeq data is used as ground truth. a-b) Boxplots showing difference in methylation calls from the tool and EMSeq data for all Cs neighboring mCGmCG in *O.sativa* (a), unmethylated Cs neighboring mCGmCG in *E.coli* (b).

### Fig S20

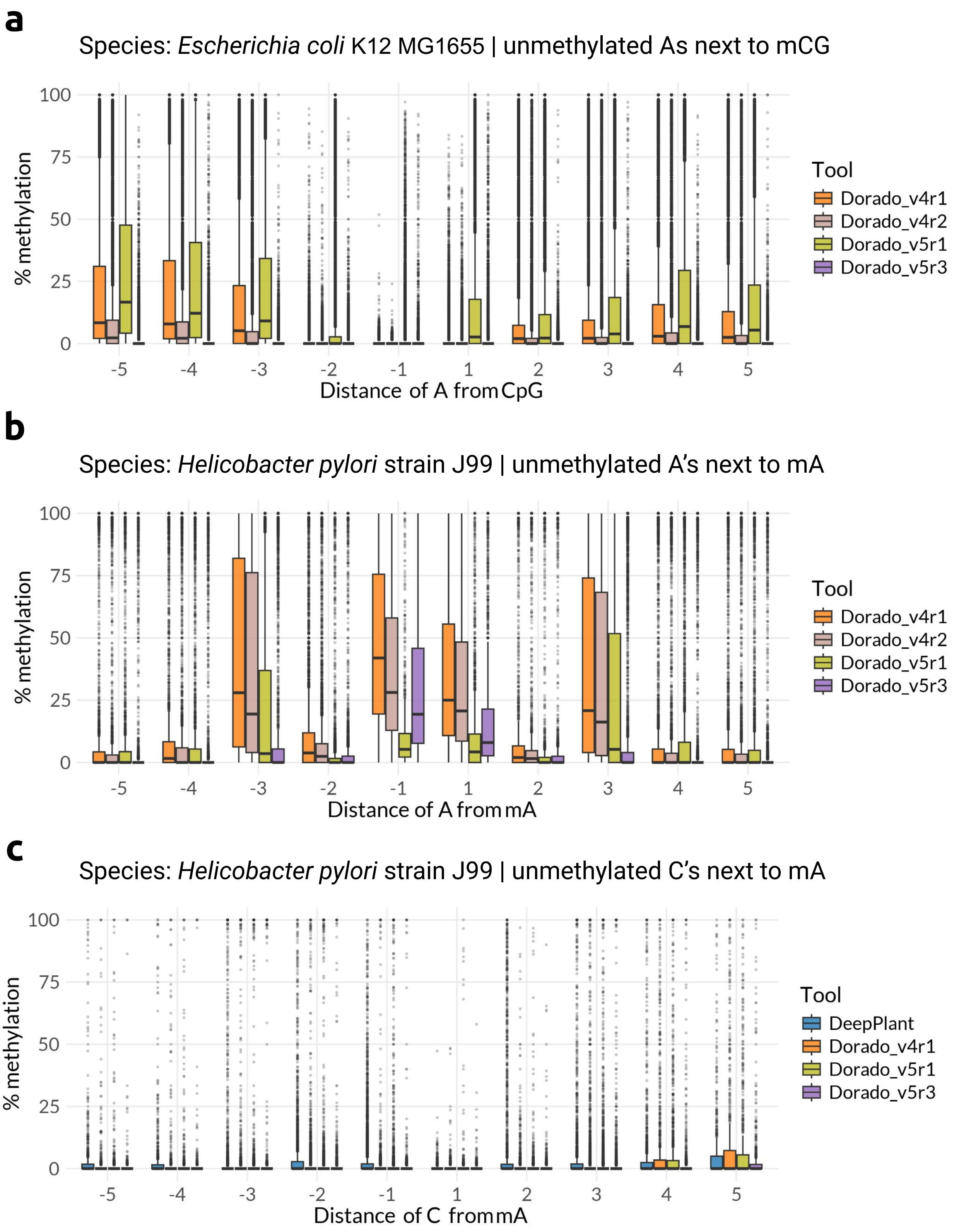


Fig S20: Evaluating the effect of neighboring modifications on adenine methylation calling. Methylation percentage for a cytosine as a function of its distance from methylated sites. a-c) Boxplots showing methylation percentage from each tool for unmethylated As neighboring mCG sites in *E.coli* (a), unmethylated As neighboring mA sites in *E.coli* (b), unmethylated Cs neighboring mA sites in *H.pylori* J99 (c).

### Fig S21

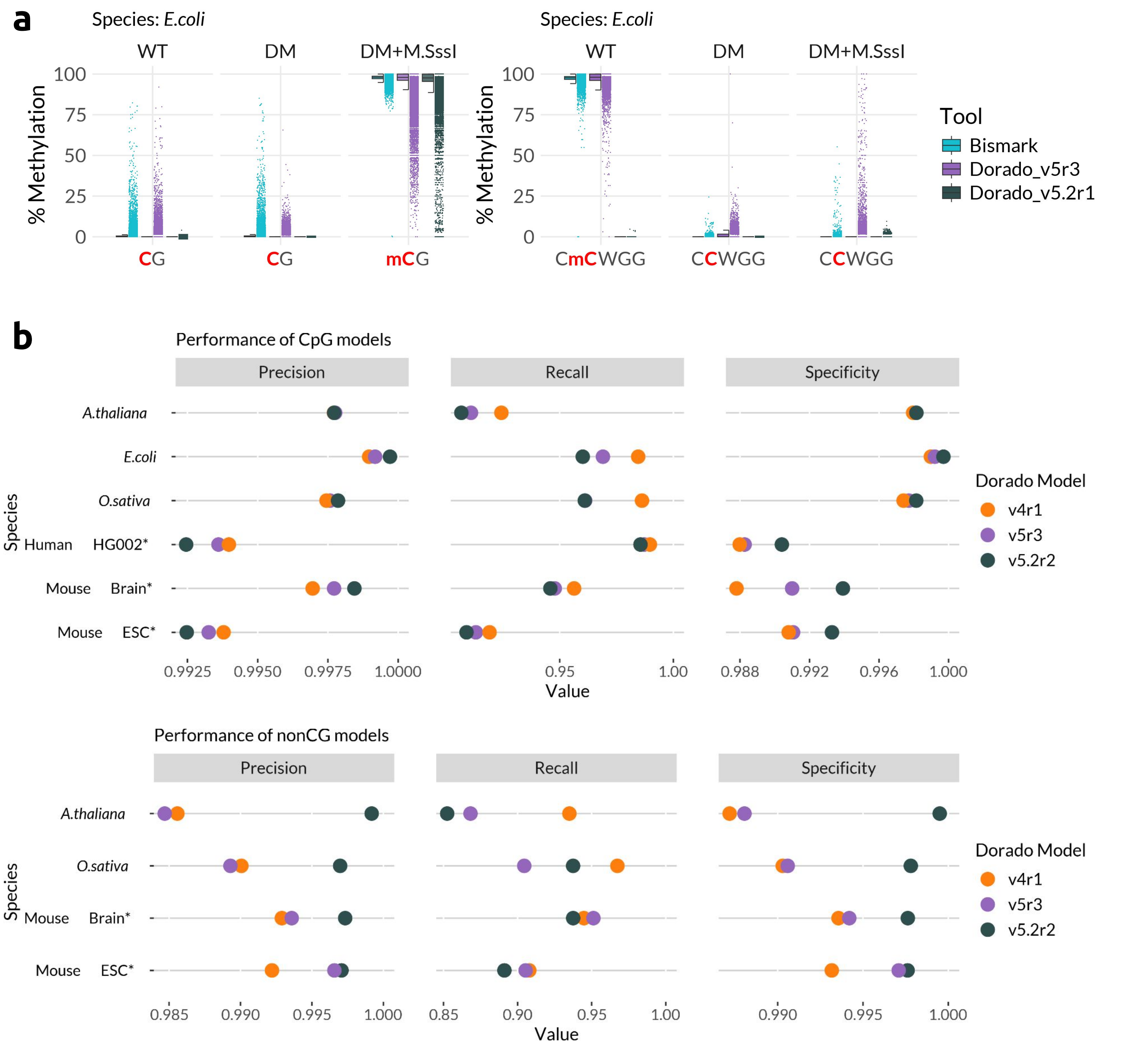


Fig S21: Analysis of the new v5.2 models of Dorado in CpG and non-CG contexts. a) Performance of v5.2r1 models. Jitter-box plots depicting the percentage methylation in CG and CCWGG motifs in *E.coli* DM, WT and DM+M.SssI datasets. The profiled base is highlighted in red font. The methylation status of the bases is indicated with a preceding "m". The all context v5.2r1 model completely failed to report non CG methylation. b) Read level evaluation of v5.2r2 CpG (up) and all-context C (down) models in comparison to v4r1 and v5r3 models. Precision (left), Recall (middle), Specificity (right) scores are depicted as colored dots on a number line (x-axis). Each line represents a dataset. The issue that was seen in v5.2r1 model has been rectified in the r2 models.

### Fig S22

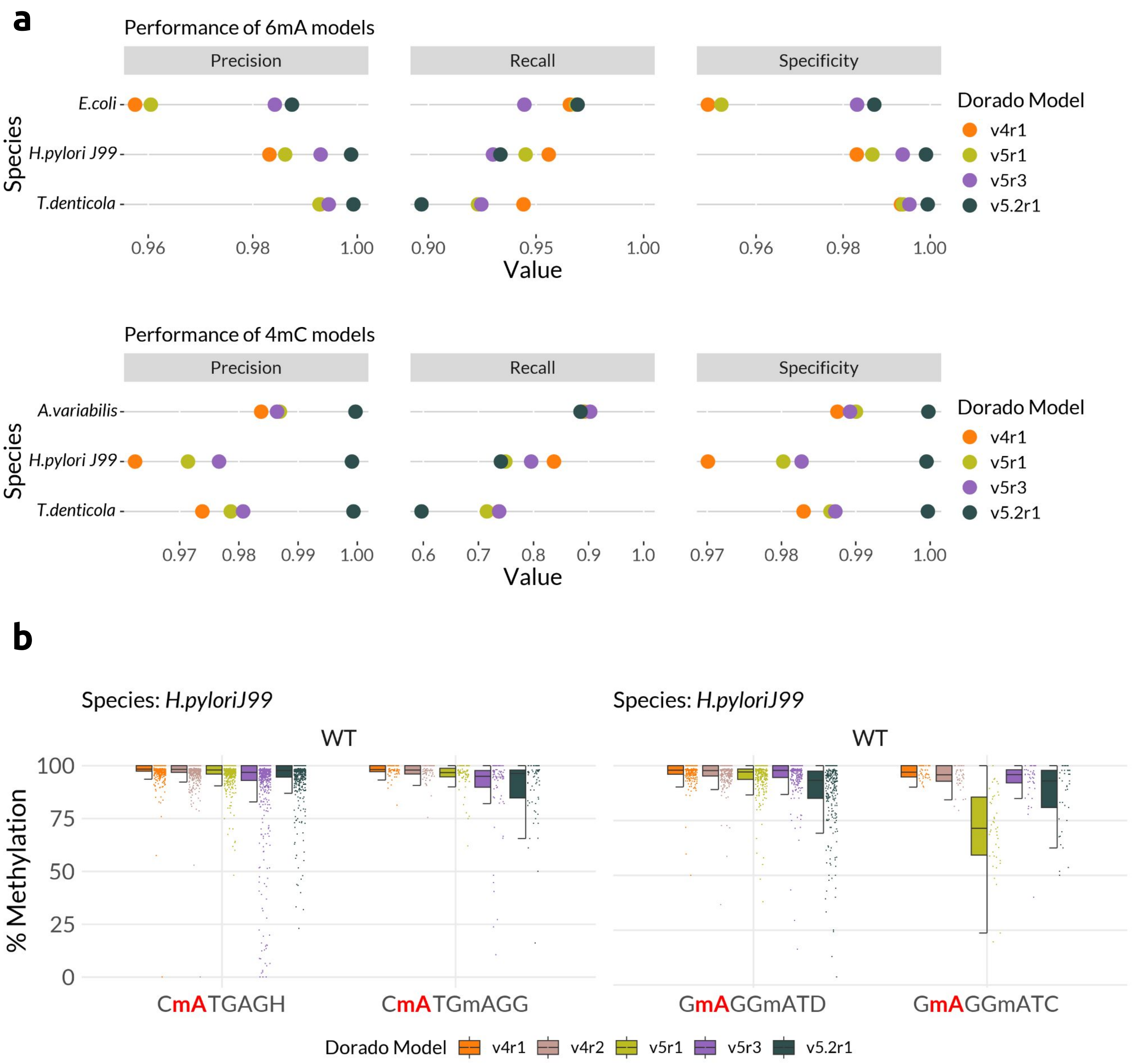


Fig S22: Analysis of the new v5.2 6mA and 4mC models of Dorado. a) Read level evaluation of v5.2r1 6mA (up) and 4mC (down) models in comparison to v4r1, v5r1 and v5r3 models. Precision (left), Recall (middle), Specificity (right) scores are depicted as colored dots on a number line (x-axis). Each line represents a dataset. b) Jitter-boxplots depicting the percent methylation of profiled As in the contexts CaTGAGN and GaGGATN, in *H.pylori* str J99. In each context the mA preceding an umA is profiled, in contrast to profiling an mA preceding an mA.

### Fig S23

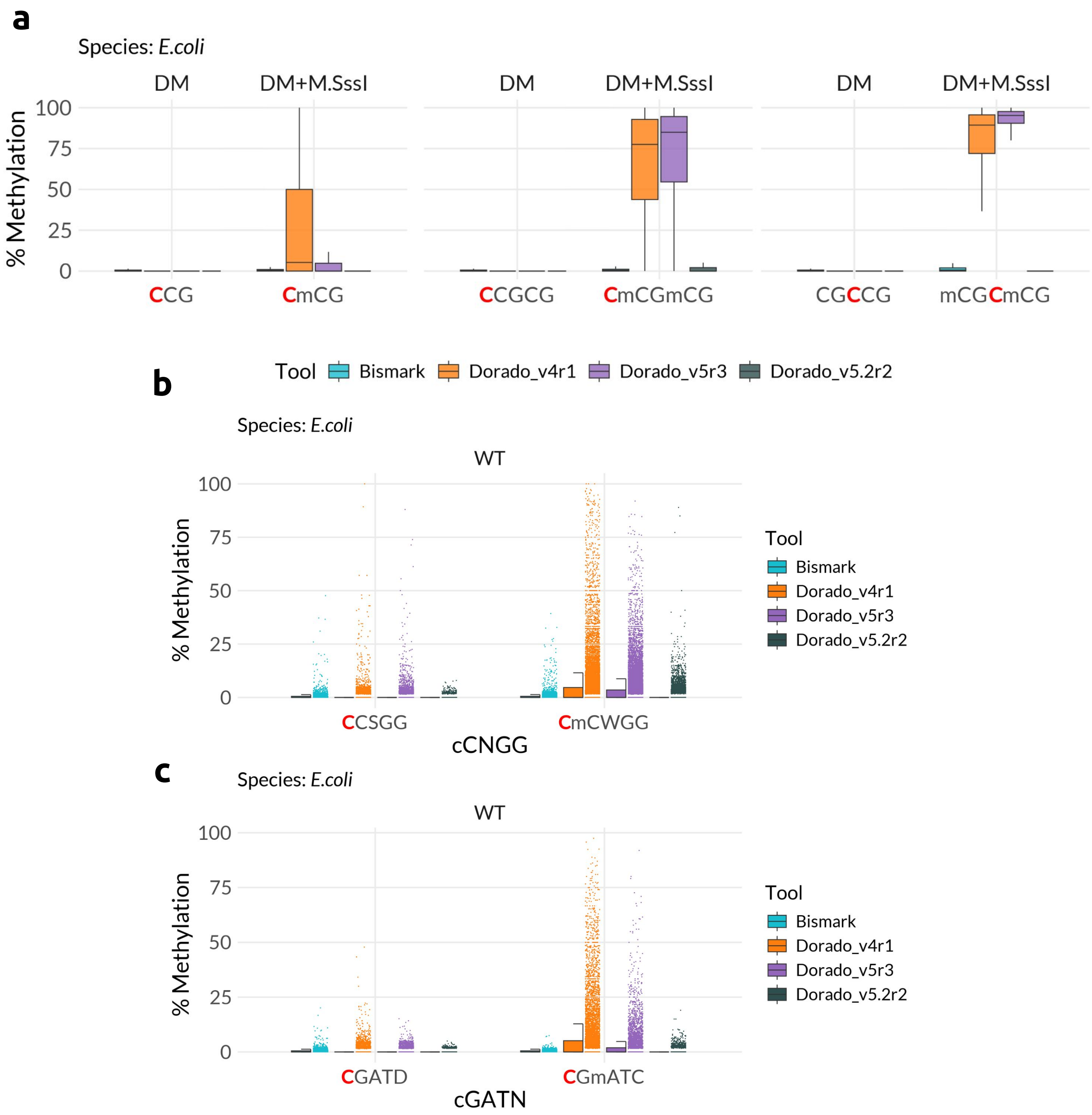


Fig S23: Evaluating the effect of neighboring modifications on performance of v5.2 models, in comparison to v4r1 and v5r3 models in various contexts, using the *E.coli* dataset. Profiled base is shown in red. The methylation status of the bases is indicated with a preceding "m". a) Box plot depicting effect of mC on umC in various contexts in *E.coli* MSssI sample. b) Jitter-box plot depicting effect of mC on umC in CCNGG context (CCSGG, unmethylated; CCWGG, methylated) in *E.coli* WT sample. c) Jitter-box plot depicting effect of mA on umC in cGATN context (CGATD, unmethylated; CGATC, methylated) in *E.coli* WT sample.

Fig S24

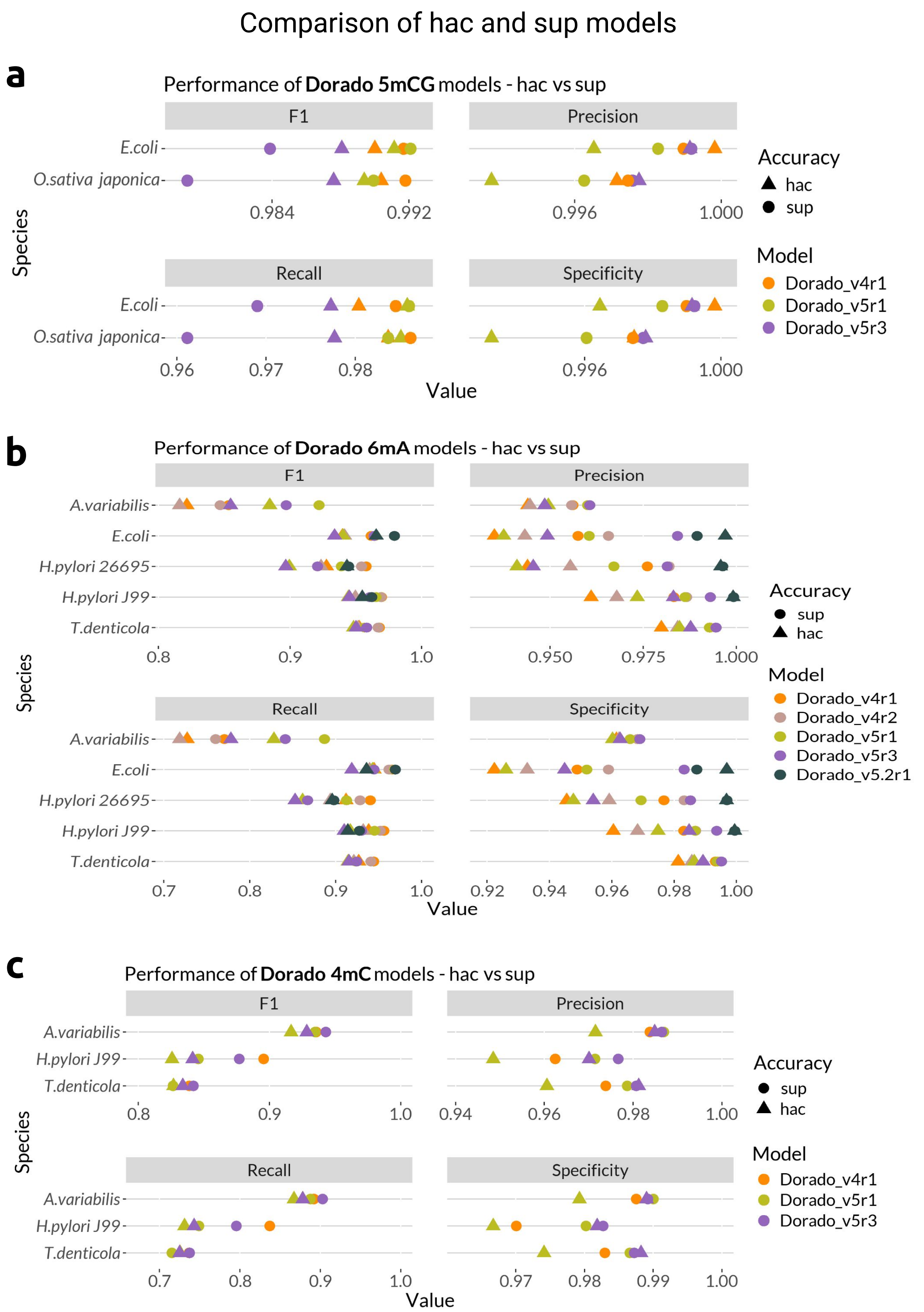


Fig S24: Read-level comparison of Dorado 5mCG, 6mA and 4mC high-accuracy (hac) and super-accuracy (sup) models. The performance metrics, F1, Precision, Recall, and Specificity are plotted on a number line (x-axis) colored by tools. Each line represents a dataset. The sup models are plotted as triangles and the hac models as dots. a) The performance metrics for the Dorado 5mCG models of the rice and *E.coli* data are plotted on a number line. b) The performance metrics for both Dorado 6mA sup and hac models are compared across all bacterial species. c) The performance metrics for Dorado 4mC models for *A.variabilis*, *H.pylori* J99 and *T.denticola* data are plotted on a number line.

### Fig S25

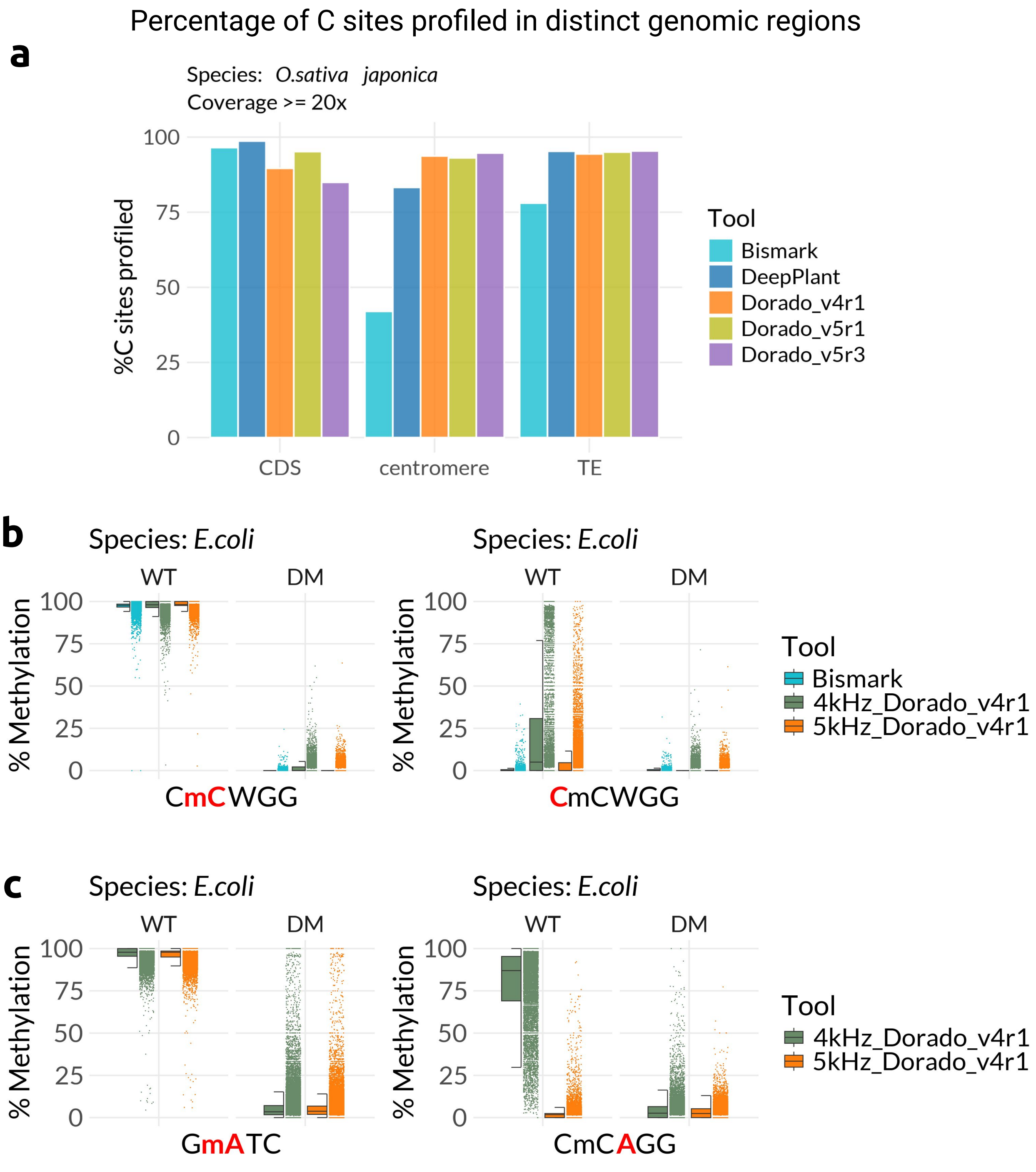


Fig S25: Sites profiled by EMSeq and Nanopore, and susceptibility of 4kHz and 5Hz models to neighboring methylation. a) Percentage of cytosines in all contexts profiled (and covered by >=20 reads) in coding sequences, centromeres and transposable elements of the rice genome by different tools. The percentage of Cs profiled (y-axis) from the total genome at each distinct genomic region (x-axis) are shown as bar plots, colored by the methylation calling tools. b-c) Performance of Dorado 4kHz and 5kHz v4 models compared to the ground truth EMSeq data in the CCWGG (b) and GATC contexts (c) profiled in *E.coli* WT (methylated) and DM (unmethylated) samples (left). Effect of neighboring mC on unmethylated C (b), neighboring mC on unmethylated A (c) in CCWGG context in *E.coli* WT (methylated) and DM (unmethylated) samples (right).
